# ADAR1p150 Prevents MDA5 and PKR Activation via Distinct Mechanisms to Avert Fatal Autoinflammation

**DOI:** 10.1101/2023.01.25.525475

**Authors:** Shi-Bin Hu, Jacki Heraud-Farlow, Tao Sun, Zhen Liang, Ankita Goradia, Scott Taylor, Carl R Walkley, Jin Billy Li

**Affiliations:** Department of Genetics, Stanford University, Stanford, CA, 94305, USA; St Vincent’s Institute of Medical Research, Fitzroy, Victoria, 3065, Australia; Department of Medicine, Eastern Hill Precinct, Melbourne Medical School, University of Melbourne, Fitzroy, Victoria, 3065, Australia

**Keywords:** ADAR1, RNA editing, PKR, MDA5, dsRNA, innate immunity

## Abstract

Effective immunity requires the innate immune system to distinguish foreign (non-self) nucleic acids from cellular (self) nucleic acids. Cellular double-stranded RNAs (dsRNAs) are edited by the RNA editing enzyme ADAR1 to prevent their dsRNA structure pattern being recognized as viral dsRNA by cytoplasmic dsRNA sensors including MDA5, PKR and ZBP1. A loss of ADAR1-mediated RNA editing of cellular dsRNA activates MDA5. However, additional RNA editing-independent functions of ADAR1 have been proposed, but a specific mechanism has not been delineated. We now demonstrate that the loss of ADAR1-mediated RNA editing specifically activates MDA5, while loss of the cytoplasmic ADAR1p150 isoform or its dsRNA binding activity enabled PKR activation. Deleting both MDA5 and PKR resulted in complete rescue of the embryonic lethality of *Adar1p150*^-/-^ mice to adulthood, contrasting with the limited or no rescue by removing MDA5, PKR or ZBP1 alone, demonstrating that this is a species conserved function of ADAR1p150. Our findings demonstrate that MDA5 and PKR are the primary *in vivo* effectors of fatal autoinflammation following the loss of ADAR1p150.

## Introduction

Adenosine-to-inosine (A-to-I) RNA editing, catalyzed by the adenosine deaminase acting on RNA (ADAR) family of enzymes, is one of the most prevalent RNA modifications in metazoans^1–5^. The ADAR enzymes bind to regions of double-stranded RNA (dsRNA) and convert targeted adenosines into inosine. There are two catalytically active ADAR enzymes in mammals, ADAR1 and ADAR2. Unlike ADAR2, which is highly expressed in the central nervous system, ADAR1 is widely expressed across tissues. ADAR1 uniquely has two isoforms: the p110 isoform is constitutively expressed and localizes to the nucleus, while the p150 isoform is primarily cytoplasmic and can be induced by interferon (IFN)^6,7^.

Cytosolic dsRNAs, both endogenous/cellular derived and viral, can trigger an innate immune response upon recognition by cytosolic RNA-sensing receptors including MDA5, RIG-I, PKR, OAS-RNaseL, and ZBP1^8^. Upon dsRNA recognition, MDA5 and RIG-I activate type I interferon (IFN) signaling via MAVS, TBK1, and IRFs to induce the expression of IFNα/β and IFN-stimulated genes (ISGs)^9,10^. PKR and OAS-RNaseL exert direct antiviral activity through translational shutdown^11^ and RNA cleavage^12^, respectively. ZBP1, together with ADAR1p150, are the only mammalian Zα domain containing proteins and bind to Z-form nucleic acids^13–16^. ZBP1 engages dsRNA triggering inflammation and cell death^17,18^. While the RNA-sensing system effectively detects cytosolic dsRNAs, how does it discriminate “self” dsRNAs from “non-self”? Human and mouse genetics studies have revealed that ADAR1-mediated A-to-I RNA editing is a critical mechanism^19–22^. In addition to contributing to the diversity of the transcriptome^1,23^, RNA editing by ADAR1 marks endogenous dsRNAs as “self” molecules^19–22,24^. In humans, ADAR1 loss-of-function mutations^22,25^ and MDA5 gain-of-function mutations^26,27^ have been associated with elevated type I IFN signaling in rare autoinflammatory diseases, such as Aicardi Goutières Syndrome (AGS). In common inflammatory diseases, genetic risk variants that collectively reduce the editing level of nearby dsRNAs are associated with elevated IFN responses^28^.

*Adar1*^-/-^ and *Adar1p150*^-/-^ mice die *in utero* at embryonic day (E) ~11.5-12.0^29–31^. *Adar1^E861A/E861A^* mice, with a catalytically inactive ADAR1 protein that is incapable of RNA editing, largely phenocopy the *Adar1^-/-^* mice dying *in utero* at E13.5. This demonstrates an essential requirement for the RNA editing function of ADAR1^19^. Concurrent MDA5 deletion (*Ifih1*^-/-^) rescued the *Adar1^E861A/E861A^* mice to a full life span^19,32^. Therefore ADAR1-mediated A-to-I RNA editing marks endogenous cellular dsRNAs as “self”, suppressing their recognition as “non-self” by MDA5 and preventing the subsequent innate immune response. This has been confirmed in human cells lines. However, removal of MDA5 or its downstream effector MAVS rescued the embryonic lethality of the *Adar1*^-/-^ mice (lacking the ADAR1 protein) to only ~2 days of age^20,21^, indicating additional protein-dependent, but RNA editing-independent, functions of ADAR1.

Long preceding the delineation of the ADAR1-dsRNA-MDA5 axis, a leading hypothesis to account for the embryonic lethality of *Adar1*^-/-^ mice was the activation of PKR. However, PKR loss (*Eif2ak2*^-/-^) did not rescue the embryonic lethality of *Adar1*^-/-^ mice^30^. In contrast to the *in vivo* observation, ADAR1 has been demonstrated to inhibit PKR activation in various cell lines^24,33–43^. This has been attributed to both RNA-editing and RNA-binding functions of ADAR1, however, there is some disparity between studies. Multiple studies have identified a subset of cancer cells that are sensitive to loss of ADAR1^33,38,44^. Interestingly, this dependency can be rescued *in vitro* by removal of PKR but not MDA5^33,38,44^. These observations reveal discrepancies between mouse models and cultured human cell line data: in mouse models, deletion of MDA5 or MAVS, but not PKR, rescued the lethality of ADAR1 loss, while in the cultured human cell lines, PKR, but not MDA5, had a significant contribution. Whether this incongruity is due to altered requirements for different sensors between cell types/species or due to the *in vivo* tissue environment and extrinsic signals has not been determined. Recently, removal of ZBP1 or its downstream effectors was reported to extend the survival of *Adar1*^-/-^*Mavs*^-/-^ and *Adar1p150*^-/-^*lfih1*^-/-^ mice^45–47^. However, the rescue was incomplete, suggesting the additional functions of ADAR1 that remain elusive.

We now report that the dsRNA binding activity, but not A-to-I editing activity, of ADAR1p150 protects cells from IFN-induced stress and death by preventing hyperactivation of PKR. ADAR1p150, even in its catalytically inactive form, competes with PKR to bind endogenous dsRNAs. Thus, ADAR1 has discrete roles in regulating dsRNA-mediated innate immunity: preventing MDA5 activation via RNA editing, and suppressing PKR through RNA binding competition. We demonstrate *in vivo* that the activation of PKR is responsible for the postnatal lethality of *Adar1*^-/-^*Ifih1*^-/-^ and *Adar1p150*^-/-^*lfih1*^-/-^ mice, with both *Adar1*^-/-^*Ifih1*^-/-^*Eif2ak2*^-/-^ mice and *Adar1p150*^-/-^*lfih1*^-/-^*Eif2ak2*^-/-^ surviving to adulthood. Most strikingly, we see the complete rescue and survival of *Adar1p150*^-/-^*lfih1*^-/-^*Eif2ak2*^-/-^, demonstrating that MDA5 and PKR signaling are the primary *in vivo* effectors of lethal autoinflammation following the loss of ADAR1p150. Our findings reconcile the *in vitro* and *in vivo* observations and demonstrate essential discrete roles for the RNA editing and binding activity of ADAR1p150 in preventing activation of MDA5 and PKR, respectively, in both human and mouse.

## Results

### Removal of PKR partially rescues *Adar1*^-/-^*Ifih1*^-/-^ mice

As ADAR1 prevented PKR activation in both human and mouse cell lines, and this is at least partially independent of RNA editing activity^24,33–42^, we reasoned that PKR may be activated in *Adar1*^-/-^*Ifih1*^-/-^ mice which die soon after birth. Loss of PKR alone did not modify survival of *Adar1*^-/-^ mice^30^, so we tested if it may play a role when MDA5 was also removed. PKR is a kinase activated by dsRNA-induced autophosphorylation, that in turn phosphorylates the translation initiation factor eIF2α leading to translational shutdown and the integrated stress response (ISR)^8,11,48^. We first assessed p-eIF2α, the marker of PKR activation in mouse cells due to the lack of p-PKR antibody for mouse samples, in primary tail fibroblasts from *Adar1*^+/+^*Ifih1*^-/-^, *Adar1^E861A/E861A^Ifih1*^-/-^ and *Adar1*^-/-^*Ifih1*^-/-^ mice. Both WT and E861A cells had low levels of p-elF2α. In contrast, the *Adar1*^-/-^ cells had elevated basal p-elF2α that was further increased upon IFNβ treatment (Figure 1A), consistent with previous results in human cells^24^. We then assessed primary kidney protein lysate from *Adar1*^+/+^*Ifih1*^-/-^, *Adar1^E861A/E861A^Ifih1*^-/-^ and *Adar1*^-/-^*lfih1*^-/-^ mice isolated on the day of birth. *Adar1*^-/-^*Ifih1*^-/-^ kidneys had elevated p-eIF2α levels (Figure 1B) and activation of the ISR gene expression program (Figures S1A-S1C), both indicative of PKR activation *in vivo.* These were both absent in *Adar1^E861A/E861A^Ifih1*^-/-^ mice indicating expression of a catalytically inactive ADAR1 protein was sufficient to prevent PKR activation *in vivo*.

**Figure 1.**
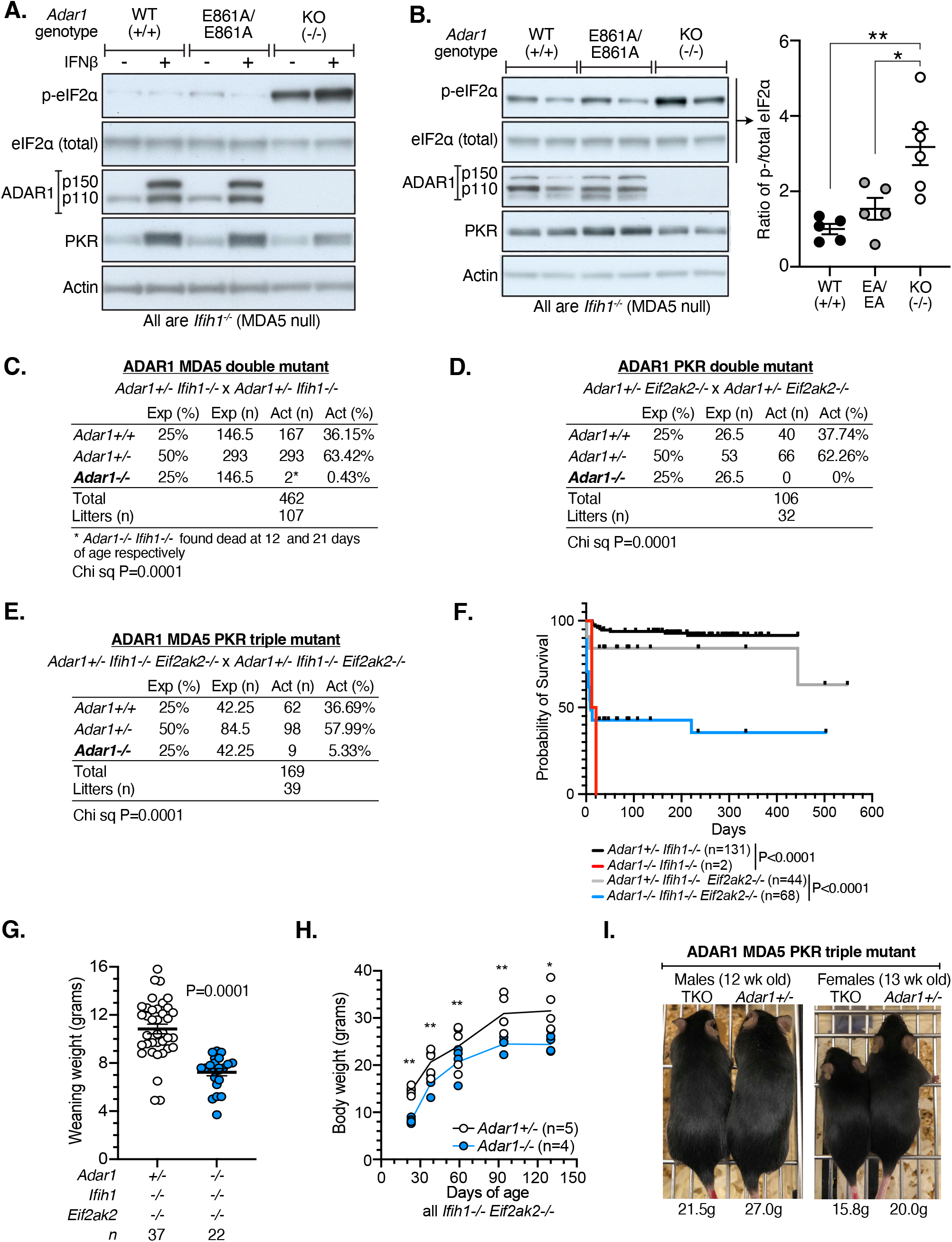
PKR activation causes the postnatal lethality of *Adar1*^-/-^*Ifih1*^-/-^ mice. (A) Western blot of murine tail fibroblasts generated from day of birth mice of the indicated genotypes treated for 24 hrs with mouse IFNβ or vehicle. Representative blot from n=2 mice. (B) Whole protein lysate from kidney collected on day of birth from *Adar1*^+/+^*Ifih1*^-/-^, *Adar1^E861A/E861A^Ifih1*^-/-^ and *Adar1*^-/-^*Ifih1*^-/-^ mice. Protein isolates from two independent mice per genotype are shown. Right, quantification of phospho-eIF2α and total eIF2α from Western blot from n=5 mice. Significance was determined by one-way ANOVA with multiple comparisons. (C) -(E) Breeding data for generation of *Adar1*^-/-^*Ifih1*^-/-^ (C), *Adar1*^-/-^*Eif2ak2*^-/-^ (D), and *Adar1*^-/-^*Ifih1*^-/-^*Eif2ak2*^-/-^ (E) mice. All breeding data are derived from inbreeding of pairs where only the *Adar1* allele is heterozygous (*Ifih1* and/or *Eif2ak2* are homozygous null). Statistical significance determined with Chi Squared test. See also Figures S1D and S1E. (F) Kaplan Meier survival curve of mice with the indicated genotypes. Number of animals as indicated for each genotype. Control mice included littermates with the *Adar1*^-/-^ genotype. Mice from additional breeding pairs with different genotypes to those shown in (E) were included in the survival curve. *P* values were calculated by log-rank tests. (G) Weaning weights of sibling *Adar1*^-/-^*Ifih1*^-/-^*Eif2ak2*^-/-^ and *Adar1*^+/-^*Ifih1*^-/-^*Eif2ak2*^-/-^ (control) mice. (H) Monthly weights of sibling *Adar1*^-/-^*Ifih1^-/-^*Eif2ak2*^-/-^* and *Adar1*^+/-^*Ifih1*^-/-^*Eif2ak2*^-/-^ (control) mice. See also Figure S1H. (I) Representative photos of male and female sibling *Adar1*^-/-^*Ifih1*^-/-^*Eif2ak2*^-/-^ and *Adar1*^+/-^*Ifih1*^-/-^*Eif2ak2*^-/-^ (control) mice at the indicated age. See also Figure S1G.

Having established that PKR was activated in *Adar1*^-/-^*Ifih1*^-/-^ mice, we tested if the early postnatal lethality of *Adar1*^-/-^*Ifih1*^-/-^ mice was due to PKR. We established *Adar1*^-/-^*Iih1*^-/-^*Eif2ak2*^-/-^ (ADAR1/MDA5/PKR protein deficient) animals with *Adar1^E861A/E861A^* as controls (Figures 1C–1E). The loss of MDA5 or PKR alone reproduced the previously reported results: normal representation at the day of birth (Figure S1D) but then completely penetrant early post-natal lethality for *Adar1*^-/-^*Ifih1*^-/-^ (Figure 1C)^21^ and no viable *Adar1*^-/-^*Eif2ak2*^-/-^ mice (Figure 1D)^30^ being recovered. In contrast, we recovered viable *Adar1^-/-^Ifih1*^-/-^*Eif2ak2*^-/-^ animals with ~40% surviving long term to adulthood (oldest >95 weeks old; Figures 1E–1F and S1E-S1F). The rescued *Adar1*^-/-^*Ifih1*^-/-^*Eif2ak2*^-/-^ animals were runted at weaning and this persisted throughout life (Figures 1G–1I and S1G-S1H). The *Adar1*^-/-^*Ifih1*^-/-^*Eif2ak2*^-/-^ are fertile and capable of breeding. The *Adar1^E861A/E861A^* were rescued by loss of MDA5 (Figures S2A-S2B), as expected and previously reported^32^, but not PKR (Figure S2C). The *Adar1^E861A/E861A^Ifih1*^-/-^*Eif2ak2*^-/-^ mice were recovered at the expected mendelian ratio, were slightly runted at weaning and then had a normal longterm survival (Figures S2D-S2H), comparable to our previous analysis of the *Adar1^E861A/E861A^Ifih1*^-/-^ animals^32^. These data demonstrate that when the ADAR1 protein is absent, but not when an editing deficient protein is expressed, both PKR and MDA5 are activated *in vivo*. Therefore, functions of ADAR1 independent of A-to-I editing are required to prevent PKR activation *in vivo*.

### ADAR1 suppresses IFN-induced PKR activation independent of RNA editing in human cells

To understand the basis for PKR engagement in an ADAR1 protein deficient setting, we took advantage of human HEK293T cells with homozygous ADAR1 null (ADAR1^KO^; no ADAR1 protein expressed) and RNA editing deficient knock-in mutation (ADAR1^E912A^, homologous to mouse ADAR1^E861A^)^49^ that we recently generated and characterized^50^. The majority of A-to-I RNA editing was absent in both ADAR1^KO^ and ADAR1^E912A^ cells^50^. No significant cellular phenotype or basal ISG induction was observed in either ADAR1 mutant cell genotype (Figure S3A and Table S1), consistent with previous findings in independently generated ADAR1 deficient HEK293T cells^21,24^. However, when treated with IFNα the proliferation of ADAR1^KO^ cells, but not of WT or ADAR1^E912A^ cells, was significantly reduced (Figures 2A and S3B). Consistent with previous findings^24^, IFNα treatment induced a higher level of PKR phosphorylation in ADAR1^KO^ cells than in WT cells (Figure 2B). Importantly and consistent with our *in vivo* observations and some reports^33,36^, expression of an editing-dead protein was sufficient to prevent PKR activation (Figure 2B). This contrasts with previous studies that used complementation of ADAR1^KO^ cells with overexpression of ADAR1 mutant proteins to conclude that editing activity was required to fully suppress PKR activation^24,37^. Note that as an ISG itself, total PKR is also increased in response to IFNα treatment which is independent of its phosphorylation and activation^24^. There was formation of G3BP1 containing stress granules in the ADAR1^KO^ cells, but not in WT or ADAR1^E912A^ cells, following IFNα treatment (Figure 2C). ADAR1^E912A^ cells form G3BP1 stress granules upon arsenite treatment, demonstrating that the lack of ADAR1-mediated A-to-I editing does not prevent granule formation *per se* (Figure S3C). This is consistent with a recent report showing that ADAR1 RNA-binding but not RNA-editing activity was required for partial inhibition of stress granule formation in response to arsenite treatment, although the effects were more subtle than for IFNα-induced stress granules reported here^43^. To establish the requirement of PKR, we used short hairpin RNAs (shRNAs) to knockdown PKR expression (Figure 2D). The knockdown of PKR prevented both the formation of stress granules (Figure 2E) and the reduced proliferation (Figure 2F) of ADAR1^KO^ cells in response to IFNα. Therefore, expression of an editing-deficient ADAR1 protein from the endogenous locus is sufficient to prevent PKR activation, PKR-dependent formation of stress granules and reduced proliferation of cells in response to IFNα treatment *in vitro*.

**Figure 2.**
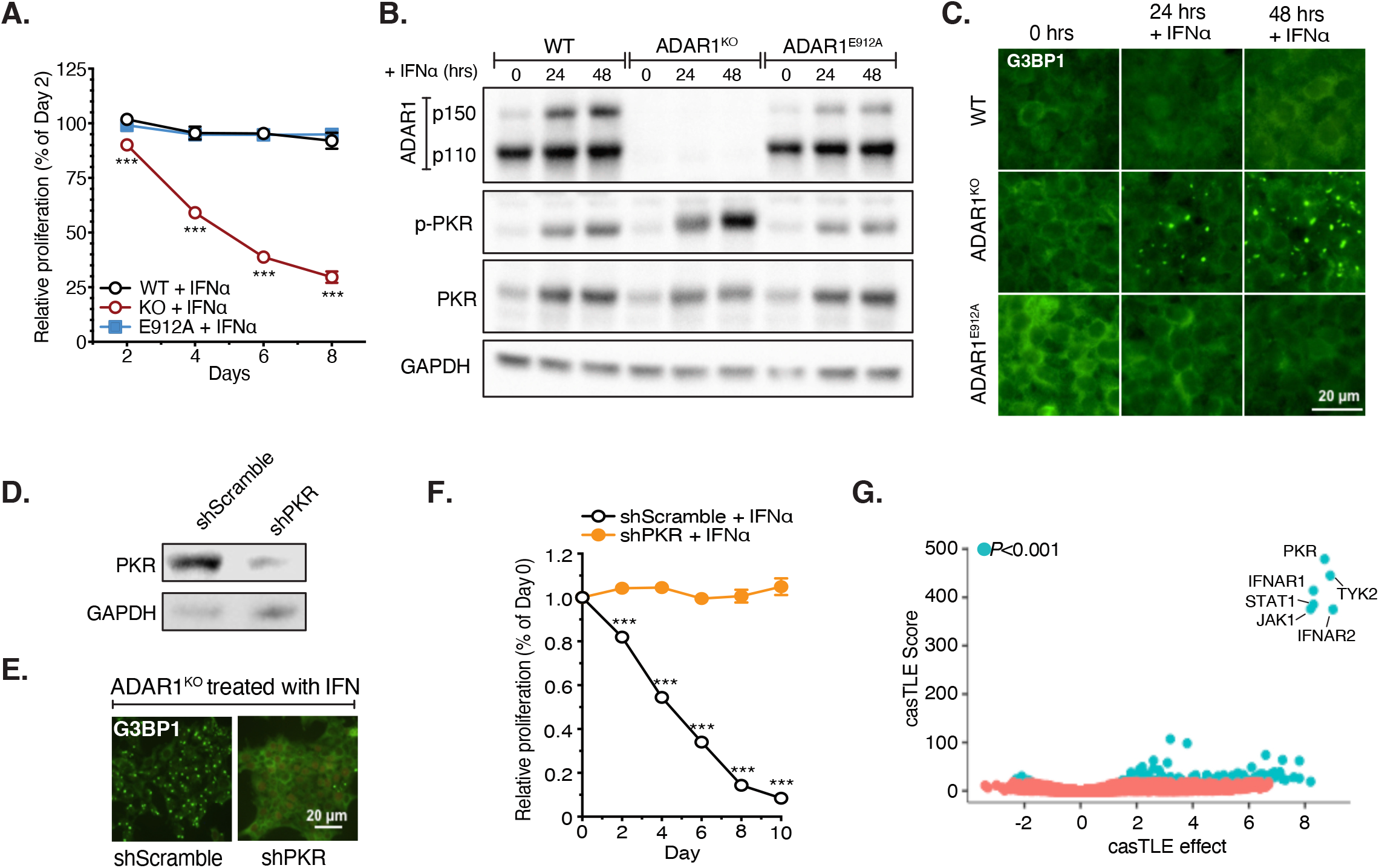
ADAR1 suppresses IFN-induced PKR activation in human cells. (A) The effect of IFNα treatment on the cell proliferation of WT, ADAR1^KO^, and ADAR1^E912A^ HEK293 cells. Cells were treated with 10 ng/ml IFNα. The detailed procedure of the assay was described in Figure S3B and Method. Data from 3 replicates are shown as the mean +/- SEM. Significance determined by 2-way ANOVA with multiple comparisons. (B) Western blot analysis of PKR phosphorylation. Cells were treated with 10 ng/ml IFNα for 0, 24, or 48 hours. Representative result is shown from three replicates. See Data S1 for the other replicates. (C) Stress granule formation upon IFNα treatment in WT, ADAR1^KO^, and ADAR1^E912A^ HEK293T cells. Cells were treated as panel (B) followed by immunofluorescence detection of G3BP1 (green). (D) Knockdown of PKR by shRNA (shPKR) in ADAR1^KO^ HEK293T cells, revealed by Western blot. Representative result is shown from replicates. See Data S1 for the other replicates. (E) Stress granule staining (G3BP1) in scramble shRNA or PKR shRNA-treated ADAR1^KO^ cells. (F) The effect of PKR depletion on the cell proliferation. Data from 3 replicates are shown as the mean +/- SEM. Significance determined by 2-way ANOVA with multiple comparisons. (G) Genome-wide CRISPR KO screening in ADAR1^KO^ U937 cell. Screen data were analyzed by casTLE^67^. Data from two repeats are presented. CasTLE effect indicates the most likely effect size as determined by casTLE. CasTLE score indicates the confidence level in that effect size. CasTLE *P* value is the estimated *P* value from the casTLE score. See also Figure S3F.

To better understand the genetic landscape of this response, we undertook a genome-wide CRISPR loss of function screen in the ADAR1^KO^ U937 cells treated with IFNα (Figures S3D-S3F). As a suspension cell line, the U937 cells were more amenable to screening methods than HEK293T cells. IFNαtreatment reduced the proliferation of ADAR1^KO^ U937 cells, as we observed in HEK293Ts (Figures S3D-S3E). The most highly enriched loss of function candidate was PKR (Figure 2G and Table S2). The enriched candidates also included loss of the core components of the type I IFN signaling pathway, such as IFNAR1/2, TYK2, STAT1, and JAK1 (Figure 2G and Table S2), validating the robustness of the screen. Importantly, no additional genes were significantly enriched in the screen, suggesting that PKR was the primary mediator accounting for the IFNα-induced phenotypes in these cells. These data demonstrate that the presence of an ADAR1 protein was required to suppress PKR activation and that this is a species conserved editing independent function of ADAR1.

### MDA5-induced IFN signal activates PKR in the absence of ADAR1

In mice, MDA5-dependent sensing of unedited cellular RNA is activated when ADAR1 is deleted or its editing activity is lost^19–21^. However, ADAR1^KO^ or ADAR1^E912A^ human HEK293T cells do not have spontaneous innate immune activation, likely due to very lowly expressed MDA5 (Table S1)^50^, necessitating the treatment with IFNα. To determine if expression of MDA5 in the absence of IFN treatment was sufficient to recapitulate the pathway activation in HEK293T cells, we established a doxycycline inducible MDA5 expression system in WT, ADAR1^KO^ and ADAR1^E912A^ HEK293Ts (Figures 3A–3B). Upon induction of MDA5 expression, IFN and ISGs were induced in both ADAR1^KO^ and ADAR1^E912A^ cells (Figure 3B). Interestingly, there was a rapid increase in p-PKR levels (Figure 3C) and PKR-dependent proliferation arrest (Figures 3D–3E) in the ADAR1^KO^ cells compared to WT or ADAR1^E912A^ cells. Note there was a slight increase of p-PKR levels in ADAR1^E912A^ cells relative to WT (Figure 3C). We reasoned this was due to the IFN signal induced by MDA5-mediated detection of unedited RNAs in ADAR1^E912A^ cells (Figure 3B), since IFN modestly increased p-PKR level in WT or ADAR1^E912A^ cells (Figure 2B). The activation of PKR upon MDA5 induction appeared to be mediated by IFN signaling, as inhibition of the signaling cascade downstream of MDA5/MAVS (TBK1i) or Type I IFN receptor signaling (JAKi) prevented the activation of PKR (Figure 3F). Furthermore, knockdown of MDA5 did not abolish IFN treatment-induced PKR activation (Figure 3G), suggesting that PKR activation was not directly dependent on MDA5. These data are consistent with PKR activation being dependent on IFN-induced *de novo* transcription as previously described^24^. Taken together these data in HEK293T cells demonstrate that in the absence of ADAR1 protein, PKR activation required type I IFN signaling. The stimulus can either be derived from binding by cytokine of the type I IFN receptors or from MDA5 engaging with unedited dsRNA within the cell.

**Figure 3.**
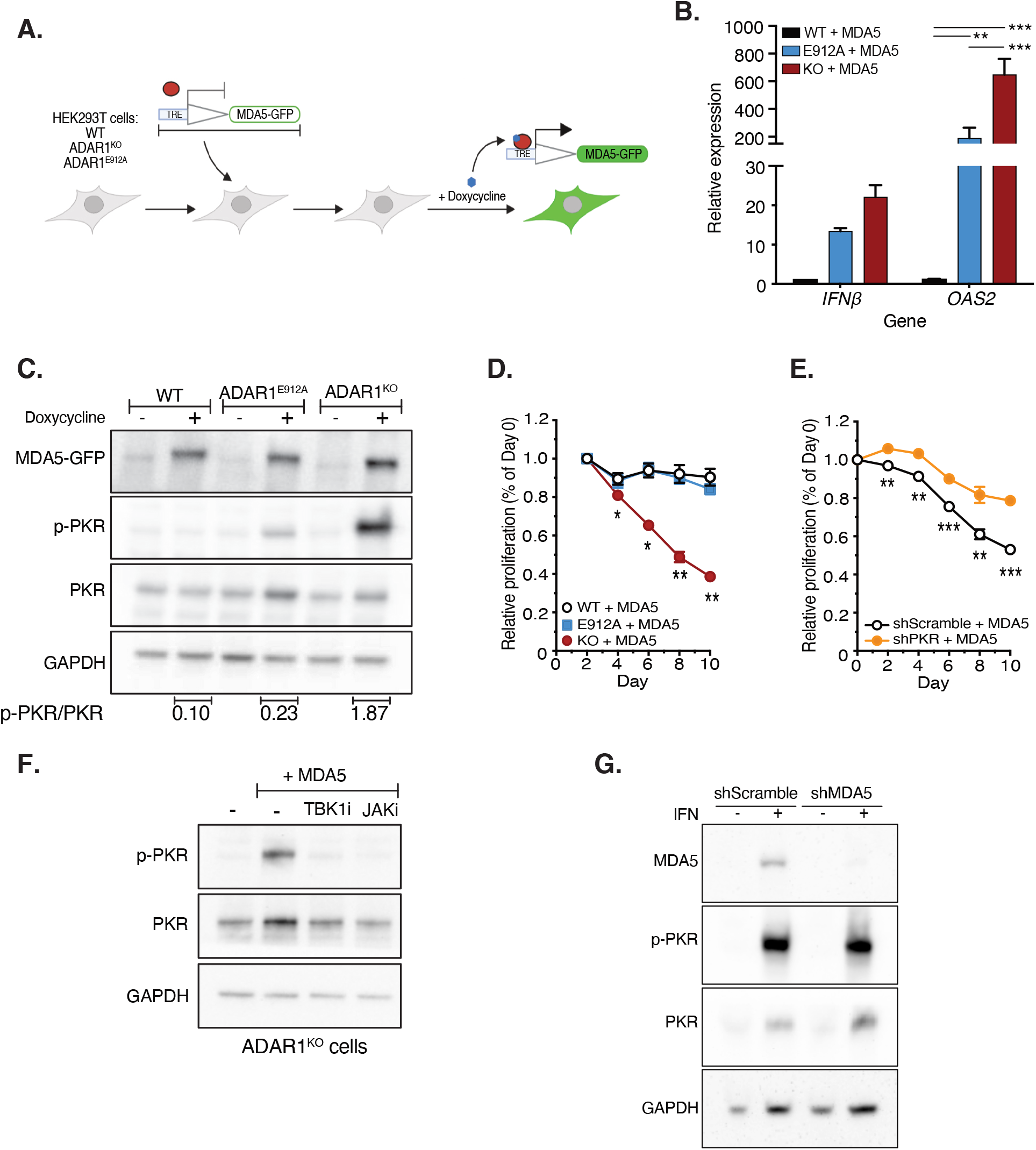
MDA5-induced IFN signaling activates PKR in the absence of ADAR1. (A) Experimental outline for results shown in this figure. 1μg/ml doxycycline was added to the cell for 48 hours to induce MDA5 overexpression. (B) Real-time PCR measurement of *IFNB* and *OAS2* in WT, ADAR1^KO^, and ADAR1^E912A^ cells that overexpressed MDA5. Data from 3 replicates are shown as the mean +/- SEM. Significance was determined by multiple t test. (C) Western blot analysis of PKR phosphorylation in WT, ADAR1^KO^, and ADAR1^E912A^ cells plus/minus MDA5 overexpression. Representative result is shown from replicates. See Data S1 for the other replicates. (D) Cell proliferation in WT, ADAR1^KO^, and ADAR1^E912A^ cells that overexpressed MDA5. (E) The effect of PKR depletion on cell proliferation of ADAR1^KO^ cells that overexpressed MDA5. (F) The effect of TBK1 or JAK inhibition on MDA5-induced PKR activation in ADAR1^KO^ cells. 10μM MRT67307 (TBK1 inhibitor) or 5μM Ruxolitinib (JAK inhibitor) along with 1 μg/ml doxycycline was added to the cell for 48 hours. (G) Western blot analysis of PKR phosphorylation in scramble shRNA or MDA5 shRNA-treated ADAR1^KO^ cells. ADAR1^KO^ cells were infected with lentivirus expressing shScramble or shMDA5 followed by IFNα treatment for 48 hours. Data from 3 replicates are shown as the mean +/- SEM in (D) and (E). Significance determined by 2-way ANOVA with multiple comparisons.

### ADAR1 regulates the interaction of MDA5 and PKR in monocytes

In HEK293T cells, PKR activation required IFN treatment^24^ or MDA5 activation. However, *Adar1*^-/-^*Ifih1*^-/-^ mice, lacking MDA5 or exogenous IFN treatment, have spontaneous PKR activation. We reasoned that the requirement of exogenous IFN signal for PKR activation may vary amongst cell types^24,33–43^. To overcome the need for IFN treatment, we explored the U937 monocytic cell line and differentiated it to macrophage-like cells with phorbol myristate acetate (PMA) treatment (Figure S4A)^51^. Differentiation of U937 to macrophages increased the overall expression level of ISGs (Figure S4B and Table S3). The ADAR1^KO^ U937 cells had a subtle basal change of ISG expression (Figure 4A and Table S3). However, when U937 cells were differentiated into macrophages, there was a significantly elevated ISG signature (Figures 4A–4B and Table S3), PKR phosphorylation (Figure 4C), and cell death (Figure 4D) in ADAR1^KO^ cells compared to controls. The depletion of MDA5 prevented ISG expression (Figures 4E–4F and S4C), but it did not fully rescue PKR activation (Figure 4E), cell viability (Figure 4G), or stress granule formation (Figure 4H) caused by ADAR1 loss. In contrast, depletion of PKR rescued cell viability (Figure 4G) and prevented stress granule formation (Figure 4H), but it did not reduce ISG expression (Figure 4F). These results demonstrated that loss of ADAR1 can activate PKR in particular human cell types in the absence of IFN treatment or MDA5 induction. This may be due to variable expression levels of both sensors and RNA substrates in different cell types. Importantly, it is consistent with the *in vivo* data where PKR was activated in *Adar1*^-/-^*Ifih1*^-/-^ mice (Figures 1A–1B), despite there being no MDA5 or IFN treatment.

**Figure 4.**
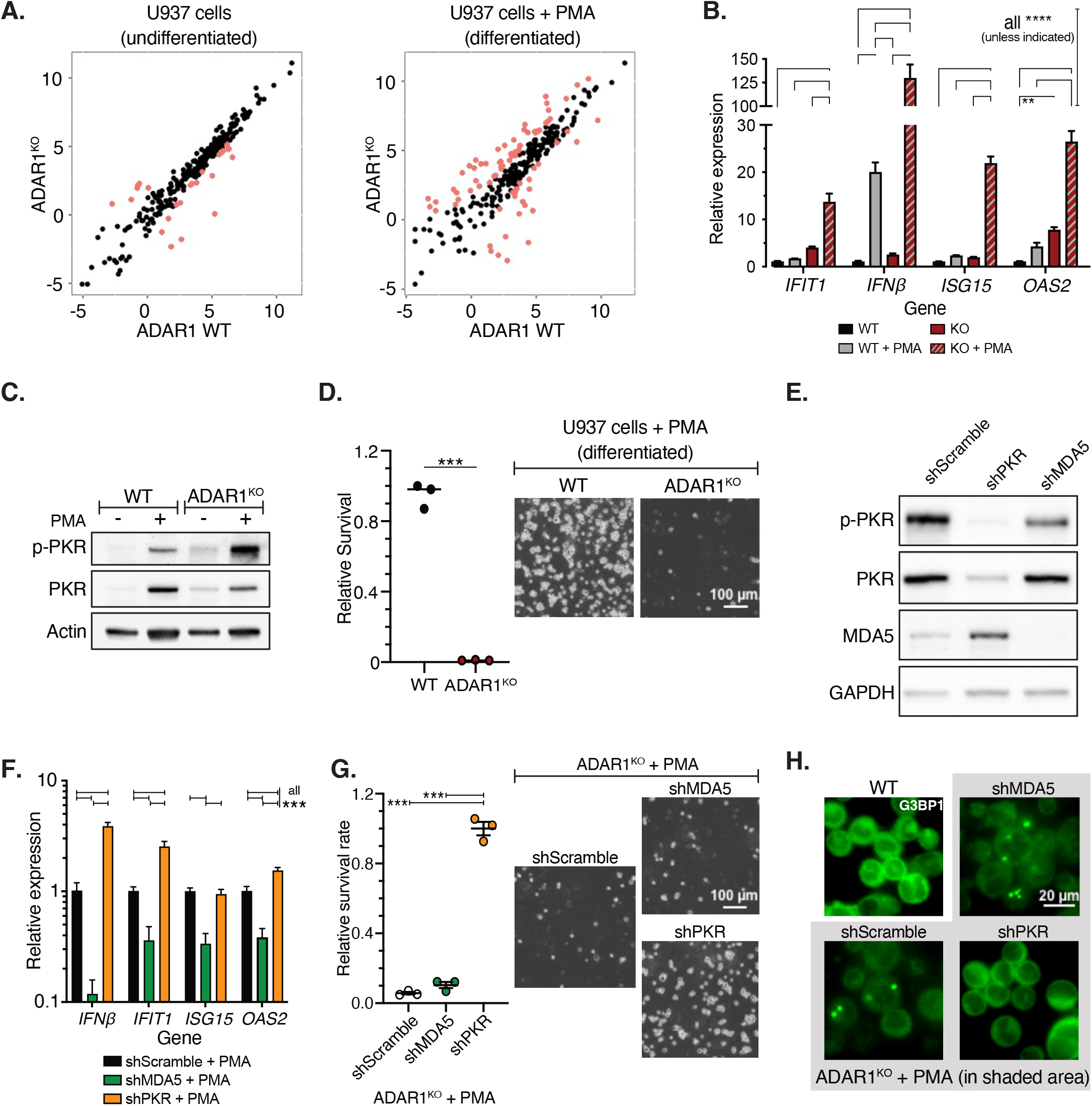
ADAR1 regulates MDA5 and PKR in macrophages. (A) Expression comparison of ISGs between WT and ADAR1^KO^ in U937 cells (left) and U937-differentiated macrophages (right). The gene expression levels were measured by TPM (Transcripts Per Kilobase Million) and log2 transformed. Genes with more than 2-fold change are in pink. (B) Real-time PCR measurement of representative ISGs before or after PMA-induced differentiation in WT and ADAR1^KO^ U937 cells. Data from 3 replicates are shown as the mean +/- SEM. Significance was determined by multiple t test. (C) Western blot analysis of PKR phosphorylation before or after PMA-induced differentiation in WT and ADAR1^KO^ U937 cells. Representative result is shown from replicates. See Data S1 for the other replicates. (D) The survival of WT and ADAR1^KO^ U937-differentiated macrophages. Left, FACS analysis of the survived macrophages, normalized to input amount of U937 cells for differentiation. Right, a representative bright-field view of the confluency of WT and ADAR1^KO^ macrophage. Significance was determined by unpaired t test with 3 replicates. (E) Depletion of PKR or MDA5 by shRNAs in ADAR1^KO^ U937 cells followed by PMA-induced differentiation. The expression of MDA5, PKR, and phosphorylated PKR was revealed by Western blot. Representative result is shown from replicates. See Data S1 for the other replicates. (F) The effect of MDA5 depletion on ISGs expression in ADAR1^KO^ U937-differentiated macrophages. Data from 3 replicates are shown as the mean +/- SEM. Significance was determined by multiple t test. (G) The effect of PKR or MDA5 depletion on the survival of ADAR1^KO^ U937-differentiated macrophages. Left, FACS analysis of the survived macrophages, normalized to input amount of U937 cells for differentiation. Right, a representative bright-field view of the confluency of the macrophage. Significance was determined by unpaired t test with 3 replicates. (H) Stress granule staining in macrophages of WT, ADAR1^KO^, and ADAR1^KO^ cells rescued with PKR or MDA5 depletion.

### Cytoplasmic ADAR1p150 RNA binding suppresses PKR activation

Next, we sought to understand how ADAR1 restricted PKR activation. As the two ADAR1 isoforms display distinct cellular localization with p110 in the nucleus and p150 cytoplasmic, we first determined which isoform was required to suppress PKR activation. Upon re-introduction of p110 or p150 into ADAR1^KO^ HEK293T cells (Figure S5A), only p150 significantly suppressed PKR phosphorylation (Figure 5A) and stress granule formation (Figure 5B). In addition to the distinct cellular localization, p150 differs from p110 by having a N-terminal Zα domain^1^. To distinguish whether p150’s function on PKR suppression was Zα domain dependent or localization dependent, we employed ADAR2, naturally lacking a Zα domain, for further verification. Wild-type or a cytoplasm-localized ADAR2 mutant (cytoADAR2; lacking the nuclear localization signal)^50^ was introduced into the ADAR1^KO^ cells with the appropriate cellular localization (Figure S5A). Like ADAR1p150, cytoADAR2 suppressed PKR phosphorylation (Figure 5A) and stress granule formation (Figure 5B) following IFN treatment, in contrast to WT ADAR2 and ADAR1p110. Previous reports suggested that p110 plays a role in suppression of PKR activation^24,37^. We observed quite variable effects from p110 overexpression with no statistical difference to control empty vector. However together with the rescue by cytoADAR2, our findings demonstrate that the cytoplasmic localization of an ADAR protein played a critical role in suppressing IFN-induced PKR activation, whereas the Zα domain was not essential.

**Figure 5.**
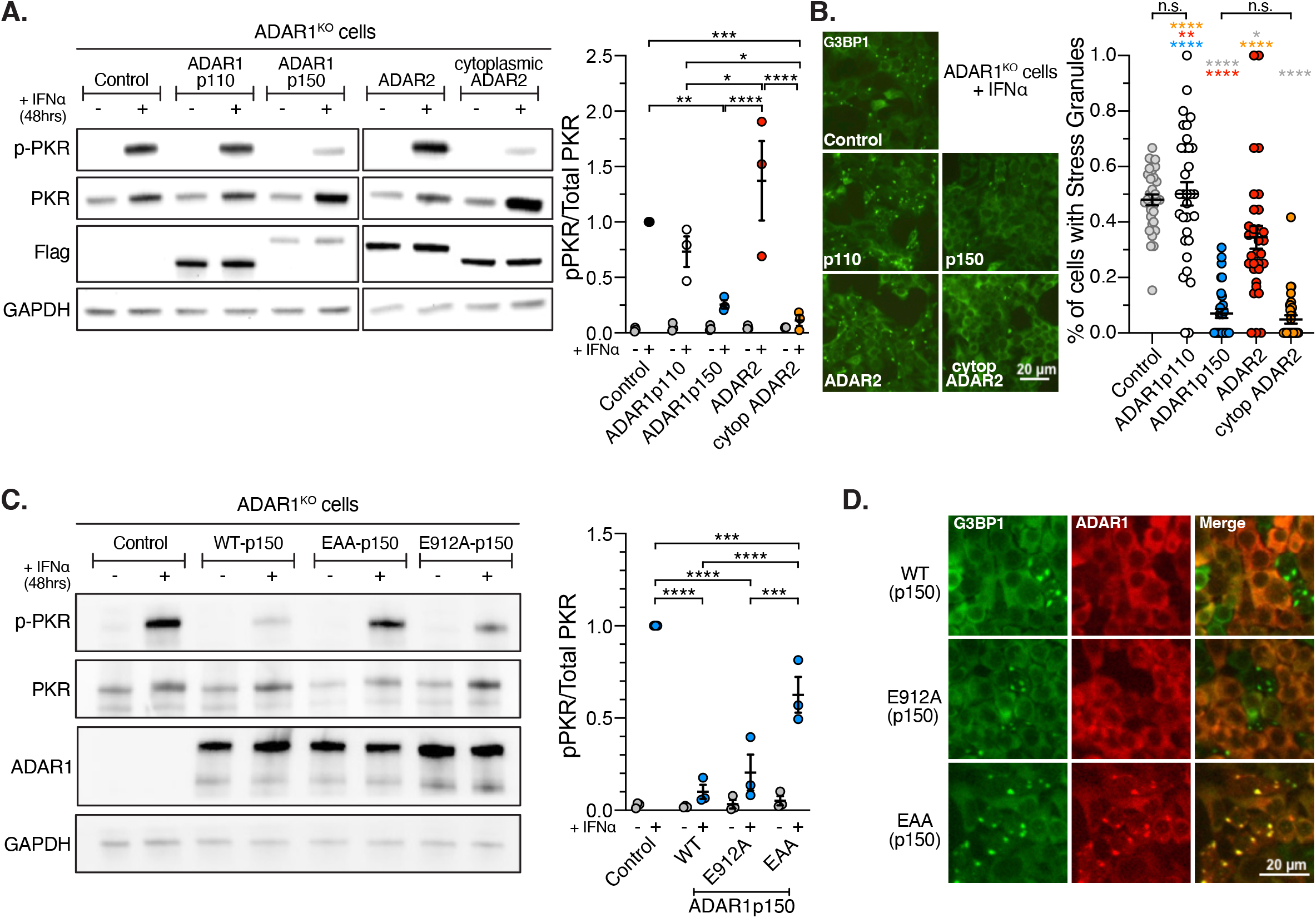
RNA binding activity of cytoplasmic ADAR is required for inhibition of PKR. (A) Western blot analysis of PKR phosphorylation in ADAR1^KO^ cells rescued with different ADAR proteins. ADAR1^KO^ cells were infected with lentivirus expressing ADAR1p110, ADAR1p150, ADAR2, or ADAR2 mutant which lacked a nuclear-localization signal (cytoplasmic ADAR2), followed by IFNα treatment for 48 hours. Representative result is shown from three replicates. See Data S1 for the other replicates. Right, quantification of pPKR level normalized to total PKR from three replicates. (B) Stress granule staining in ADAR1^KO^ HEK293T cells rescued with different ADAR proteins. ADAR1^KO^ cells were infected with lentivirus expressing ADAR1p110, ADAR1p150, ADAR2, or ADAR2 mutant which lacked a nuclear-localization signal (cytoADAR2), followed by IFNα treatment for 48 hours and immunofluorescence of G3BP1. Right, analysis of the percentage of cells with stress granules in left. Each data point represents the percentage of stress granule positive cells from random fields. In total, more than 290 cells were assessed. (C) Western blot analysis of PKR phosphorylation in ADAR1^KO^ cells rescued with different ADAR1p150 proteins. ADAR1^KO^ cells were infected with lentivirus expressing WT, E912A (editing deficient mutant), or EAA (RNA binding deficient mutant) ADAR1p150 followed by IFNα treatment for 48 hours. Representative result is shown from three replicates. See Data S1 for the other replicates. Right, quantification of pPKR level normalized to total PKR from three replicates. (D) Stress granule staining in ADAR1^KO^ HEK293T cells rescued with different ADAR1p150 proteins. ADAR1^KO^ cells were infected with lentivirus expressing WT, E912A (editing deficient mutant), or EAA (RNA binding deficient mutant) ADAR1p150 followed by IFNα treatment for 48 hours and immunofluorescence detection of G3BP1 (green) and ADAR1 (red).

Having established the centrality of ADAR1p150 in the suppression of PKR activation, we tested the rescue of ADAR1^KO^ HEK293T cells with ADAR1p150 WT, an RNA editing deficient mutant (E912A), or an RNA binding deficient mutant (EAA)^52^. Both the WT and E912A mutant p150 were equally effective at preventing IFN-induced PKR activation (Figure 5C). In contrast, the EAA mutant lacking RNA binding activity was the least effective at complementing ADAR1 loss, demonstrating that RNA binding was the dominant function for suppressing PKR activation. This contrasts with a previous report that concluded that RNA-binding and RNA-editing were equally important for suppression of PKR activation^24^. The reason for the difference to these results is not clear, however further analysis of stress granule formation under the same conditions was consistent with a specific requirement for RNA-binding, not RNA-editing for suppression of PKR activation (Figure 5D). Taken together, these data demonstrate that ADAR1p150 primarily suppressed PKR activation through its RNA binding activity but independently of A-to-I editing.

### ADAR1p150 binding to dsRNAs prevents PKR binding

To understand how RNA binding by ADAR1p150 could suppress PKR activation we first assessed whether ADAR1 could directly bind to PKR. An exogenous flag-tagged PKR co-immunoprecipitated (co-IP) small amounts of ADAR1 relative to input, but the interaction was reduced upon RNase digestion and increased after adding dsRNAs in the co-IP reaction (Figure 6A). This indicated that the interaction of ADAR1 and PKR was largely dsRNA-dependent, and that ADAR1 and PKR can bind the same RNA targets.

**Figure 6.**
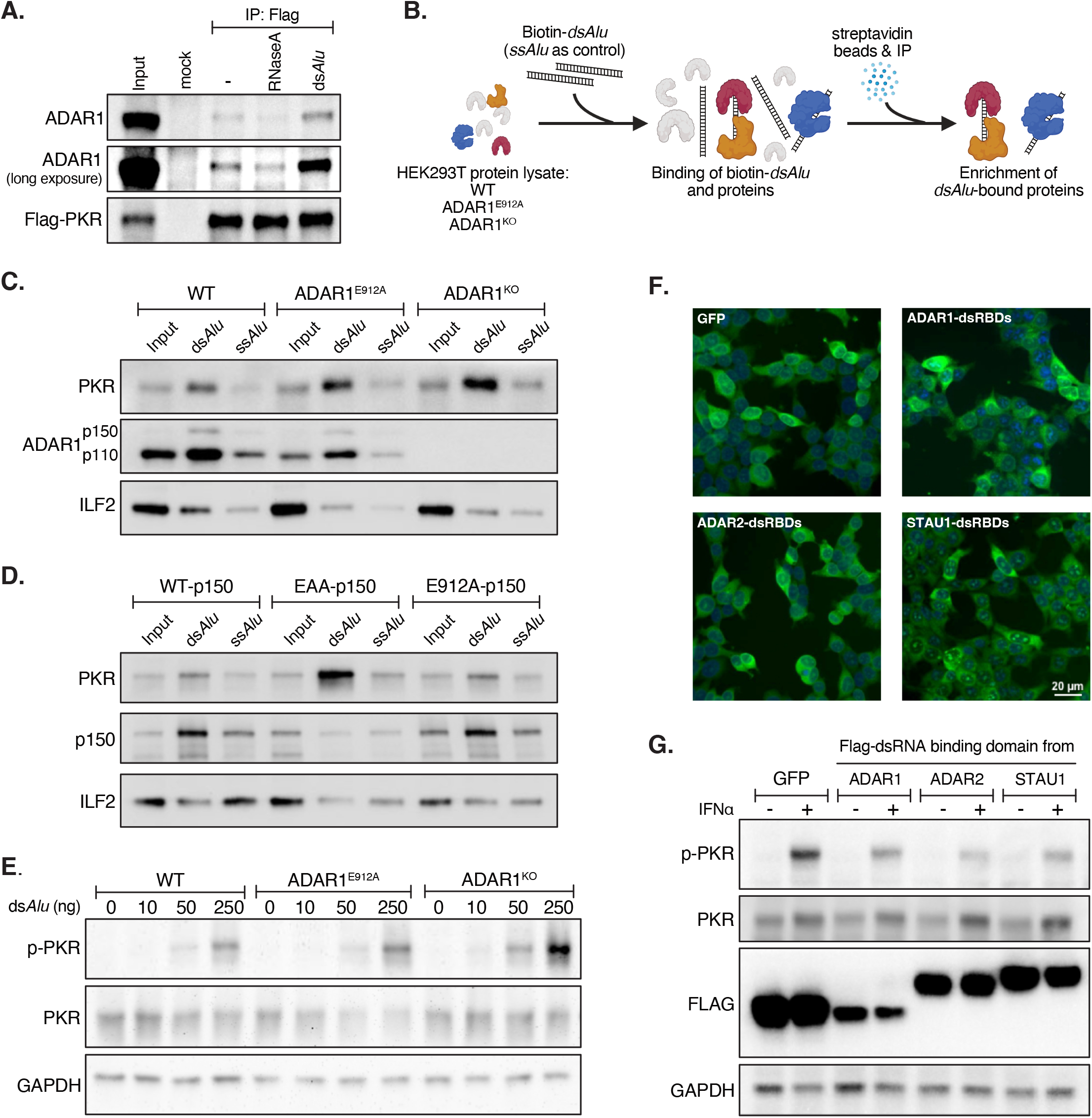
ADAR1p150 competes with PKR to bind dsRNA for PKR inhibition. (A) Analysis of the interaction between ADAR1 and PKR by exogenous flag-tagged PKR (flag-PKR) co-immunoprecipitation. Flag-PKR protein was precipitated by flag antibody from the cell lysate of WT HEK293T cells in the presence of RNase A or dsRNA derived from an *Alu* repeat element (ds*Alu*). (B) Schematic of Biotin-labelled RNA pulldown assay shown in (C) to test the dsRNA binding capacity of PKR in WT, ADAR1^KO^, and ADAR1^E912A^ HEK293T cells. The cell lysate was incubated with biotin-labeled dsRNA (ds*Alu*) or ssRNA (ss*Alu*) to enrich the RNA binding proteins, followed by western blot. See Method for detail. (C) Western blot of RNA pulldown described in (B). (D) Biotin-labeled RNA pulldown revealed dsRNA binding capacity of PKR in the presence of different p150 mutants. Cell lysates from ADAR1^KO^ cells rescued with WT, EAA, or E912A p150 were used for the Biotin-labeled RNA pulldown assay. (E) The sensitivity of PKR to dsRNA stimulation in WT, ADAR1^KO^, and ADAR1^E912A^ HEK293T cells. Different amounts of dsRNAs were transfected in the cells for 6 hours. Phosphorylation of PKR analyzed by Western blot. See also Figures S6A and S6B. (F) The dsRNA binding domains from ADAR1, ADAR2, or STAU1 were overexpressed in ADAR1^KO^ cells. Localization of the domains was imaged using shown by anti-Flag immunofluorescence. (G) Phosphorylation of PKR analyzed by Western blot following overexpression of the dsRNA binding domains from ADAR1, ADAR2, or STAU1 in ADAR1^KO^ cells (imaged in panel F) and the cells were then treated with IFNα for 48h. Representative result is shown from replicates. See Data S1 for the other replicates.

Since ADAR1 and PKR could target common RNA substrates, we hypothesized that ADAR1 might compete with PKR to bind these dsRNAs. To test that, *in vitro* transcribed dsRNAs were used to precipitate proteins from cell lysates to determine the affinity between dsRNAs and PKR in the presence or absence of ADAR1 (Figure 6B). DsRNA enriched more PKR protein in the absence of ADAR1 (ADAR1^KO^) than in the presence of an ADAR1 protein (WT or ADAR1^E912A^) (Figure 6C). Background binding to the *ssAlu* by PKR and ADAR1 is likely due to antisense transcription by T7 polymerase leading to the formation of small amounts of dsRNA or the secondary structure of single *Alu* elements^53^. The same assay was performed using cell lysate from ADAR1^KO^ cells expressing different p150 mutants. The loss of ADAR1p150 RNA binding (EAA mutant) increased the association of the dsRNAs with PKR, compared to when either WT or editing deficient (E912A mutant) ADAR1p150 were re-expressed (Figure 6D). These results demonstrate that competition for dsRNA binding between ADAR1p150 and PKR can occur *in vitro*. Based on a competition model, we speculated that ADAR1 loss would lower the threshold for activation of PKR by dsRNAs. When either *in vitro* transcribed dsRNAs or polyI:C were titrated into the cells, phosphorylation and activation of PKR occurred with a lower level of dsRNAs in ADAR1^KO^ cells than in WT or editing deficient E912A cells (~5 fold more sensitive in ADAR1^KO^ cells) (Figures 6E, and S6A-S6B).

We then tested if ADAR1’s dsRNA binding domains (dsRBDs) alone would be sufficient to suppress PKR activation. There are three dsRBDs shared between ADAR1p110 and ADAR1p150, with the third containing the nuclear localization signal. To overexpress dsRBDs in the cytoplasm, we cloned the first two dsRBDs from ADAR1 and introduced it to the ADAR1^KO^ cells and confirmed the cytoplasmic localization (Figure 6F). Strikingly, the overexpressed dsRBDs alone were sufficient to suppress PKR activation (Figure 6G). Extending this observation, cytoplasmic expression of the dsRBDs from other dsRNA binding proteins (dsRBPs) ADAR2 and STAU1 (Figure 6F) suppressed PKR activation (Figure 6G), indicative of a more generalizable model of competition between dsRBPs for cytoplasmic dsRNA substrates. Indeed, other dsRBPs, such as STAU1, were reported to suppress PKR activation through binding dsRNA substrates^54^. Collectively, these data demonstrate that binding, but not editing, of cytoplasmic dsRNAs by ADAR1p150 can inhibit PKR activation by cellular dsRNAs.

### Removal of PKR and MDA5 fully rescues *Adar1p150*^-/-^ mice

The survival of *Adar1*^-/-^*Ifih1*^-/-^*Eif2ak2*^-/-^ mice to adulthood unlike the *Adar1*^-/-^*Ifih1*^-/-^ was consistent with the model of PKR activation occurring when there was no ADAR1 protein present. However, the *Adar1*^-/-^*Ifih1*^-/-^*Eif2ak2*^-/-^ mice were runted with high rates of post-natal lethality in the first 2-4 weeks after birth (Figure 1F). As we demonstrated *in cellulo*, it was the presence of ADAR1p150 protein that regulated both MDA5 and PKR activation. We reasoned that the early death of *Adar1*^-/-^*Ifih1*^-/-^*Eif2ak2*^-/-^ mice could be attributable to a lack of ADAR1p110. *Adar1p110*^-/-^ animals survived to birth, but less than 20% survived to two weeks of age^7^. The *Adar1p110*^-/-^animals did not show evidence of an active innate immune response and loss of MDA5 did not modify the survival of the *Adar1p110*^-/-7^. Since the survival rate of *Adar1*^-/-^*Ifih1*^-/-^*Eif2ak2*^-/-^ mice was comparable to that of *Adar1p110*^-/-^ *mice*^7^, we predicted that *Adar1p150^-/-^* mice should be fully rescued by removal of both MDA5 and PKR. To directly test this, we used an *Adar1p150*^-/-^ allele (p.L196CfsX6) that we recently identified and characterized as a p150 isoform specific loss of function mutant^55^.

We first determined if the loss of MDA5 alone could rescue *Adar1p150*^-/-^. The *Adar1p150*^-/-^*Ifih1*^-/-^ animals were underrepresented and the majority died within the first month of life (Figures 7A–7B). This is concordant with the recent results from the previously described *Adar1p150*^-/-^ allele crossed to an *Ifih1*^-/-^^46^. We then established crosses to generate *Adar1p150*^-/-^*Ifih1*^-/-^*Eif2ak2*^-/-^ mice. Strikingly, we recovered *Adar1p150*^-/-^*Ifih1*^-/-^*Eif2ak2*^-/-^ at the expected Mendelian ratio (Figures 7C and S7A) and the rescued mice did not have early postnatal lethality as was seen with the *Adar1*^-/-^*Ifih1*^-/-^*Eif2ak2*^-/-^ mice that lack both p110 and p150 (Figure 7D; oldest *Adar1p150*^-/-^*Ifih1*^-/-^*Eif2ak2*^-/-^ >42 weeks old). The *Adar1p150*^-/-^*Ifih1*^-/-^*Eif2ak2*^-/-^ mice had normal weaning weight (Figure 7E), in contrast to the runting observed for the *Adar1*^-/-^*Ifih1*^-/-^*Eif2ak2*^-/-^ (Figures 1G–1H). The adult *Adar1p150*^-/-^*Ifih1*^-/-^*Eif2ak2*^-/-^ mice (males and females) were fertile (Figure S7A) and also had healthy outward appearance (Figure 7F) and a normal weight (Figure S7B), and were comparable to control genotypes following histopathology assessment (Data S2). We additionally recovered *Adar1p150*^-/-^*Ifih1*^-/-^*Eif2ak2*^+/-^ mice (PKR heterozygous) at the expected Mendelian ratio and these have demonstrated normal long-term survival (Figures S7C-7D; oldest >45 weeks of age), indicating a dosage dependent mechanism of PKR activation. These data demonstrate that MDA5 and PKR activation account for the essential *in vivo* functions of ADAR1p150 protein in mice and the combined loss of both can fully rescue the *Adar1p150*^-/-^ mice to adulthood.

**Figure 7.**
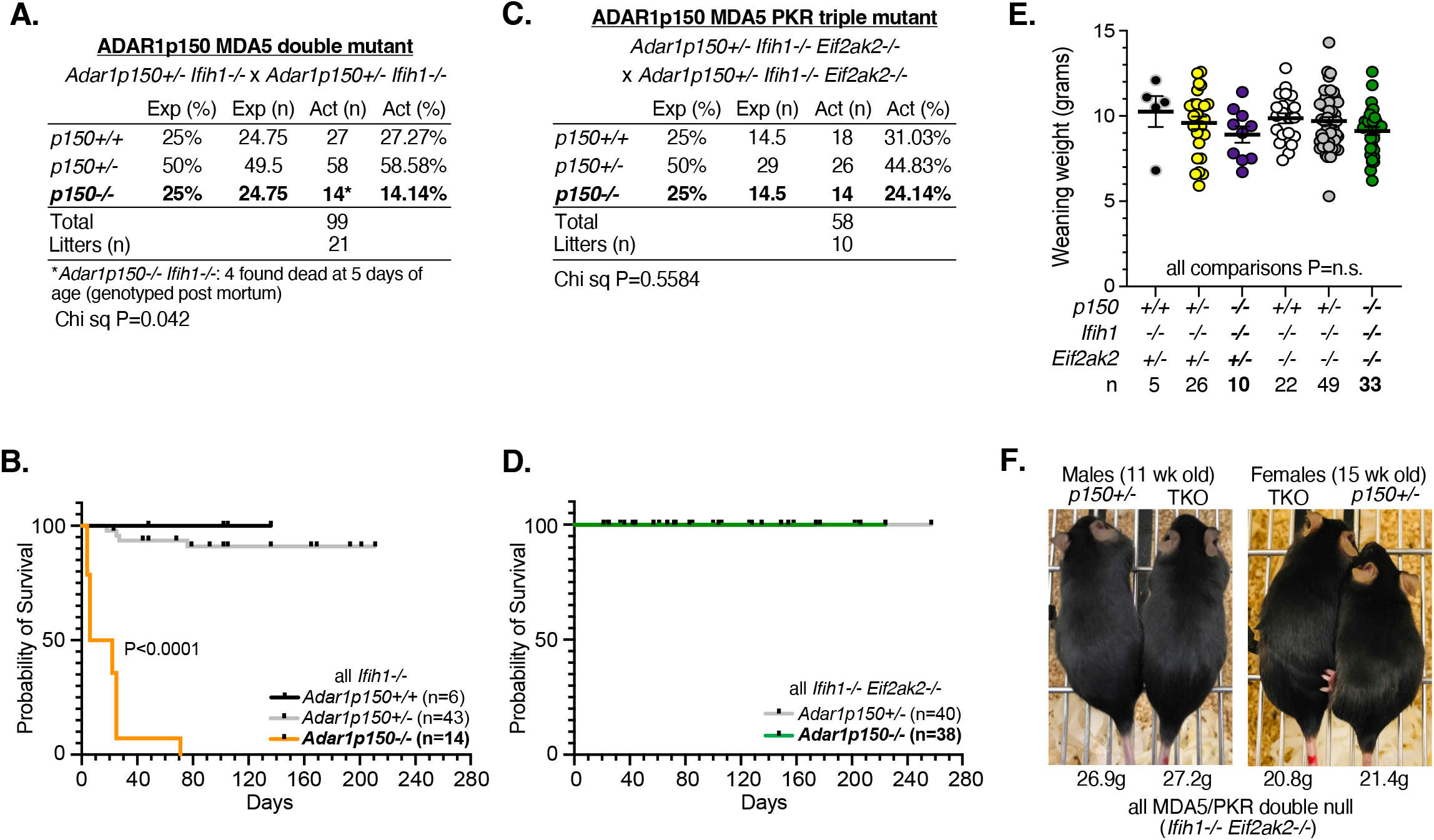
Removal of MDA5 and PKR fully rescues the lethality of *Adar1p150*^-/-^ mice. (A) Breeding data of *Adar1p150*^-/-^*Ifih1*^+/-^ mice. Breeding data from inbreeding of *Adar1p150*^-/-^*Ifih1*^-/-^ breeding pairs. Statistical significance determined with Chi Squared test. (B) Kaplan Meier survival curve of mice with the indicated genotypes. Number of animals as indicated for each genotype. Control mice included littermates with the *Adar1p150*^+/+^ and *Adar1p150*^+/-^ genotype. *P* values were calculated by log-rank test. (C) Breeding data of *Adar1p150*^-/-^*Ifih1*^-/-^*Eif2ak2*^-/-^ mice. Breeding data from inbreeding of *Adar1p150*^+/-^*Ifih1*^-/-^*Eif2ak2*^-/-^ breeding pairs. Statistical significance determined with Chi Squared test. (D) Kaplan Meier survival curve of mice with the indicated genotypes. Number of animals as indicated for each genotype. Control mice included littermates with the *Adar1p150*^+/-^ genotype. *P* values were calculated by log-rank test. (E) Weaning weights of mice of the indicated genotypes irrespective of breeding pair genotype. No statistical difference. (F) Representative photos of male and female sibling *Adar1p150*^-/-^ *Ifih1*^-/-^ *Eif2ak2*^-/-^ and *Adar1p150*^+/-^*Ifih1*^-/-^ *Eif2ak2*^-/-^ (control) at the indicated age.

## Discussion

Through combining human cell lines, genetic screening and *in vivo* murine genetics, we demonstrate that the engagement of cytoplasmic dsRNA sensors downstream of ADAR1 is a species-conserved response determined by both RNA editing and RNA binding activities of the cytoplasmic ADAR1p150. We demonstrate that discrete functions of ADAR1p150 determines MDA5 or PKR engagement: A-to-I editing of endogenous dsRNA specifically prevents MDA5 activation, while PKR activation is prevented by editing-independent, competitive dsRNA binding in the cytoplasm.

The function of PKR as a dsRNA sensor is seemingly well-known but in fact quite elusive. PKR forms dimers on dsRNAs, leading to activation through autophosphorylation and translational shutdown by phosphorylating the translation initiation factor eIF2α^11^. Knockout of PKR does not rescue the *Adar1*^-/-^ mice^30^, which for a long time suggested the lack of physiological role of PKR in *Adar1*^-/-^ mice. This appeared to contradict many *in vitro* observations that PKR is activated in the absence of ADAR1^24,33-43^. We found that in some, but not all, cells IFN treatment is required to observe the PKR/ADAR1 nexus, which helps explain and resolve the discrepancy between previous observations in cells and *in vivo*. Our results confirmed that *in vivo* PKR activation is a significant contributor to the early post-natal death of *Adar1*^-/-^*Ifih1*^-/-^ and *Adar1p150*^-/-^*Ifih1*^-/-^ mice^21,46^. The embryonic lethality of *Adar1*^-/-^*Eif2ak2*^-/-^ mice^30^ is most likely due to the activation of MDA5. Furthermore, the lack of PKR activation in the viable *Adar1^E861A/E861A^Ifih1^-/-^* mice is consistent with the retained expression of an RNA-binding competent, albeit editing deficient, ADAR1 protein^32,56^. Therefore, the understanding arising from our work reconciles ADAR1’s RNA editing dependent and independent functions.

Our work reveals that MDA5 and PKR are the primary, and non-redundant, *in vivo* effectors of organismal lethality following the loss of ADAR1p150. The best evidence is that the *Adar1p150*^-/-^ mice are fully rescued at the Mendelian ratio by the removal of both MDA5 and PKR. Although the dsRBDs of alternative dsRBPs can prevent PKR activation *in vitro*, in the mice lacking ADAR1p150 and MDA5, other dsRBPs do not physiologically compensate and are unable to prevent fatal PKR activation. A simple explanation is that, unlike the overexpressed dsRBDs *in vitro*, the abundance of other dsRBPs *in vivo* is substantially lower than PKR. Future work will be needed to mechanistically understand the role of other dsRBPs in relation to ADAR1p150, MDA5 and PKR *in vivo*.

One dsRBP of recent interest with a role described in ADAR1-dependent pathologies is ZBP1^39,45-47,57-61^. While these studies indicated that the ADAR1p150 Zα domain dampens Z-RNA sensing by ZBP1, the relative contribution of ZBP1, MDA5 and PKR to these phenotypes has not been addressed. Among the genotypic combinations involving ADAR1 deficiency and ZBP1 deletion reported, it is of particular relevance that the loss of ZBP1 increased the survival of *Adar1p150*^-/-^*Ifih1*^-/-^ mice from ~5 days after birth to only 27 days of age^46^. In contrast, we were able to demonstrate the complete rescue of the postnatal lethality of *Adar1p150*^-/-^*Ifih1*^-/-^ mice by PKR removal. This comparison argues for an *in vivo* difference between abrogation of PKR compared to ZBP1. Furthermore, the genetic analyses suggest that ZBP1 acts downstream of MDA5 and PKR activation as one component of lethal autoinflammation.

A question that arises from this work is why A-to-I editing is sufficient to inhibit MDA5 activation but RNA binding by ADAR1p150 is required to prevent PKR activation. One contributing factor may be that the mode of activation of MDA5 and PKR by dsRNA is different. To be activated, MDA5 forms filaments on dsRNAs^62^, with the requirement for long base-paired dsRNA. PKR acts a dimer on dsRNAs, which has been characterized to require only a short length of the double strand structure^63^. This activation mode is consistent with the mechanism by which ADAR1 suppresses MDA5 and PKR, as editing on multiple sites may disrupt the integrity of the double-stranded structure which affects MDA5 filamentation^50,64^, while only competitive binding will block the dimerization of PKR on a relatively short region of dsRNAs. Future work is needed to answer the question of whether the same or different endogenous dsRNA substrates activate MDA5 and PKR *in vivo*. One hypothesis is that the endogenous dsRNAs for MDA5 and PKR activation are different populations. However, ADAR1 is required to prevent cellular dsRNA activating both MDA5 and PKR, suggesting that the endogenous dsRNAs either for MDA5 or PKR activation should be part of ADAR1’s substrates. In addition, the major type of endogenous dsRNAs in HEK293T is derived from IR*Alus*, and the IR*Alus* RNAs have been reported to activate both MDA5^50,64^ and PKR^54,65^. However, this does not preclude the possibility that different pools of ADAR1 substrates may activate MDA5 and PKR independently. The alternative hypothesis is that although PKR and MDA5 share some endogenous dsRNA substrates, they may have different sensitivity to activation by dsRNAs. In support of this, the overexpression of MDA5 in ADAR1^KO^ or ADAR1^E912A^ cells lead to ISG induction directly, indicating that the basal expressed dsRNAs in normal HEK293T are sufficient for MDA5 activation. In contrast, although PKR protein is expressed in the HEK293T cells, deletion of ADAR1 alone did not activate PKR. This suggests MDA5 may be sensitive to lower levels of dsRNA. PKR activation required IFN-induced transcription^24^ which has been reported to induce the production of more dsRNAs^44^. Basal level of endogenous dsRNAs was not sufficient to activate PKR and IFN signaling-induced dsRNAs are required for PKR activation in the absence of ADAR1. Our finding that overexpression of dsRBDs inhibited PKR activation may explain why PKR is less sensitive, as other dsRBDs containing proteins may additionally compete with PKR for dsRNA binding which may contribute to the higher dsRNA threshold for PKR activation.

The distinction between the RNA editing and RNA binding activities of ADAR1p150 will be important for the development of targeted ADAR1 inhibitors. ADAR1 inhibition has emerged as a priority immuno-oncology target^66^, sensitizing tumors to immunotherapy and overcoming resistance to immune checkpoint blockade in mice^33,38,44^. In tumor cell lines the loss of both MDA5 and PKR was required to abrogate the immunotherapy sensitizing effects of ADAR1 loss^44^. This is consistent with the model we propose as these studies utilized ADAR1 protein deficient contexts. For priming an immune response by turning “cold” tumors “hot”, it may be sufficient and preferred to selectively activate MDA5 but not PKR. This could be achieved by transiently inhibiting ADAR1’s RNA editing activity, specifically engaging MDA5 and stimulating type I IFN signaling, but avoiding excessive PKR activation which may be deleterious to normal tissues. On the other hand, inhibiting both editing and binding activities of ADAR1 could be applied to engage a distinct response in tumor cells via activating both MDA5 and PKR. Therefore, the mechanistic understanding from this study lays a foundation for strategies to optimize the development of ADAR1 inhibitors. These data resolve a mechanism and provide direct genetic evidence that both PKR and MDA5 are activated *in vivo* when ADAR1p150 protein is absent and inhibition of both is necessary to abrogate the fatal autoinflammatory response that occurs following the loss of ADAR1p150.

## Acknowledgements

The authors would like to thank Andrew Fire, Siddharth Balachandran, Jan Carette and Brian Liddicoat for comments and discussion, Anthony Sadler (Hudson Institute, Monash University) for providing *Eif2ak2*^-/-^ mice, the Monash Genome Modification Platform (MGMP) at Monash University for the generation of the *Adar1p150*^-/-^ (p.L196C*fs*X6) mice; Monash Antibody Technology Facility (MATF) for purification of ADAR1 antibody from hybridomas; the Phenomics Australia Histopathology and Slide Scanning Service, University of Melbourne for histopathology on samples; St. Vincent’s Hospital Bioresource’s Centre for care of experimental animals; Mark Kamps (UCSD) and Xinshu Xiao (UCLA) for plasmids, Michael Bassik (Stanford) for the CRISPR library, and Addgene for plasmid distribution. The *Adar1p150*^-/-^ mutant mice were produced via CRISPR/Cas9 mediated genome editing by the Monash Genome Modification Platform (MGMP), Monash University as a node of Phenomics Australia. Schematic figures were made using BioRender.com.

This work was supported by: National Institutes of Health, USA (R35GM144100, R01GM124215 and R01GM102484 to JBL); National Health and Medical Research Council Australia (NHMRC; APP1183553 to CRW and JHF; APP1182453 to JHF); 5point Foundation (JHF); University of California, Tobacco-Related Disease Research Program (TRDRP, Award No. T31FT1755; SBH); Melbourne Research Scholarship, The University of Melbourne (ZL); Victorian State Government Operational Infrastructure Support Scheme (to St Vincent’s Institute); Services provided by Phenomics Australia (PA): This study utilized the Phenomics Australia Histopathology and Slide Scanning Service, University of Melbourne and the Monash Genome Modification Platform (MGMP), Monash University. Phenomics Australia is supported by the Australian Government Department of Education through the National Collaborative Research Infrastructure Strategy, the Super Science Initiative, and the Collaborative Research Infrastructure Scheme.

## Author contributions

Conceptualization: SBH, JHF, CRW, JBL; Methodology: SBH, JHF, TS, CRW, JBL; Investigation: SBH, JHF, TS, ZL, AG, ST, CRW, JBL; Visualization: SBH, JHF, CRW, JBL; Funding acquisition: SBH, JHF, CRW, JBL; Project administration: SBH, JHF, CRW, JBL; Supervision: JHF, CRW, JBL; Writing – original draft: SBH, JHF, CRW, JBL; Writing – review & editing: SBH, JHF, TS, ZL, AG, ST, CRW, JBL.

## Declaration of interests

JBL is a co-founder of AIRNA Bio and a consultant for Risen Pharma. All other authors declare that they have no competing interests.

## Methods

### Ethics statement

All animal experiments were approved by the St Vincent’s Hospital Melbourne Animal Ethics Committee (AEC#009/18 and AEC#016/20).

### Plasmid construction and generation of stable cell lines

The ADAR1^KO^ HEK293T cell line and ADAR1^E912A^ HEK293T cell line were described previously^50^. For human PKR knockdown, shRNA sequence, “ATAATAAAGGACTAACTGC”, was inserted into the pMK1200 vector (pMK1200 was a gift from Martin Kampmann & Jonathan Weissman (Addgene plasmid # 84219; http://n2t.net/addgene:84219; RRID: Addgene_84219)). Sequence, “AGCACTCGCATTCGGAGTCAAC”, was inserted into pMK1200 and was used as scramble shRNA. For human MDA5 knockdown, shMDA5 was purchased from Sigma (TRCN0000050849). Overexpression constructs were generated by amplification of CDS of human ADAR1, ADAR2, or dsRNA binding domains from ADAR1, ADAR2, STAU1 and integration into the AscI and PacI sites of the pCDH-puro backbone. Primers used are listed in Supplemental Table S4. The doxycycline-inducible MDA5 construct was generated by replacing dCas9 and mCherry with the human MDA5 CDS and GFP in the HR-TRE3G-dCas9-GCN4-10x-p2a-mCherry backbone. Human ADAR1p150 and its mutants (EAA and E912A) are gifts from Xinshu Xiao (UCLA). The ADAR1^KO^ U937 cell line was a single clone derived from CRISPR-mediated ADAR1 KO. The ADAR1 gRNAs were listed in Supplemental Table S4.

### Cell culture and transfection

HEK293T and U937 cells were maintained in DMEM (Gibco) and RPMI medium 1640 (Gibco), respectively, supplemented with 10% FBS and penicillin/streptomycin. Transfection was carried out with Lipofectamine 2000 (Thermofisher) according to the manufacturer’s protocol, with 80%~90% transfection efficiency in general.

### Lentivirus production and cell infection

To package lentivirus, HEK293FT cells in a 6-cm dish were co-transfected with 4μg viral vectors, 3μg of psPAX2 and 1.2μg pMD2.G. The medium containing viral particles was collected twice at 48 and 72 hr after transfection, filtered through Millex-HV Filter Unit (0.45 μm PVDF, Millipore), and stored at - 80°C until use. Filtered viral containing media was used to infect cells in the presence of 1μg/ml polybrene (Sigma).

### Stress granule staining

Cells were seeded on 18 × 18 mm coverslips (Thermo) and treated with 10ng/ml IFNα (Sigma) for 24 or 48 hrs. Cells were then fixed with 4% PFA for 15 mins at room temperature and permeabilized by 0.5% Triton X-100 for 5 mins on ice. Antibodies: mouse anti-G3BP1 (Abcam, Ab56574) and Alexa Fluor™ 488 goat anti-mouse (Invitrogen, A11001), were used to identify stress granules. Slides were mounted using ProLong™ Gold antifade reagent with DAPI (Invitrogen, P36931). Images were obtained using a LEICA DMRXA2 microscope. Images were analyzed by ImageJ.

### Cell proliferation assay

To monitor cell proliferation of HEK293T cells, 2.5 × 10^5^ WT, ADAR1^E912A^, or ADAR1^KO^ cells were mixed with 2.5 × 10^5^ WT cells that were labelled GFP. Cells were treated with or without 10ng/μl IFNα. The proportion of GFP negative cells was evaluated by flow cytometry (FSC/SSC) using a BD Accuri C6 flow cytometer on days 2, 4, 6, 8, and 10. The relative cell proportion was calculated by normalizing the GFP negative cell proportion of IFNα-treated group to that of the untreated group. To detect cell proliferation of PKR KD cells, scramble or shPKR-treated ADAR1^KO^ cells (mCherry positive) were mixed with an equal number of WT cells. The proportion of mCherry positive cells was evaluated. To assess cell proliferation of MDA5-induced cells, WT cells with inducible MDA5 (GFP positive) were mixed with WT cells, ADAR1^E912A^ cells with inducible MDA5 (GFP positive) were mixed with ADAR1^E912A^ cells, and ADAR1^KO^ cells with inducible MDA5 (GFP positive) were mixed with ADAR1^KO^ cells. The proportion of GFP-positive cells was evaluated as described above.

### CRISPR screening

The human CRISPR-Cas9 Deletion Library was a gift from Michael Bassik (Stanford University). CRISPR screening and subsequent analysis were carried out as described^68^. Approximately 240 × 10^6^ ADAR1^KO^ U937 cells with Cas9 expressed were infected with viral supernatant containing the gRNA library to achieve a >20% infection rate. Infected cells were selected by puromycin (1μg/ml) for 3 days and expanded. Cells were passaged into 4 flasks with ~240 × 10^6^ cells and 480mL media per flask. Cells in two flasks were treated with 10μg/ml IFNα. Cells in the remaining two flasks were untreated and used as the control population. Cells were cultured for 2 weeks and the concentration of cells was maintained at 0.5 × 10^6^ cells/mL. After screening, ~250 × 10^6^ cells from each flask were collected and subjected for genomic DNA extraction using Qiagen’s QIAamp DNA Blood Maxi Kit (Qiagen, Cat # 51194). The integrated gRNAs were amplified (first PCR) and Illumina adapters were added (second PCR) in a nested PCR manner using Agilent Herculase II Fusion DNA Polymerase Kit (Agilent, Cat # 600679). The PCR products were purified and sequenced using an Illumina NextSeq. 50 million reads per sample were achieved. The enrichment of screen candidate genes was analyzed using casTLE^67^. Primers used are listed in Supplemental Table S4.

### Biotin-labeled RNA pull-down

DNA fragments of an *Alu* sequence with T7 promoter on the 5’ end (ss*Alu*) or both ends (ds*Alu*) were *in vitro* transcribed with the biotin RNA-labeling mix (Roche) and T7 transcription kit (Roche). 1 × 10^7^ HEK293T cells were suspended in 1 mL RIP buffer (25mM Tris at pH7.5, 150mM KCl, 0.5mM DTT, 0.5% NP40, 2mM VRC, protease inhibitor cocktail) followed by sonication. The cell lysate was centrifuged at 13,000 rpm for 15 min at 4°C and the supernatant was collected for pre-clearing with 40μl streptavidin beads at 4°C for 40 mins. The precleared lysate was divided into two tubes, and each was supplemented with 2μg of re-natured biotin-labeled *ssAlu* or ds*Alu* followed by incubation at room temperature for 1.5 hrs. Then, 40μL streptavidin Dynabeads (Invitrogen) was added and incubated for another 1.5 hrs. The beads were washed four times for 5 min with RIP buffer containing 0.5% sodium deoxycholate and boiled in 1xLaemmli Sample Buffer (Bio-Rad) for 10 min at 100°C. The retrieved proteins were analyzed by Western blotting. Primers used are listed in Supplemental Table S4.

### Western blot

5 million cells were collected and dissolved with 200μl 1xLaemmli Sample Buffer (Bio-Rad) and heated at 100°C for 15 min. 10μl sample was loaded for PAGE. Following human antibodies were used: ADAR1 (Santa Cruz Biotechnology, sc-73408), Flag (Sigma-Aldrich, F1804), PKR (Cell Signaling Technology, 12297), phosphorylated PKR (phospho T446, Abcam, ab32036), ILF2 (Bethyl laboratories, A303-147A), GAPDH (Santa Cruz Biotechnology, sc-47724), Actin (Abcam, ab8227).

### Real-time PCR

RNA was extracted from 2-5 million HEK293T or U937 cells using TRIzol Reagent (Ambion, 15596018). Genomic DNA was removed using DNA-free DNA removal Kit (ThermoFisher, AM1906). Then, 1μg RNA was reverse transcribed using iScript Advanced cDNA Synthesis Kit (Bio-Rad, 1725073). Realtime PCR was run on the Bio-Rad CFX96 with KapaFast qPCR mix. Primers are listed in Supplemental Table S4.

### U937 differentiation

The U937 differentiation was carried out as described in^51^. 50nM phorbol myristate acetate (PMA) was added to 1 million per mL U937 cells in U937 growth medium. The cells were incubated for 3 days for differentiation. After that, the cells were suspended by Trypsin and collected for analysis of RNA or protein expression, or passaged for proliferation observation.

### Mouse lines

*Adar^E8681A/+^* (*Adar1*^*E861A*/+^; MGI allele: Adar^tm1.1Xen^; MGI:5805648), *Ifih1*^-/-^ (Ifih1 ^tm1.1Cln^), *Adar1*^-/-^ (Adar1^-/-^; MGI allele: Adar^tm2Phs^; MGI:3029862), *Adar^fl/fl^* (*Adar1^fl/fl^*; exon 7-9 floxed; MGI allele: Adar^tm1.1Phs^; MGI:3828307), *Eif2ak2*^-/-^ (Eif2ak2^tm1Cwe^; *Pkr*^-/-^; MGI:2182566 generously provided by Dr A Sadler; Hudson Institute of Medical Research, Clayton, Australia)^69^ and *Rosa26*-CreER^T2^(Gt(*ROSA*)26^Sortm1(cre/ERT2)Tyj^) mice. *Adar1p150*^-/-^ mice (nomenclature based on NCBI CCDS50963; nucleotide 587delT, p.L196C*fs*X6) were identified as an incidental mutation when using CRISPR/Cas9 targeting in C57BL/6 zygotes by the Monash Genome Modification Platform (Monash University, Clayton, Australia). Details of the allele and phenotype have been posted (https://doi.org/10.1101/2022.08.31.506069). Introduction of the mutation was confirmed by Sanger sequencing of the region in both the founders and subsequent generations. *Adar1p150*^-/-^ mice were genotyped by PCR followed by Sanger sequencing. All mice were on a backcrossed C57BL/6 background as previously described^19,32,70^. For day of birth analysis, females were plug mated and pups collected before midday on the day of birth. For genetic rescue experiments all genotypes were assessed at day 7-10 of age; with any pups found dead genotyped post-mortem. All weaning weights were recorded by the animal facility staff, other weights or parameters were recorded by the investigators. Genotyping primers provided in Supplemental Table S4.

### Mouse tail fibroblasts

Tails were collected from animals of the indicated genotypes at the day of birth to generate fibroblasts. Tail pieces were rinsed in 70% ethanol and briefly air-dried before mincing with a scalpel into small pieces. These were incubated for 20 minutes in 100uL of 0.025% Trypsin-EDTA (Gibco/Thermo Fisher) at 37°C in a 6-well plate and then 4mL of media was added (High glucose DMEM (Sigma), 10% FBS, 1%Penicillin/Streptomycin, 1% glutamax, 1% non-essential amino acids (Gibco/Thermo Fisher)). The tissue pieces were incubated in a hypoxia chamber flushed with 5% oxygen/5% carbon dioxide in nitrogen at 37°C overnight. The next day tissue chunks were further dissociated by pipetting up and down. Once clusters of cells grew out, cells were trypsinized and expanded onto 10cm plates for interferon treatment. Tail fibroblasts on 10cm plates were treated with recombinant murine interferon beta (PBL Assay Science; PBL-12405) at 250U/mL for 24 hrs in normal growth media. After 24 hours, cells were collected by trypsinization and pellets washed in cold PBS and resuspended in RIPA buffer (20mM Tris·HCl, pH8.0, 150mM NaCl, 1mM EDTA, 1% sodium deoxycholate, 1% Triton X-100, 0.1% SDS) supplemented with 1× HALT protease inhibitor and 1× PhosSTOP phosphatase inhibitor (Thermo).

### Western blot from mouse tissue

Frozen tissue samples were homogenized in 350μL of ice-cold RIPA buffer (20mM Tris·HCl, pH8.0, 150mM NaCl, 1mM EDTA, 1% sodium deoxycholate, 1% Triton X-100, 0.1% SDS) supplemented with 1x HALT protease inhibitor and 1x PhosSTOP phosphatase inhibitor (Thermo) with a mechanical homogenizer (IKA T10 basic S5 Ultra-turrax Disperser). Homogenized tissue samples were freeze-thawed and centrifuged at 13,000g for 20 minutes at 4°C and the supernatant retained for quantification and Western analysis. Protein lysates from immortalized murine myeloid cells were generated by resuspending 2 × 10^6^ washed cells in 100uL of sample lysis buffer (RIPA buffer, 5 x HALT protease inhibitor, 1 × PhosSTOP, 1 × NuPage LDS Sample and reducing buffer) and heating at 70°C for 10 minutes. Western blot using the following antibodies: ADAR1 (Rat monoclonal anti-mouse ADAR1, clone RD4B11, in-house -^19^), PKR (Abcam, EPR19374), phospho-eIF2α (Anti-EIF2S1 (phospho S51), Abcam, ab32157), total eIF2α (Cell Signalling Technology, 5324), MDA5 (Invitrogen, 33H12L34), Panactin (Sigma, MS-1295-PO) and secondary antibodies goat anti-rabbit HRP (Thermofisher, 31460), goat anti-mouse HRP (Thermofisher, 31444) goat anti-rat HRP (Thermofisher, 31470). Western bands from non-saturating exposures to film were quantified using Fiji image analysis. Phospho-eIF2α was normalised to total eIF2α.

### Real-time PCR from mouse tissues

Mouse tissues from the indicated genotypes were isolated from the pups collected at the day of birth and snap-frozen in liquid nitrogen. The tissues were homogenized in Trisure reagent using IKA T10 basic S5 Ultra-turrax Disperser. RNA was then extracted using Direct-Zol columns (Zymo Research) using the manufacturer’s instructions, followed by DNase I digest (Bioline) and clean up (Zymo clean and concentrate kit). Complementary DNA (cDNA) was synthesized using Tetro cDNA synthesis kit (Bioline). Real-time PCR was done in duplicate with Brilliant II SYBR Green QPCR Master Mix (Agilent Technologies) and primers from IDTDna. All primers were optimized to have equal efficiency (100 +/- 10%) before use. *Ppia* was used as a reference gene for relative quantification using the ΔΔCt method. Primer sequences are provided in Supplemental Table S4.

### Statistical analysis

For biological experiments, the significance of results was analyzed using the Student’s *t*-test, one-way or two-way ANOVA with multiple comparison corrections unless otherwise stated; Survival data was assessed using Kaplan Meier plots and Log-rank (Mantel-Cox) test using GraphPad Prism; For analysis of breeding data Chi-squared tests in GraphPad Prism were used to determine Mendelian ratios of offspring *P* <0.05 was considered significant. All data are presented as mean ± SEM.

### Data and materials availability

All data presented in this manuscript are available from the corresponding author upon reasonable request. Animal models are available subject to completion of materials transfer agreements (MTAs). RNA sequencing data were deposited at the Gene Expression Omnibus (GEO) under accession number GSE198386.

## Supplemental information

**Supplemental Figures 1-7**

**Supplemental Tables**

**Table S1.** Gene expression in WT, ADAR1^E912A^, and ADAR1^KO^ HEK293T cells shown as Transcripts Per Million, related to Figure 2.

**Table S2.** CasTLE analysis of CRISPR KO screening, related to Figure 2.

**Table S3.** Gene expression in WT and ADAR1^KO^ U937 cells with or without PMA, shown as Transcripts Per Million, related to Figure 4.

**Table S4.** List of DNA oligos, related to Methods.

## Supplemental Data

**Data S1.** Replicates of Western blot, related to Figures 2-6.

**Data S2**. Histological analysis reports of *Adar1*^-/-^*Ifih1*^-/-^*Eif2ak2*^-/-^ and *Adar1p150*^-/-^*Ifih1*^-/-^*Eif2ak2*^-/-^ mice, related to Figure 7.

## Notes

### Competing Interest Statement

Jin Billy Li is a co-founder of AIRNA Bio and a consultant for Risen Pharma. All other authors declare that they have no competing interests.

