## Supplemental Figures for "ADAR1p150 Prevents MDA5 and PKR Activation via Distinct Mechanisms to Avert Fatal Autoinflammation"

**Supplemental figure legends**

**Figure S1. Breeding data from *Adar1*$^{-/-}$ mice (Related to Figure 1)**

(A) RT-qPCR of RNA isolated from day of birth mouse kidneys for the indicated target ISR and ISG genes relative to *Ppia*. Data was summarized from 3 biological replicates represented as fold change compared to *Adar1*$^{+/+}$*Ifih1*$^{-/-}$ mice (n=5). Significance was determined using a 2-way ANOVA.

(B)-(C) Full data of RT-qPCR of ISR (B) and ISG (C) shown in (A).

(D)-(E) Recovery of pups of each genotype at the day of birth from *Adar1*$^{+/-}$*Ifih1*$^{-/-}$ inbreeding (D) and *Adar1*$^{+/-}$*Ifih1*$^{-/-}$*Eif2ak2*$^{-/-}$ inbreeding (E).

(F) Western blot of murine tail fibroblasts generated from day of birth mice of the indicated genotypes treated for 24 hours with mouse IFNβ or vehicle. IFNβ-treated Kusa4b10 cells (murine mesenchymal cell line) were used as a control for the MDA5 antibody.

(G) Photographs of all triple mutant mice with littermate control at indicated ages.

(H) Body weight of mice of indicated sex and genotype; weight indicated at two ages (in days of age) for each animal. Control animals are littermates; Control for F#169 weight is a *Adar1*$^{+/-}$*Ifih1*$^{-/-}$*Eif2ak2*$^{-/-}$ (het) littermate as the WT littermate was euthanized with malocclusion prior to final weight being able to be recorded.

**Figure S2. Breeding results from *Adar1*$^{E861A/E861A}$ mice (Related to Figure 1)**

(A)-(B) Recovery of pups of each genotype at the day of birth (A) or one week after birth (B) from *Adar1*$^{E861A/+}$*Ifih1*$^{-/-}$ inbreeding. Statistical significance determined with Chi Squared test.

(C) Recovery of pups of each genotype one week after birth from *Adar1*$^{E861A/+}$*Eif2ak2*$^{-/-}$ inbreeding. Statistical significance determined with Chi Squared test.

(D)-(E) Recovery of pups of each genotype at the day of birth (D) or one week after birth (E) from *Adar1*$^{E861A/+}$*Ifih1*$^{-/-}$*Eif2ak2*$^{-/-}$ inbreeding. Statistical significance determined with Chi Squared test.

(F) Kaplan Meier survival curve of mice with the indicated genotypes. Number of animals as indicated for each genotype. Control mice included littermates with the *Adar1*$^{E861A/+}$genotype.

(G) Weaning weights of sibling *Adar1*$^{E861A/E861A}$*Ifih1*$^{-/-}$*Eif2ak2*$^{-/-}$ and *Adar1*$^{E861A/+}$*Ifih1*$^{-/-}$*Eif2ak2*$^{-/-}$ (control) mice.

(H) Representative photos of male and female sibling *Adar1*$^{E861A/E861A}$*Ifih1*$^{-/-}$*Eif2ak2*$^{-/-}$ and control mice at the indicated age.

**Figure S3. Validation of ADAR1 deficiency in HEK293T and U937 (Related to Figure 2)**

(A) Analysis of differentially expressed genes between WT and ADAR1$^{E912A}$ or ADAR1$^{KO}$ HEK293T. ISGs genes are in red.

(B) Schematic work flow for the cell proliferation assay in Figure 2A. Bottom, representative FACS plot and gating for GFP negative and GFP positive populations.

(C) Stress granule staining in ADAR1$^{E912A}$ HEK293T cells. Cells were treated with 0.5mM sodium arsenite for 1hour.

(D) ADAR1 knockout efficiency in U937 cells.

(E) The effect of IFNα treatment on the cell proliferation of WT and ADAR1$^{KO}$ U937 cells. Cell numbers were counted at indicated time points. Data from 3 replicates are shown as the mean +/- SEM. Significance determined by 2-way ANOVA with multiple comparisons.

(F) Schematic outline of the genome-wide CRISPR/Cas9 knockout screen in the ADAR1$^{KO}$ U937 cells.

**Figure S4. ADAR1 deficiency in U937-differentiated macrophage (Related to Figure 4)**

(A) A schematic drawing showed the work flow of U937 differentiation.

(B) Expression comparison of ISGs between U937 and U937-differentiated macrophage in WT (left) and ADAR1$^{KO}$ (right). The gene expression levels were measured by TPM (Transcripts Per Kilobase Million) and log2 transformed.

(C) MDA5 or PKR depletion in U937 ADAR1$^{KO}$ cells using shRNAs.

**Figure S5. Localization of over-expressed proteins in HEK293T cells (Related to Figure 5)**

(A) Localization of flag-tagged exogenous ADAR proteins.

**Figure S6. Loss of ADAR1 decreases PKR's sensitivity to dsRNAs (Related to Figure 6)**

(A) PKR sensitivity to polyI:polyC (polyI:C) in the presence or absence of ADAR1. Different amounts of polyI:C was transfected into WT or ADAR1$^{KO}$ HEK293T cells for 6 hours followed by Western blot analysis of PKR phosphorylation.

(B) PKR sensitivity to dsRNA in the presence or absence of ADAR1. Different amount of *in vitro* transcribed ds*Alu* was transfected in WT or ADAR1$^{KO}$ HEK293T cells for 6 hours followed by Western blot analysis of PKR phosphorylation.

**Figure S7. Breeding data from *Adar1p150*$^{-/-}$ mice (Related to Figure 7)**

(A) Genotype recovery from inbreeding of *Adar1p150*$^{-/-}$*Ifih1*$^{-/-}$*Eif2ak2*$^{-/-}$ (triple mutant) x *Adar1p150*$^{+/-}$*Ifih1*$^{-/-}$*Eif2ak2*$^{+/-}$ breeding.

(B) Body weight of cohorts of the indicated genotypes for males (M) and females (F). Mice age ranged between 62 and 149 days of age when weighed, not paired. Significance was determined by one-way ANOVA with multiple comparisons.

(C) Breeding data of pups bred only from *Adar1p150*$^{+/-}$*Ifih1*$^{-/-}$*Eif2ak2*$^{+/-}$ breeding pairs (Adar1p150 and PKR heterozygous). Statistical significance determined with Chi Squared test.

(D) Kaplan Meier survival curve of all animals of the indicated genotype.

### Figure S1

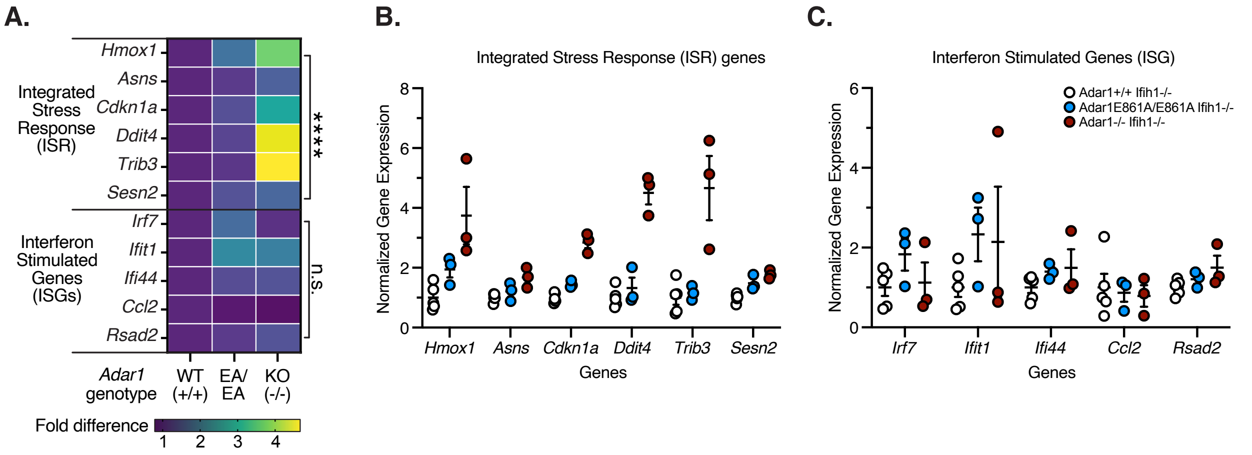


D.

**ADAR1 MDA5 double mutant**

*Adar1+/- Ifih1-/-* x *Adar1+/- Ifih1-/-*

| Day of birth | Exp (%) | Exp (n) | Act (n) | Act (%) |
|---|---|---|---|---|
| *Adar1+/+* | 25% | 12.25 | 14 | 28.57% |
| *Adar1+/-* | 50% | 24.5 | 24 | 48.98% |
| ***Adar1-/-*** | 25% | 12.25 | 11 | 22.45% |
| Total | | | 49 | |
| Litters (n) | | | 8 | |

Chi sq P=0.8238

E.

**ADAR1 MDA5 PKR triple mutant**

*Adar1+/- Ifih1-/- Eif2ak2-/-* x *Adar1+/- Ifih1-/- Eif2ak2-/-*

| Day of birth | Exp (%) | Exp (n) | Act (n) | Act (%) |
|---|---|---|---|---|
| *Adar1+/+* | 25% | 18.75 | 28 | 37.33% |
| *Adar1+/-* | 50% | 37.5 | 37 | 49.33% |
| ***Adar1-/-*** | 25% | 18.75 | 10 | 13.33% |
| Total | | | 75 | |
| Litters (n) | | | 15 | |

Chi sq P=0.0132

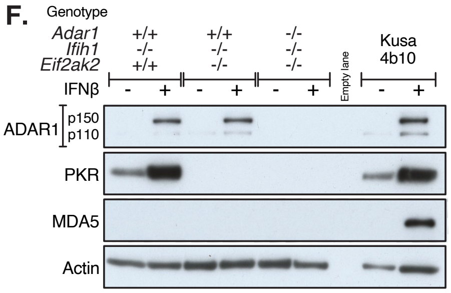


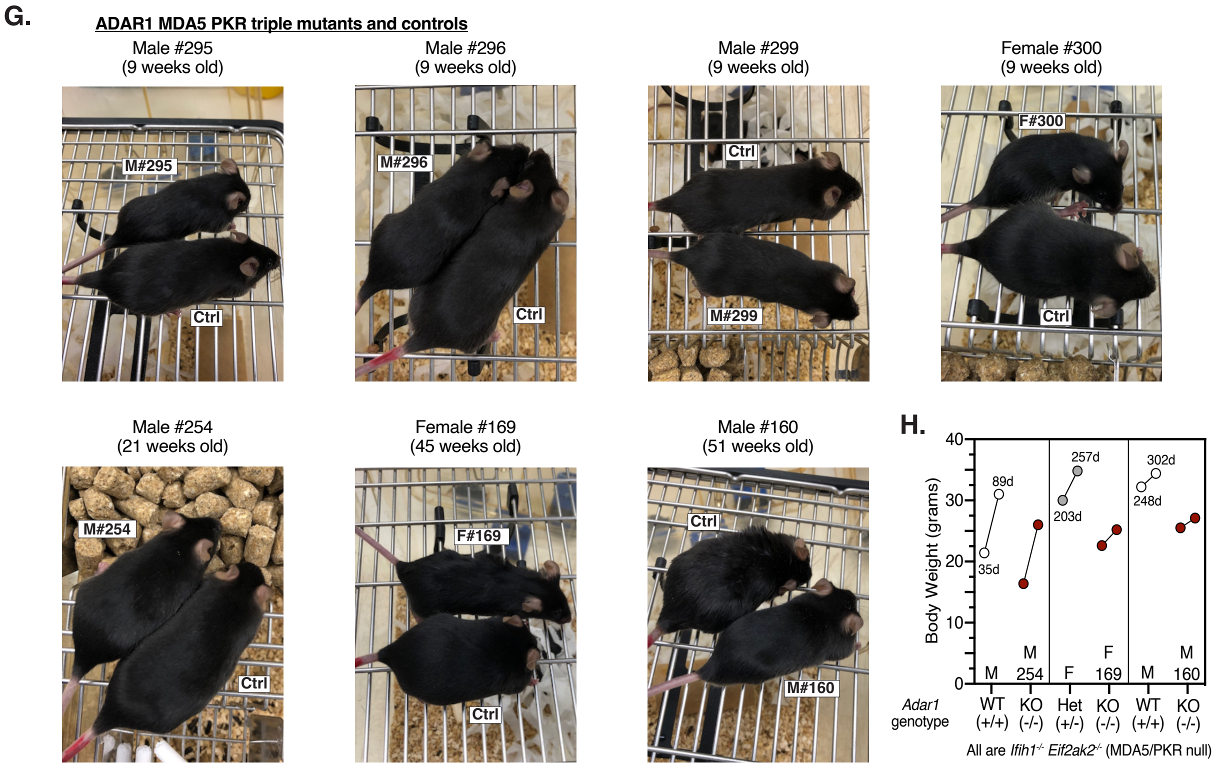

### Figure S2

**A.** **ADAR1 E861A MDA5 double mutant**

*Adar1E861A/+ Ifih1-/-* x *Adar1E861A/+ Ifih1-/-*

Day of birth

| | Exp (%) | Exp (n) | Act (n) | Act (%) |
|---|---|---|---|---|
| *Adar1+/+* | 25% | 13 | 16 | 30.77% |
| *Adar1+/-* | 50% | 26 | 23 | 44.23% |
| ***Adar1ki/ki*** | 25% | 13 | 13 | 25.00% |
| Total | | | 52 | |
| Litters (n) | | | 7 | |

Chi sq P=0.595

**B.** **ADAR1 E861A MDA5 double mutant**

*Adar1E861A/+ Ifih1-/-* x *Adar1E861A/+ Ifih1-/-*

Day 7-10

| | Exp (%) | Exp (n) | Act (n) | Act (%) |
|---|---|---|---|---|
| *Adar1+/+* | 25% | 123.75 | 121 | 34.44% |
| *Adar1+/-* | 50% | 247.5 | 287 | 57.98% |
| ***Adar1ki/ki*** | 25% | 123.75 | 87 | 17.58% |
| Total | | | 495 | |
| Litters (n) | | | 112 | |

Chi sq P=0.0002

**C** **ADAR1 E861A PKR double mutant**

*Adar1E861A/+ Eif2ak2-/-* x *Adar1E861A/+ Eif2ak2-/-*

Day 7-10

| | Exp (%) | Exp (n) | Act (n) | Act (%) |
|---|---|---|---|---|
| *Adar1+/+* | 25% | 29 | 29 | 37.07% |
| *Adar1+/-* | 50% | 58 | 58 | 62.93% |
| ***Adar1ki/ki*** | 25% | 29 | 0 | 0% |
| Total | | | 87 | |
| Litters (n) | | | 26 | |

Chi sq P=0.0001

**D.** **ADAR1 E861A MDA5 PKR triple mutant**

*Adar1E861A/+ Ifih1-/- Eif2ak2-/-* x *Adar1E861A/+ Ifih1-/- Eif2ak2-/-*

Day of birth

| | Exp (%) | Exp (n) | Act (n) | Act (%) |
|---|---|---|---|---|
| *Adar1+/+* | 25% | 13.75 | 14 | 25.45% |
| *Adar1+/-* | 50% | 27.5 | 29 | 52.73% |
| ***Adar1ki/ki*** | 25% | 13.75 | 12 | 21.82% |
| Total | | | 42 | |
| Litters (n) | | | 10 | |

Chi sq P=0.8568

**E.** **ADAR1 E861A MDA5 PKR triple mutant**

*Adar1E861A/+ Ifih1-/- Eif2ak2-/-* x *Adar1E861A/+ Ifih1-/- Eif2ak2-/-*

Day 7-10

| | Exp (%) | Exp (n) | Act (n) | Act (%) |
|---|---|---|---|---|
| *Adar1+/+* | 25% | 13 | 11 | 21.15% |
| *Adar1+/-* | 50% | 26 | 29 | 55.77% |
| ***Adar1ki/ki*** | 25% | 13 | 12 | 23.07% |
| Total | | | 52 | |
| Litters (n) | | | 13 | |

Chi sq P=0.6939

**F.**

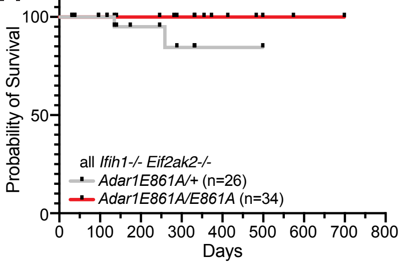


**G.** **ADAR1 E861A MDA5 PKR triple mutant**

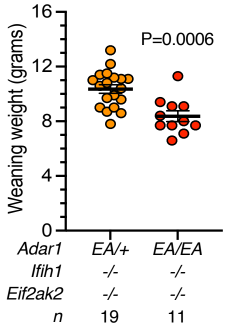


**H.** **ADAR1 E861A MDA5 PKR triple mutant**

Males (13 wk old) Females (20 wk old)

*E861A/+* TKO *Adar1+/+* TKO

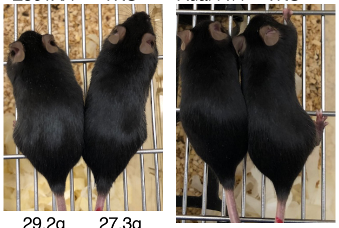

29.2g 27.3g 24.8g 22.3g

**Figure S3**

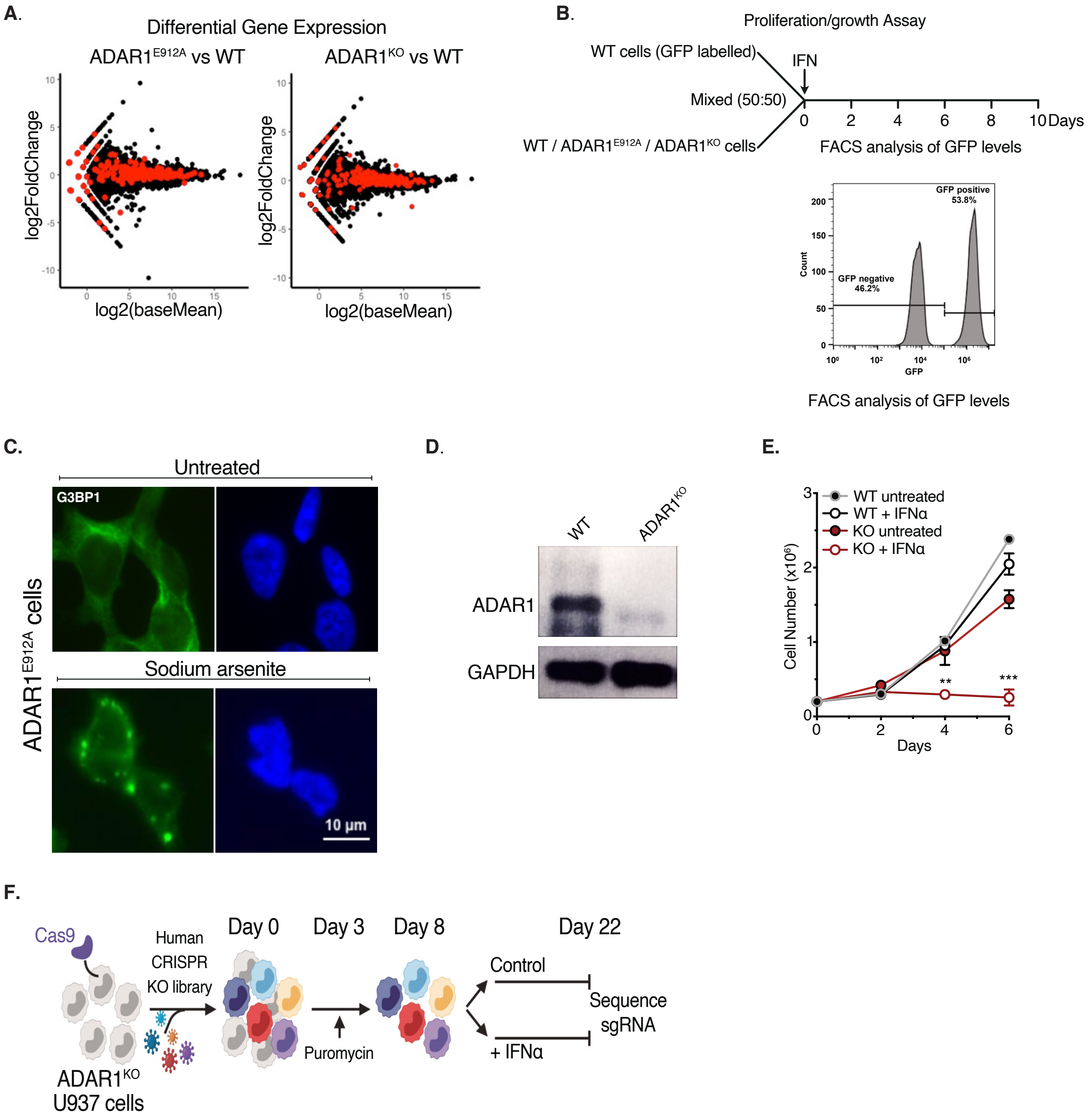
A.
Differential Gene Expression
ADAR1E912A vs WT
ADAR1KO vs WT
log2FoldChange
log2(baseMean)
B.
Proliferation/growth Assay
WT cells (GFP labelled)
IFN
Mixed (50:50)
WT / ADAR1E912A / ADAR1KO cells
0 2 4 6 8 10 Days
FACS analysis of GFP levels
GFP positive 53.8%
GFP negative 46.2%
Count
GFP
FACS analysis of GFP levels
C.
Untreated
G3BP1
Sodium arsenite
ADAR1E912A cells
10 µm
D.
WT
ADAR1KO
ADAR1
GAPDH
E.
WT untreated
WT + IFNα
KO untreated
KO + IFNα
Cell Number (x10⁶)
Days
F.
Cas9
Human CRISPR KO library
ADAR1KO U937 cells
Day 0
Day 3
Day 8
Day 22
Puromycin
Control
+ IFNα
Sequence sgRNA

**Figure S4**

**A.**

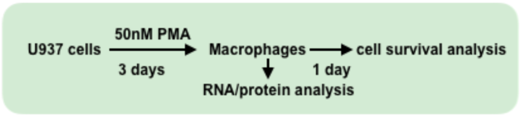


**B.**

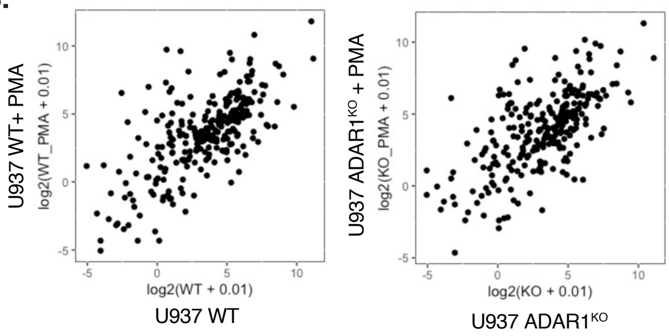


**C.**

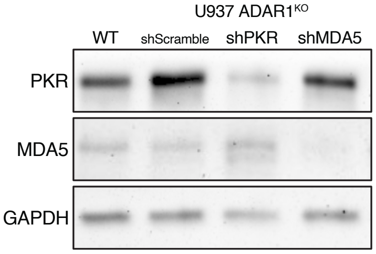

**Figure S5**

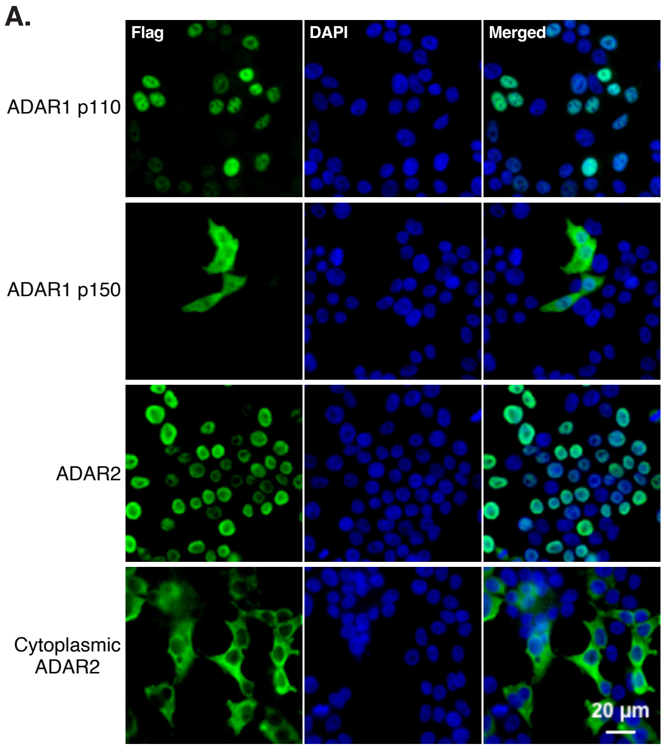
A.
Flag
DAPI
Merged
ADAR1 p110
ADAR1 p150
ADAR2
Cytoplasmic ADAR2
20 µm

**Figure S6**

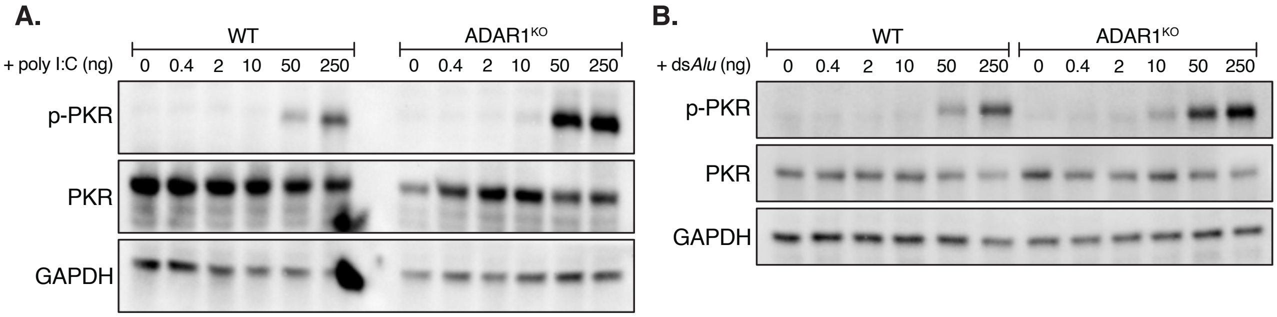
A.
WT
ADAR1KO
+ poly I:C (ng)
0 0.4 2 10 50 250 0 0.4 2 10 50 250
p-PKR
PKR
GAPDH
B.
WT
ADAR1KO
+ dsAlu (ng)
0 0.4 2 10 50 250 0 0.4 2 10 50 250
p-PKR
PKR
GAPDH

#### Figure S7

**A.**

**ADAR1p150 MDA5 PKR triple mutant breeding**

*Adar1p150-/- Ifih1-/- Eif2ak2-/-* (triple mutant) x
*Adar1p150+/- Ifih1-/- Eif2ak2+/-*

| | Exp (%) | Exp (n) | Act (n) | Act (%) |
|---|---|---|---|---|
| *p150+/-Ifih1-/- Eif2ak2+/-* | 25% | 3.75 | 5 | 33.33% |
| *p150+/- Ifih1-/- Eif2ak2-/-* | 25% | 3.75 | 4 | 26.67% |
| ***p150-/- Ifih1-/- Eif2ak2+/-*** | **25%** | **3.75** | **1** | **6.67%** |
| ***p150-/- Ifih1-/- Eif2ak2-/-*** | **25%** | **3.75** | **5** | **33.33%** |
| Total | | | 15 | |
| Litters (n) | | | 3 | |

**C.**

**ADAR1p150 MDA5 PKR+/- triple mutant**

*Adar1p150+/- Ifih1-/- Eif2ak2+/-* x
*Adar1p150+/- Ifih1-/- Eif2ak2+/-*

| | Exp (%) | Exp (n) | Act (n) | Act (%) |
|---|---|---|---|---|
| *p150+/+ Eif2ak2+/-* | 12.5% | 5.75 | 4 | 8.70% |
| *p150+/- Eif2ak2+/-* | 25% | 11.5 | 18 | 39.13% |
| ***p150-/- Eif2ak2+/-*** | **12.5%** | **5.75** | **8** | **17.39%** |
| *p150+/+ Eif2ak2-/-* | 12.5% | 5.75 | 5 | 10.87% |
| *p150+/- Eif2ak2-/-* | 25% | 11.5 | 5 | 10.87% |
| ***p150-/- Eif2ak2-/-*** | **12.5%** | **5.75** | **6** | **13.04%** |
| Total | | | 46 | |
| Litters (n) | | | 9 | |

Chi sq P=0.1144; all *Ifih1-/-*

**B.**

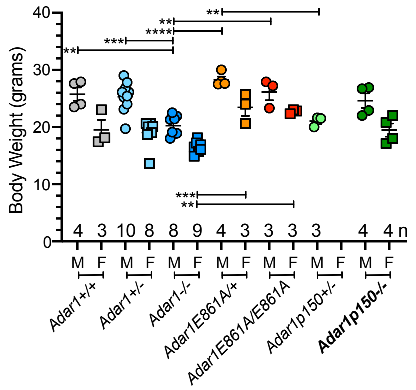


all *Ifih1-/- Eif2ak2-/-*

**D.**

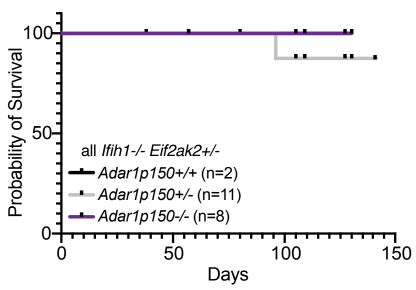
