## Supplemental Data S1 for "ADAR1p150 Prevents MDA5 and PKR Activation via Distinct Mechanisms to Avert Fatal Autoinflammation"

### Figure 2B

##### Replicate 1 (Presented)

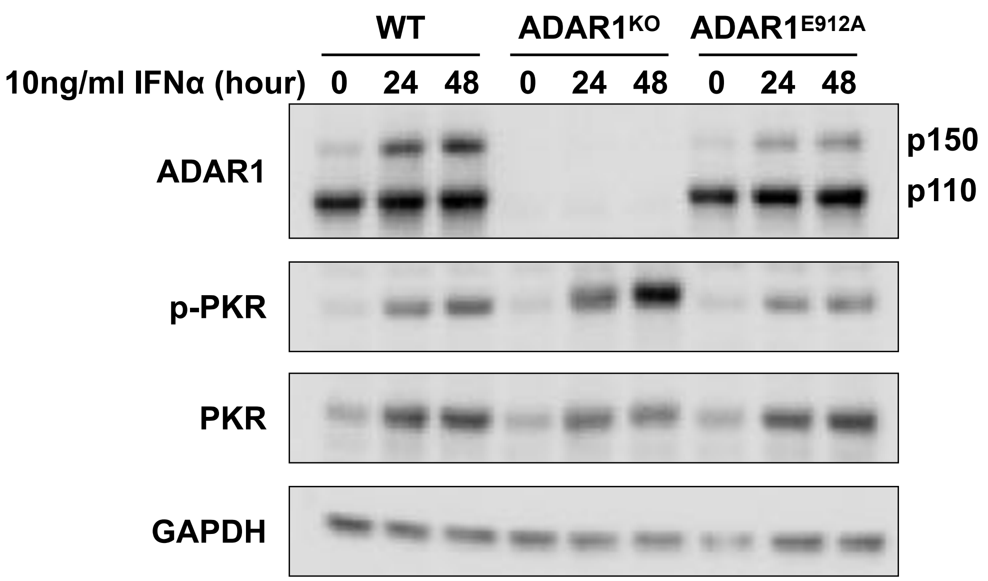


##### Replicate 2

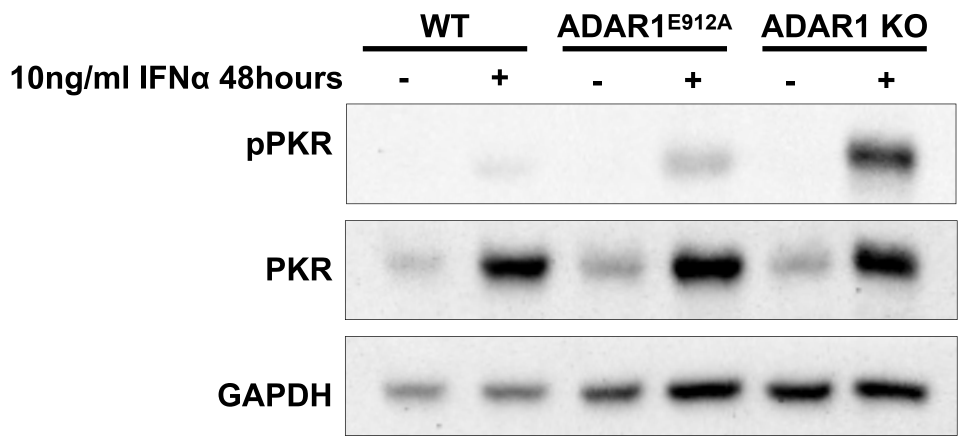


##### Replicate 3

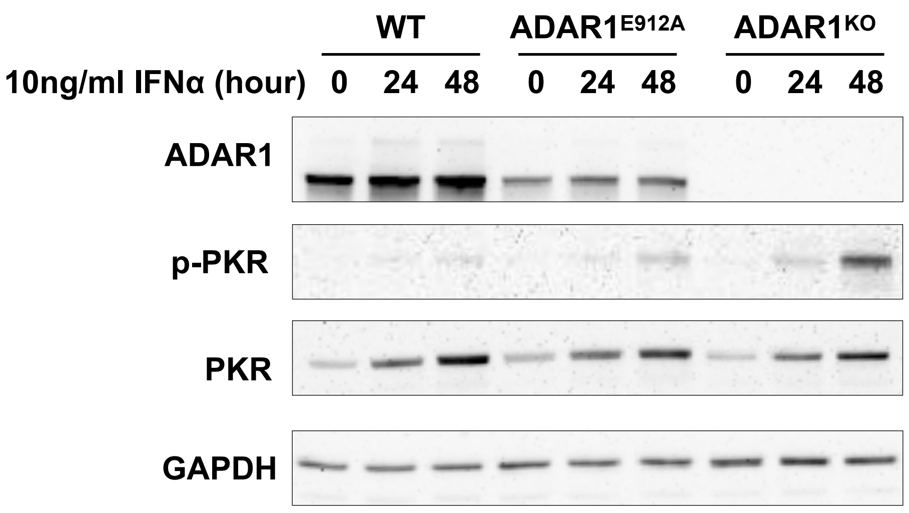

### Figure 2D

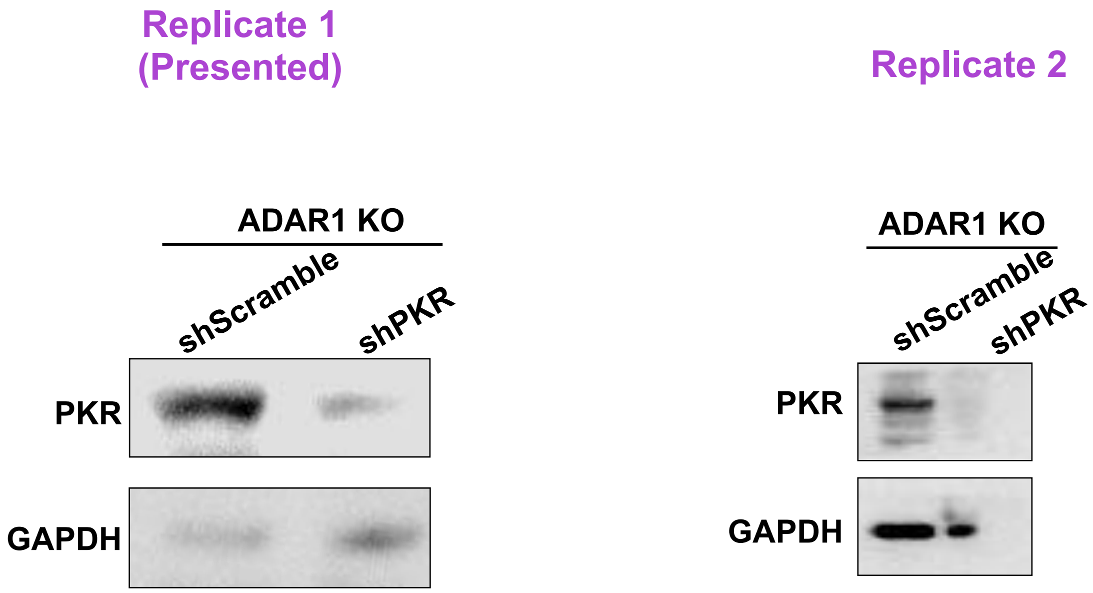
Replicate 1
(Presented)
ADAR1 KO
shScramble
shPKR
PKR
GAPDH
Replicate 2
ADAR1 KO
shScramble
shPKR
PKR
GAPDH

### Figure 3C

Replicate 1
(Presented)

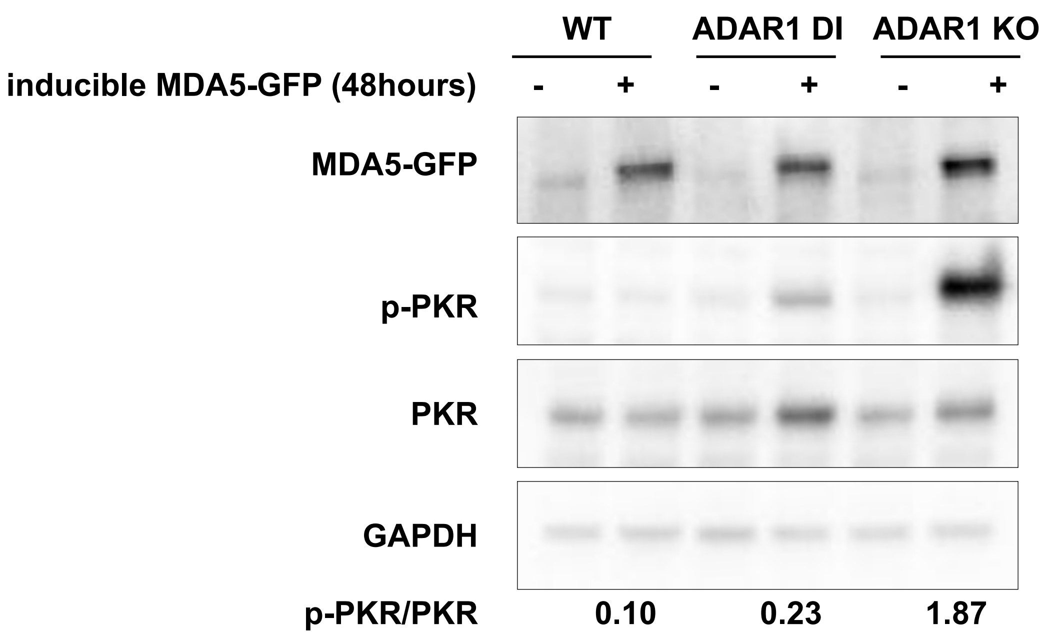


Replicate 2

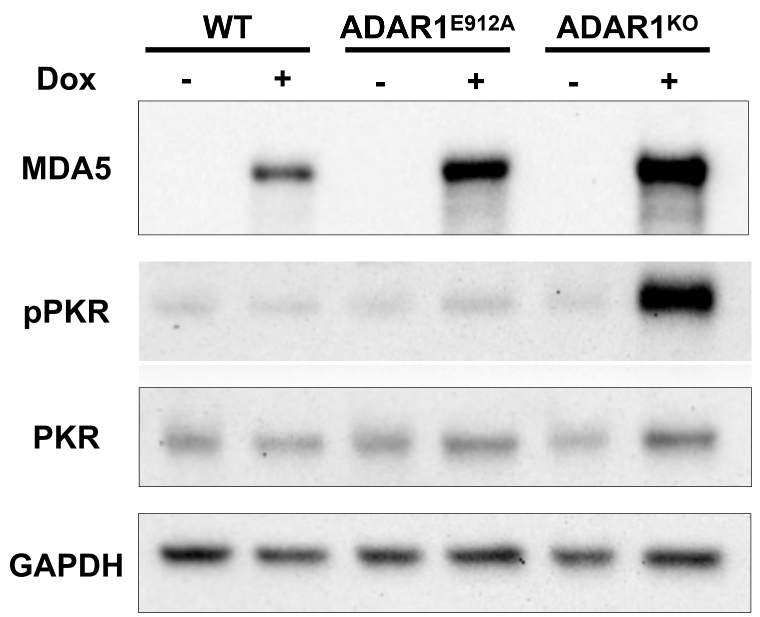

### Figure 3F

Replicate 1
(Presented)

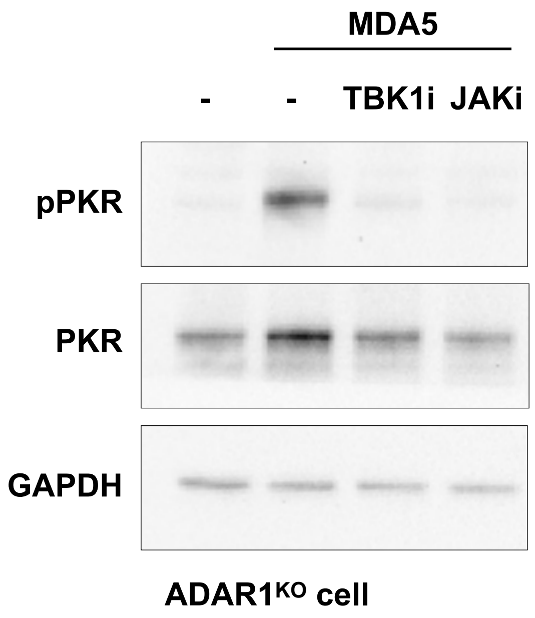


Replicate 2

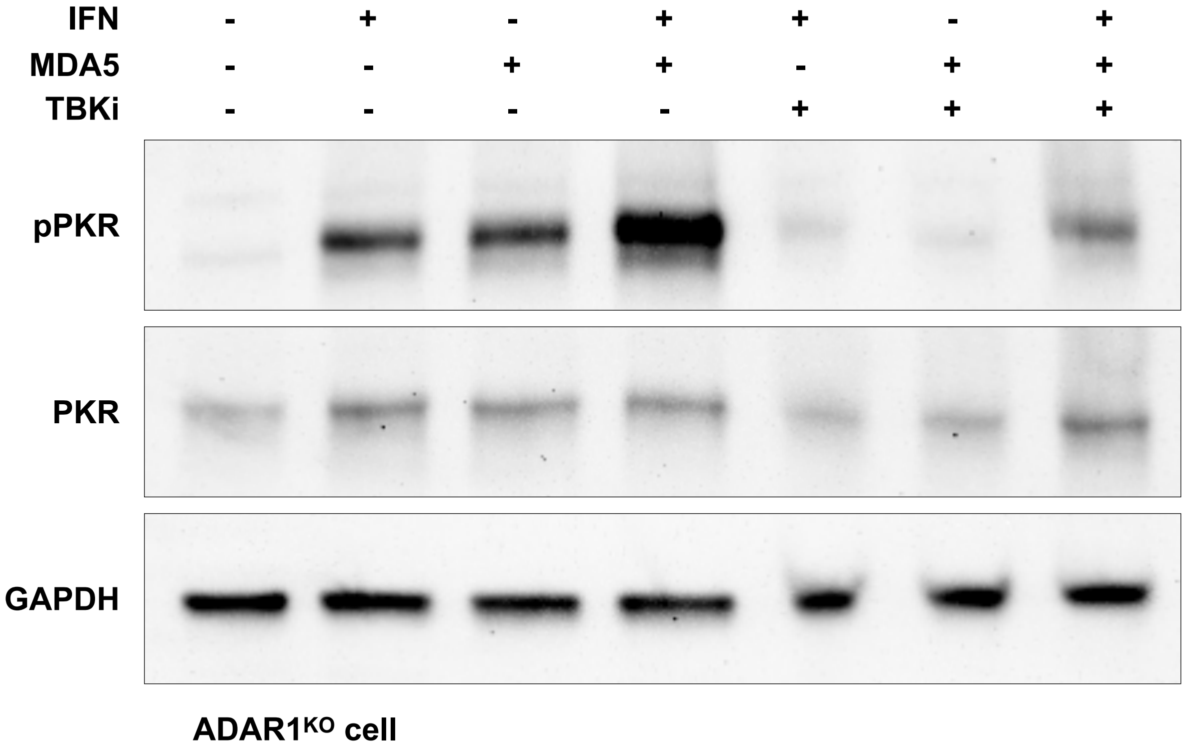

### Figure 3F

Replicate 1
(Presented)

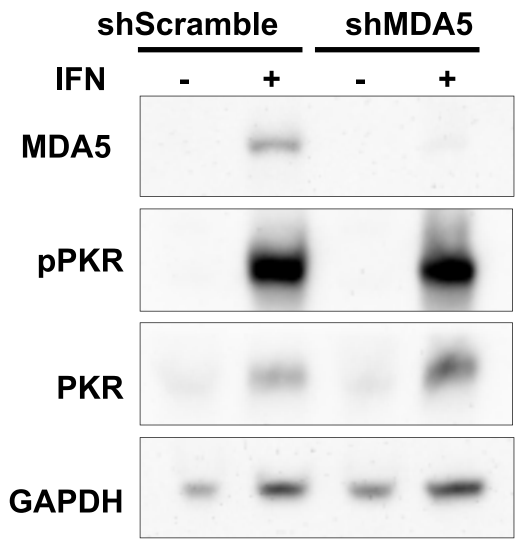


ADAR1$^{KO}$ cells

### Figure 4C

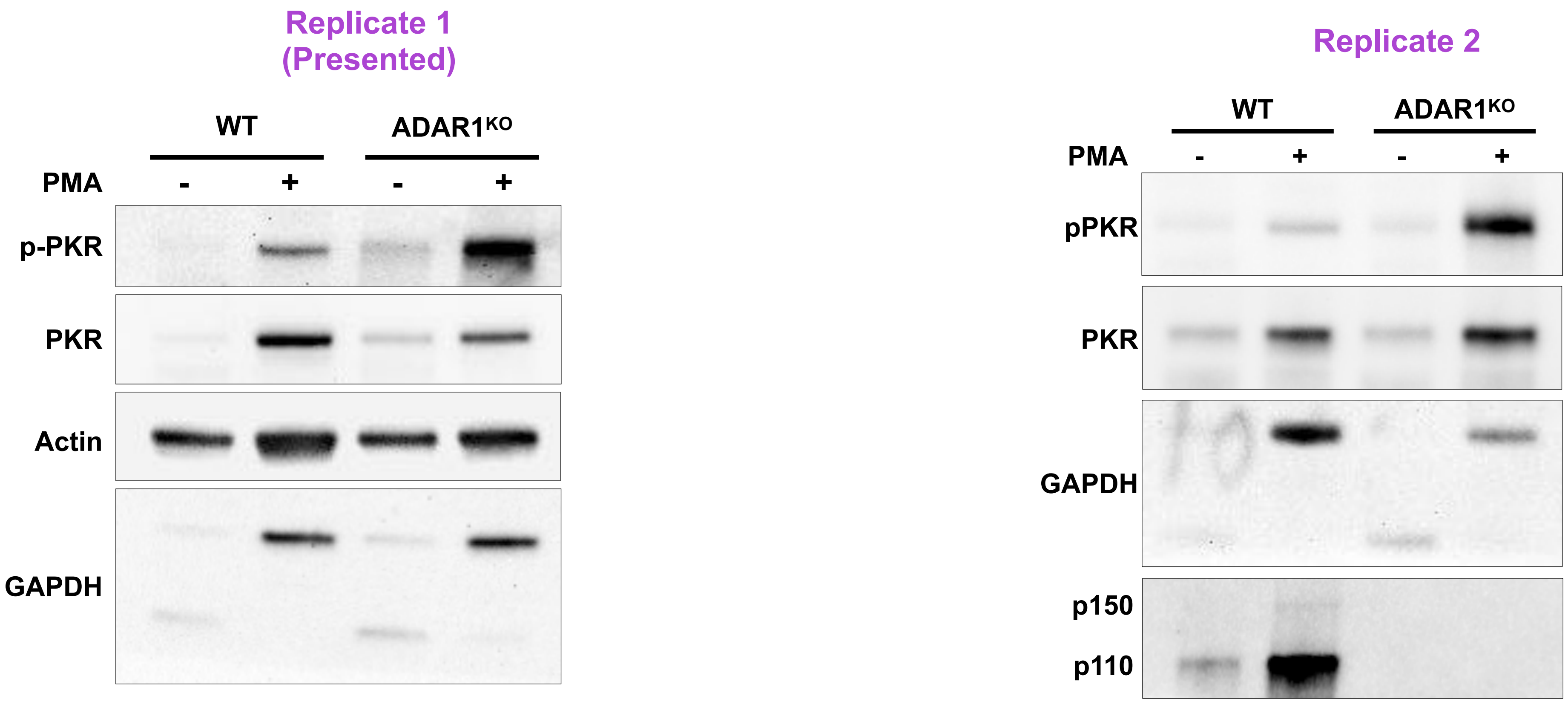
Replicate 1
(Presented)
WT
ADAR1KO
PMA
-
+
-
+
p-PKR
PKR
Actin
GAPDH
Replicate 2
WT
ADAR1KO
PMA
-
+
-
+
pPKR
PKR
GAPDH
p150
p110

### Figure 4E

Replicate 1
(Presented)

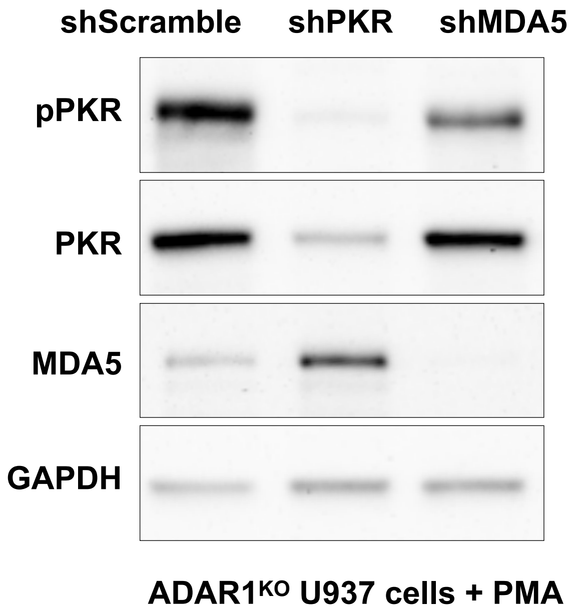


Replicate 2

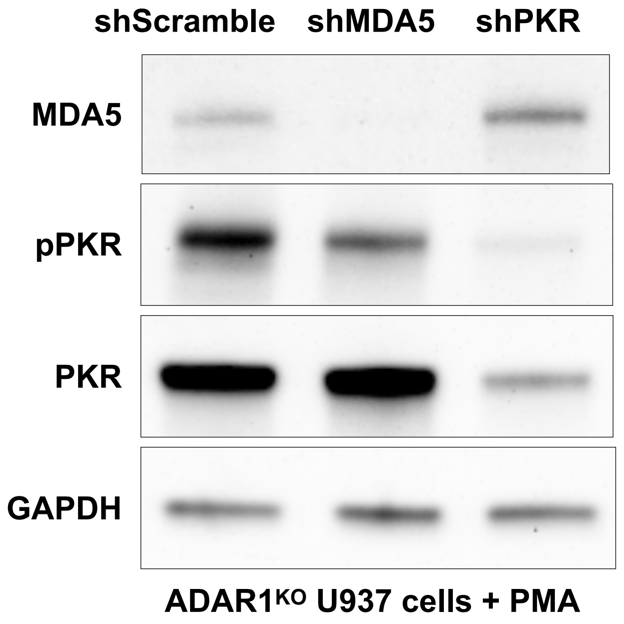

#### Figure 5A

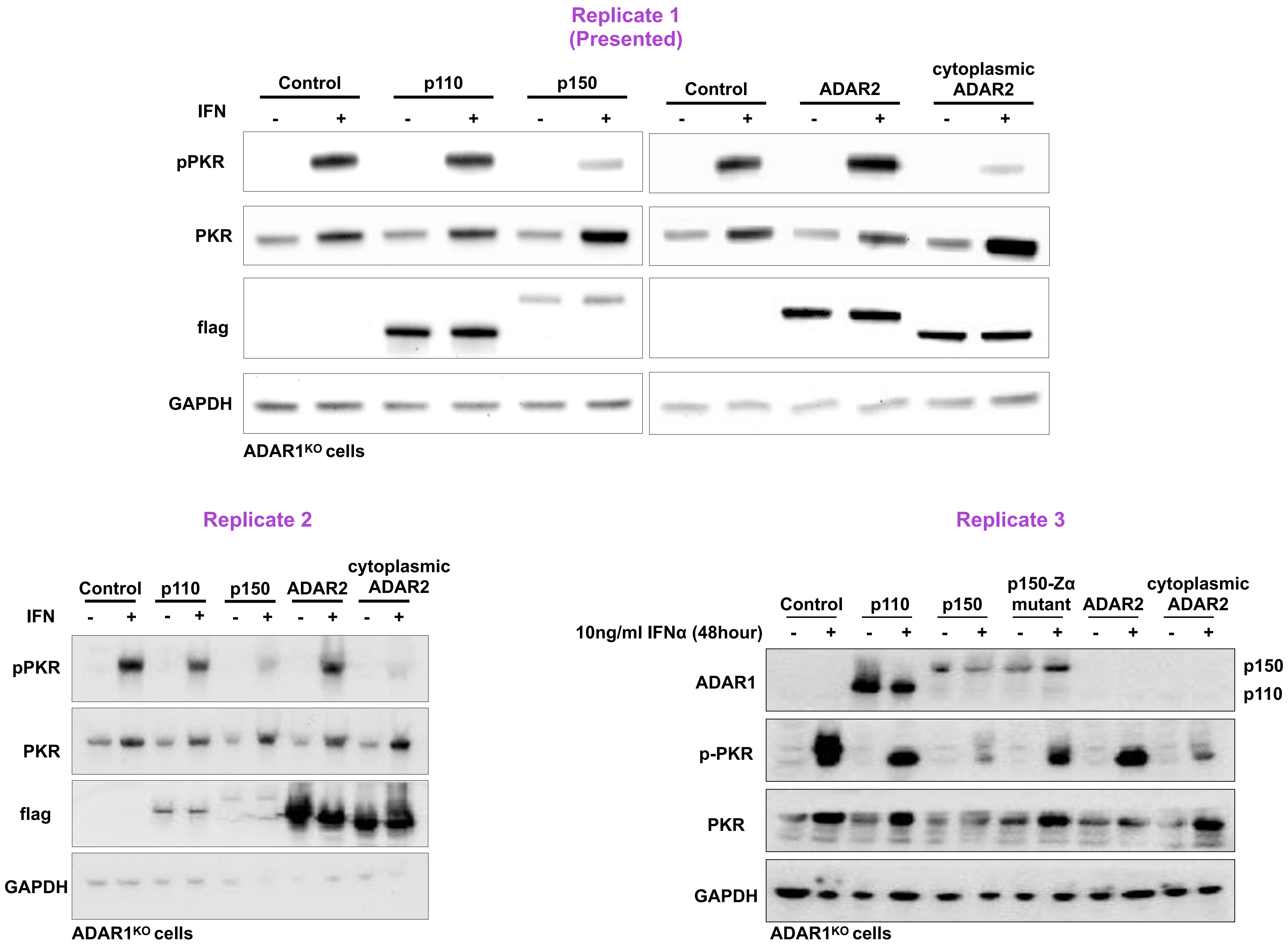

Replicate 1
(Presented)
Control
p110
p150
Control
ADAR2
cytoplasmic ADAR2
IFN
- + - + - + - + - + - +
pPKR
PKR
flag
GAPDH
ADAR1^KO cells
Replicate 2
Control
p110
p150
ADAR2
cytoplasmic ADAR2
IFN
- + - + - + - + - +
pPKR
PKR
flag
GAPDH
ADAR1^KO cells
Replicate 3
Control
p110
p150
p150-Zα mutant
ADAR2
cytoplasmic ADAR2
10ng/ml IFNα (48hour)
- + - + - + - + - + - +
ADAR1
p150
p110
p-PKR
PKR
GAPDH
ADAR1^KO cells

### Figure 5C

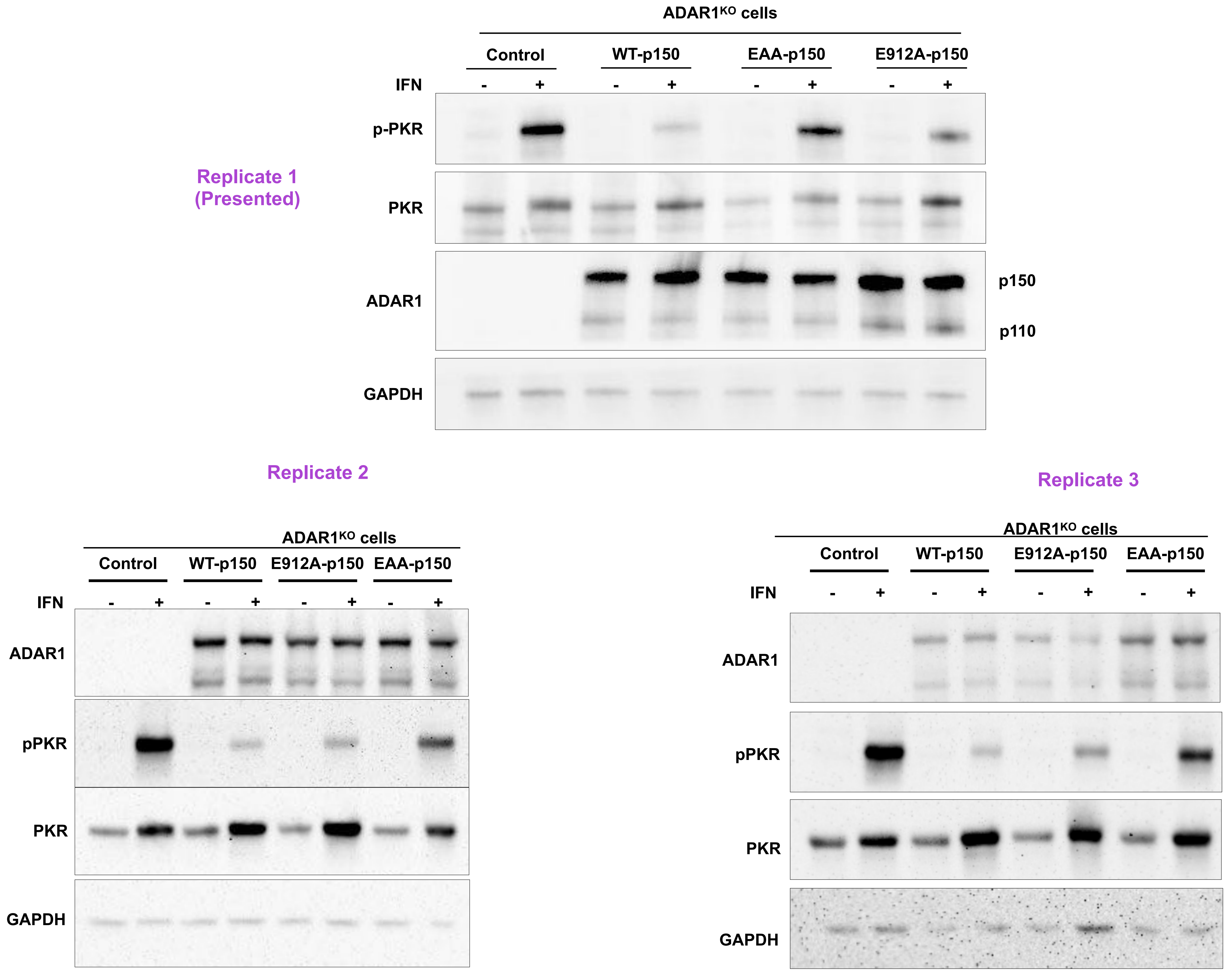
ADAR1KO cells
Control
WT-p150
EAA-p150
E912A-p150
IFN
- + - + - + - +
p-PKR
Replicate 1
(Presented)
PKR
ADAR1
p150
p110
GAPDH
Replicate 2
ADAR1KO cells
Control
WT-p150
E912A-p150
EAA-p150
IFN
- + - + - + - +
ADAR1
pPKR
PKR
GAPDH
Replicate 3
ADAR1KO cells
Control
WT-p150
E912A-p150
EAA-p150
IFN
- + - + - + - +
ADAR1
pPKR
PKR
GAPDH

### Figure 6A

Replicate 1
(Presented)

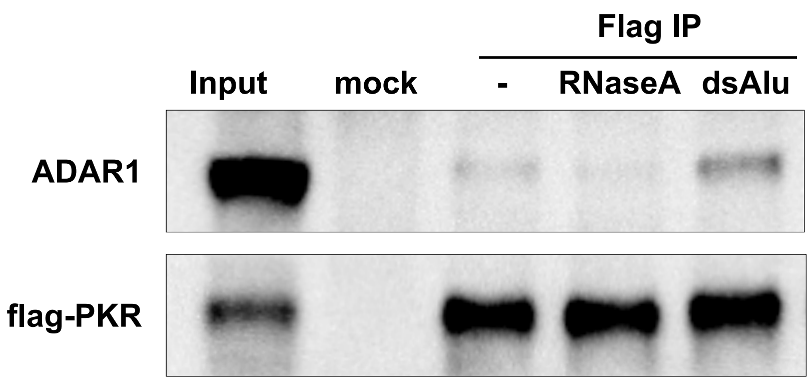

### Figure 6C

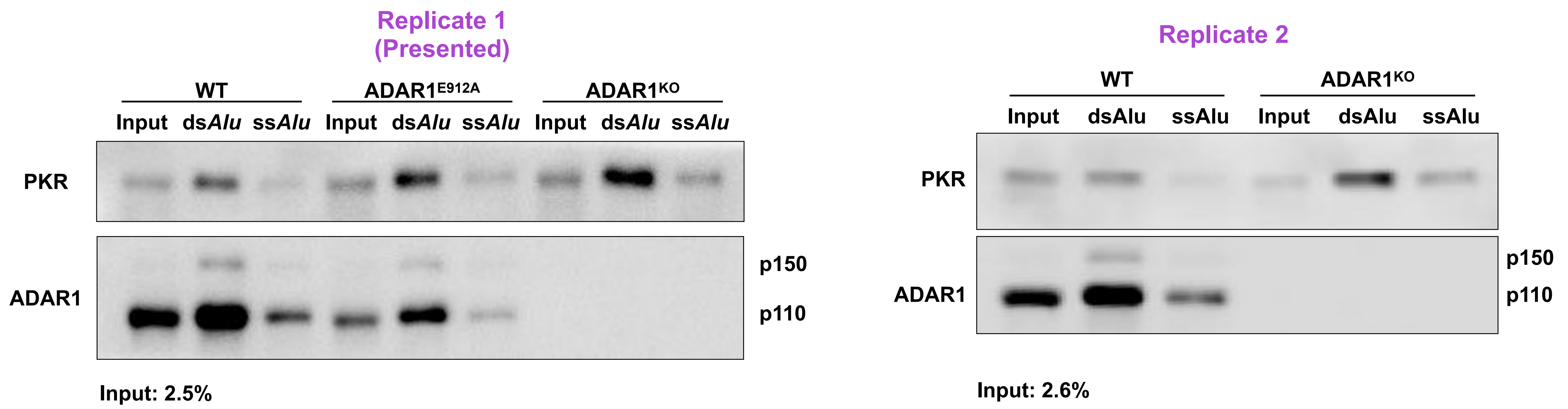
Replicate 1
(Presented)
WT
ADAR1E912A
ADAR1KO
Input dsAlu ssAlu Input dsAlu ssAlu Input dsAlu ssAlu
PKR
ADAR1
p150
p110
Input: 2.5%
Replicate 2
WT
ADAR1KO
Input dsAlu ssAlu Input dsAlu ssAlu
PKR
ADAR1
p150
p110
Input: 2.6%

### Figure 6D

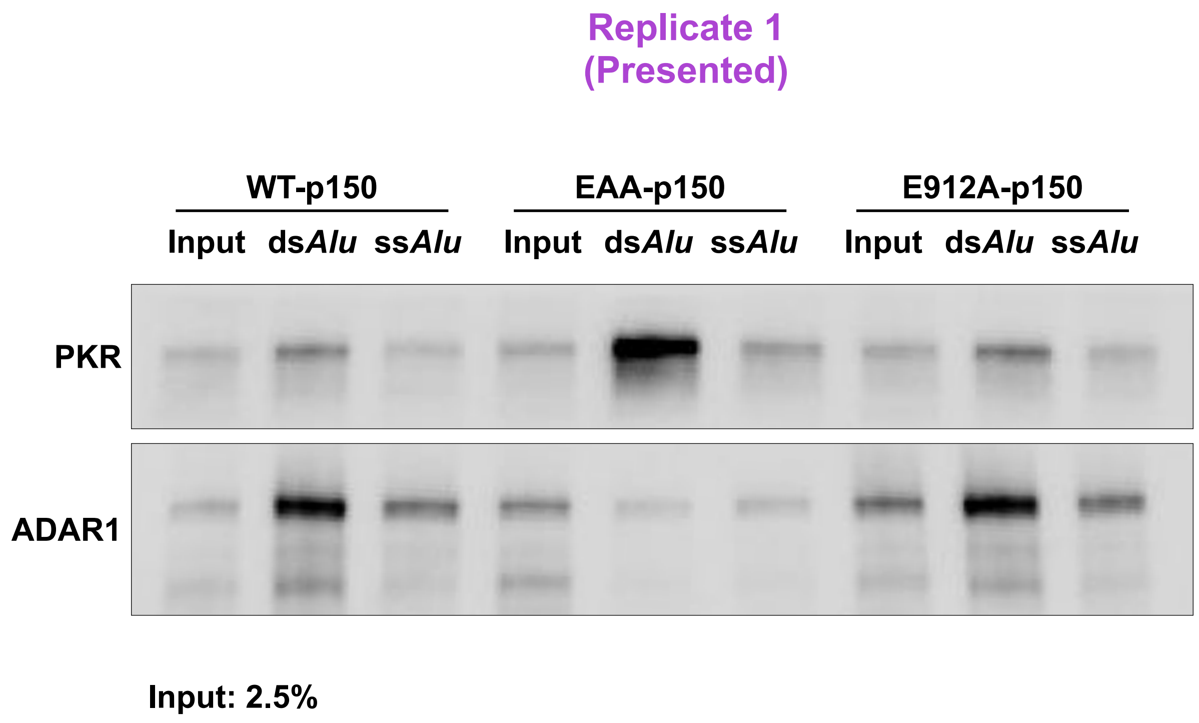
Replicate 1
(Presented)
WT-p150
EAA-p150
E912A-p150
Input dsAlu ssAlu
Input dsAlu ssAlu
Input dsAlu ssAlu
PKR
ADAR1
Input: 2.5%

### Figure 6E

Replicate 1
(Presented)

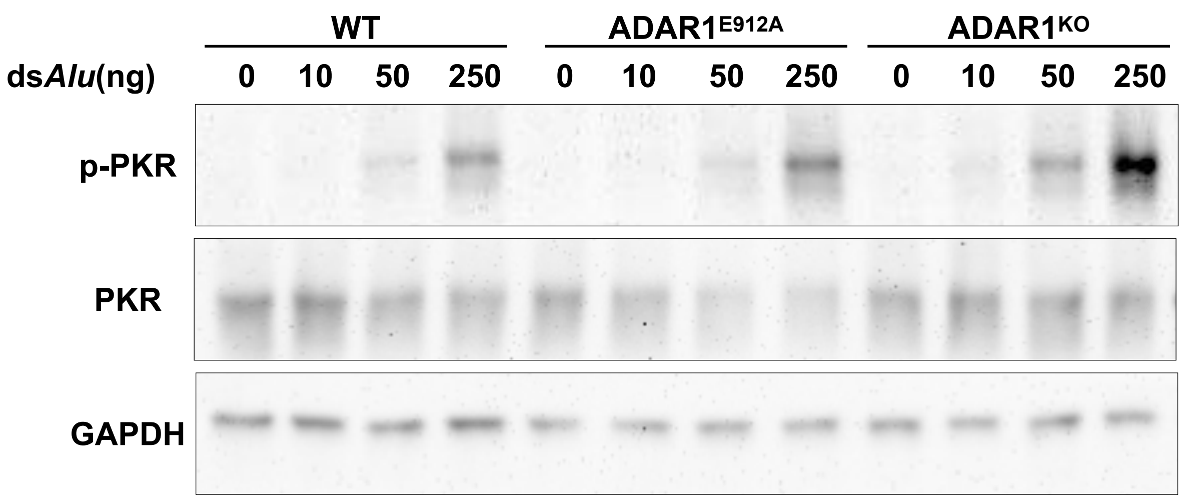


Replicate 2
(Presented in Figure S6B)

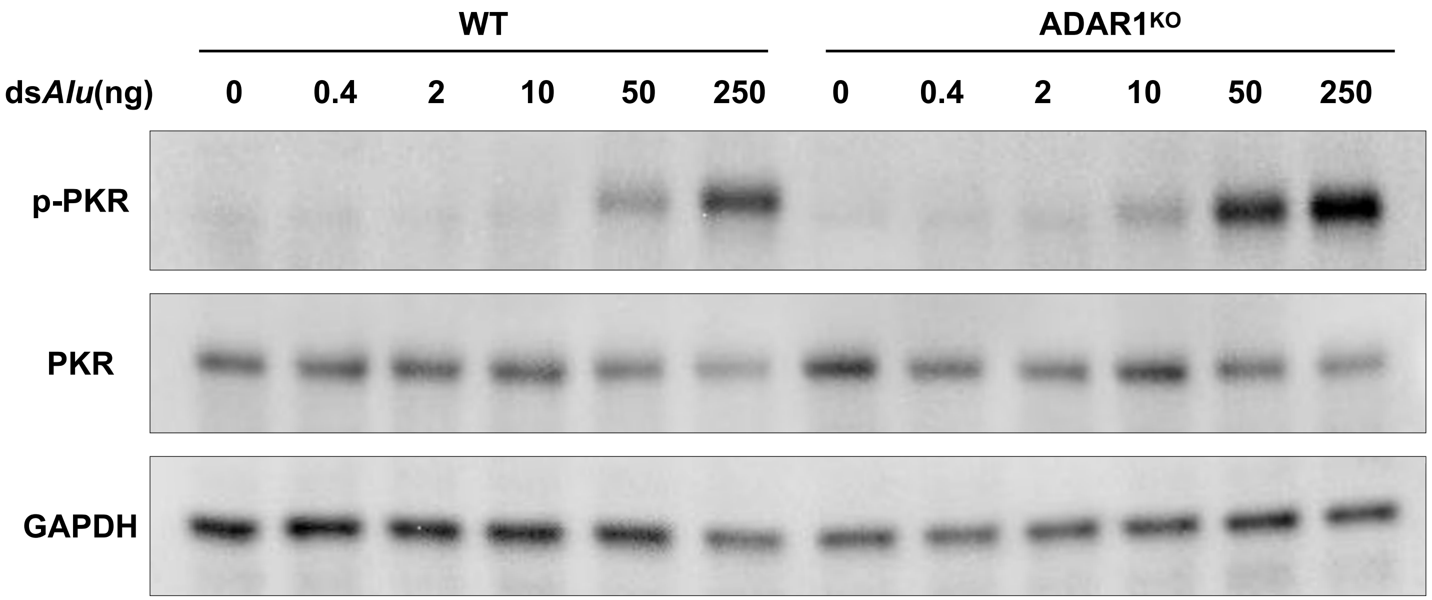

### Figure S6A

Replicate 1
(Presented)

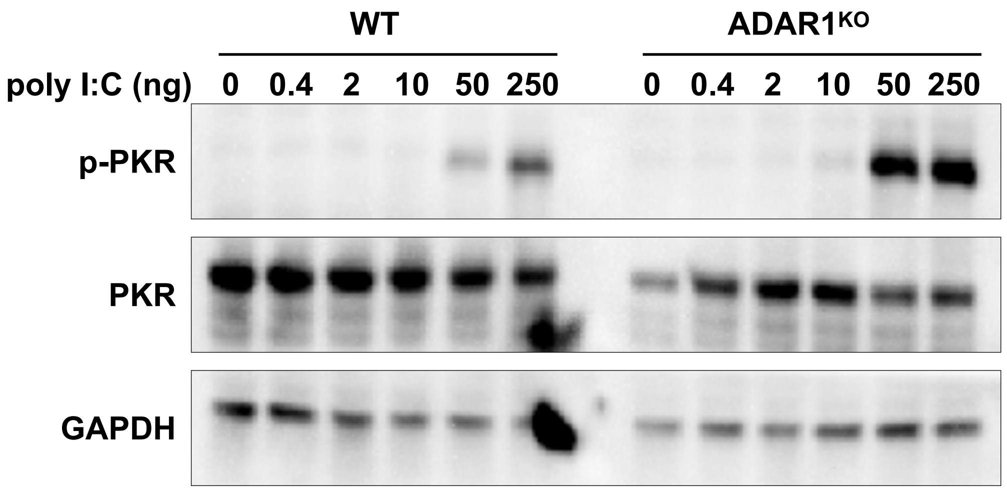


12 well plate. harvest 8h after transfection

Replicate 2

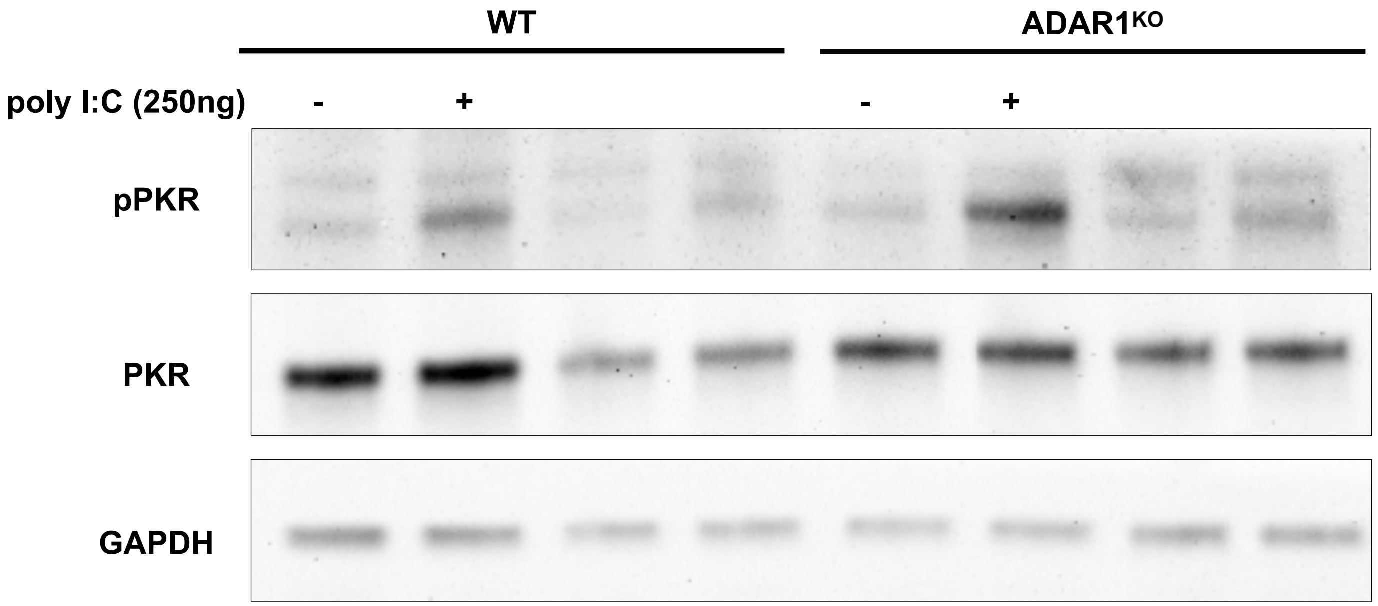


6 well plate. 250 ng/ml poly I:C for 6h.

### Figure 6G

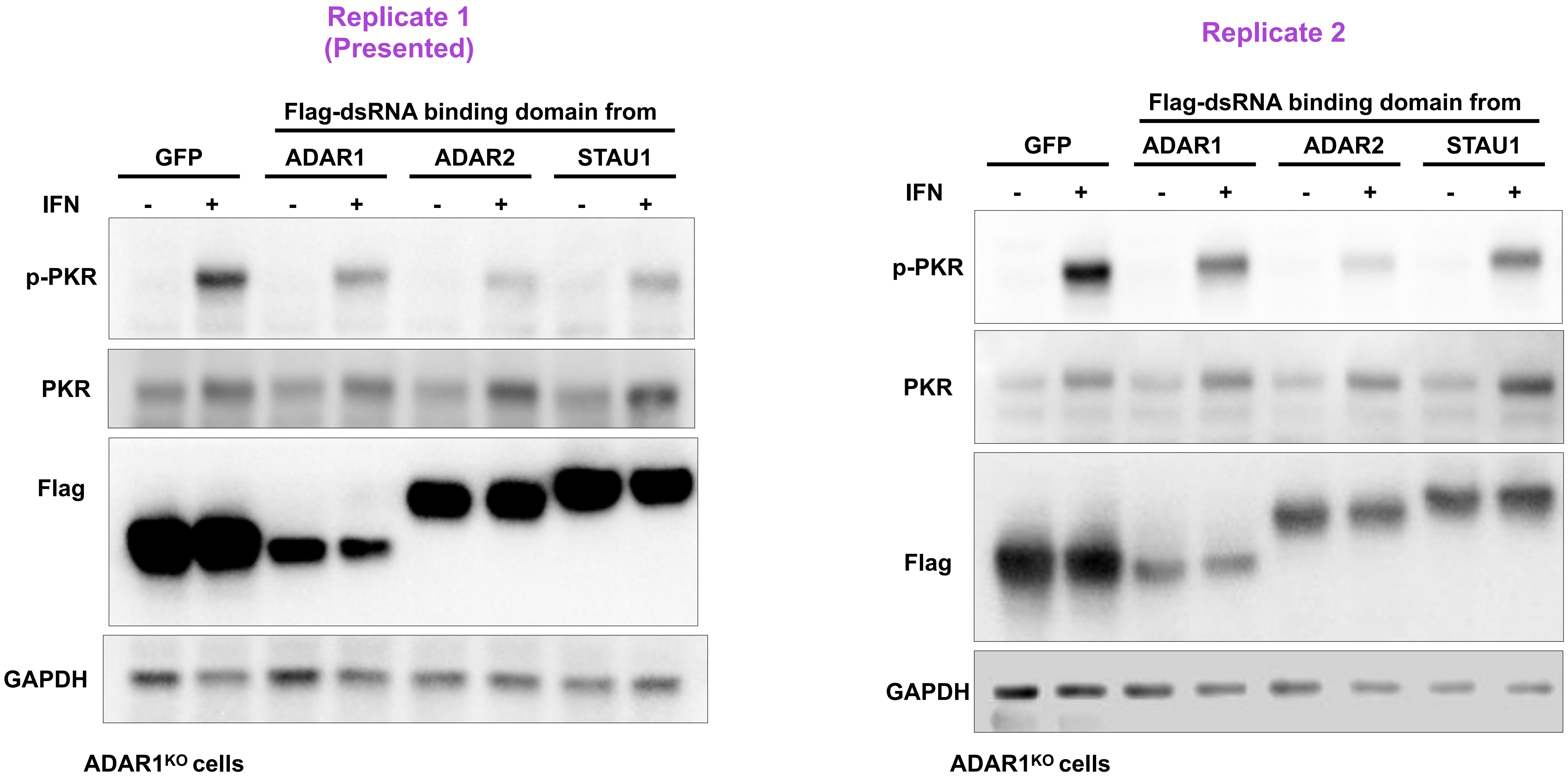
Replicate 1
(Presented)
Flag-dsRNA binding domain from
GFP
ADAR1
ADAR2
STAU1
IFN
- + - + - + - +
p-PKR
PKR
Flag
GAPDH
ADAR1KO cells
Replicate 2
Flag-dsRNA binding domain from
GFP
ADAR1
ADAR2
STAU1
IFN
- + - + - + - +
p-PKR
PKR
Flag
GAPDH
ADAR1KO cells
