## Supplemental Data S2 for "ADAR1p150 Prevents MDA5 and PKR Activation via Distinct Mechanisms to Avert Fatal Autoinflammation"

**Contacts:**

(text removed CW190123)

Phenomics Australia Histopathology and
Slide Scanning Service
The University of Melbourne
Department of Anatomy and Neuroscience
Grattan Street, PARKVILLE, VIC 3010

(text removed CW190123)

### 9.1 Histopathology Report

**Case Number** APN22/057 St Vincent's Institute (Carl Walkley)

**Registration Date** Tue 30/08/2022

**A1 genotype**

**Animal Details**

**WT** 154, Adar1L196C/L196C Ifih1-/- Pkr-/-
DOB: 171/22 (20/06/22), 107 days old, Female, 18.7g, Black

**WT** 491, Adar1-/- Ifih1-/- PKR-/-
DOB: 154/22 (03/06/22), 124 days old, Female, 23.3g, Black

**WT** 499, Adar1-/- Ifih1-/- PKR-/-
DOB: 161/22 (10/06/22), 117 days old, Male, 27.0g, Black

**p150-/-** 151, Adar1L196C/L196C Ifih1-/- Pkr-/- (triple mutant)
DOB: 169/22 (18/06/22), 109 days old, Female, 21.4g, Black

**p150-/-** 155, Adar1L196C/L196C Ifih1-/- Pkr-/- (triple mutant)
DOB: 171/22 (20/06/22), 107 days old, Female, 20.0g, Black

**p150-/-** 167, Adar1L196C/L196C Ifih1-/- Pkr-/-
DOB: 197/22 (16/07/22), 81 days old, Male, 26.9g, Black

**Adar1-/-** 492, Adar1-/- Ifih1-/- PKR-/- (triple mutant)
DOB: 154/22 (03/06/22), 124 days old, Female, 18.7g, Black

**Adar1-/-** 500, Adar1-/- Ifih1-/- PKR-/- (triple mutant)
DOB: 161/22 (10/06/22), 117 days old, Male, 20.4g, Black

**Adar1-/-** 513, Adar1-/- Ifih1-/- PKR-/-
DOB: 164/22 (10/06/22), 117 days old, Male, 23.1g, Black

**DoD / Necropsy** Wed 05/10/2022

**Death** $CO_2$

**Origin** St Vincent's Institute

**Treatment** Knock-out
Animals will be 81-124 days on 5th Oct; mixed sexes.

Three control, 3 Adar1 deficient, 3 adar1p150 isoform deficient.
All MDA5 PKR null (Ifih1-/- Eif2ak2-/-).

? Grossly normal weights; (text removed CW190123)

(C.Walkley 30/08/2022)

**Species / Breed / Strain** C57BL/6

**Animal Health Facility** St. Vincent's Bioresources Centre

The mice were housed in Room 5.

POSITIVE for Helicobacter spp, Helicobacter hepaticus, Mouse Norovirus, Chilomastix bettencourti, Entamoeba muris.

**Organs Examined**

Adrenal glands, Bladder, Bone marrow, Brain, Cecum, Cervix, Clitoral gland, Colon, Duodenum, Epididymes, Eyes, Gall bladder, Harderian glands, Head, Heart, Hind leg (Long bone, Bone marrow, Synovial joint, Skeletal muscle), Ileum, Jejunum, Kidney, Liver, Lungs, Mammary tissue, Mesenteric lymph node, Ovaries, Oviducts, Pancreas, Penis, Preputial gland, Prostate glands, Salivary glands and Regional lymph nodes, Seminal vesicles, Skin, Spinal cord, Spleen, Sternum, Stomach, Tail, Testes, Thymus, Thyroids, Trachea, Uterus, Vagina

**Macroscopic Observations**

P.A. iLab request number: PAHSSS-CW-196
Date of Necropsy: October 05, 2022 09:00
Courier/Transportation details: text removed CW190123

Body Condition Scoring (BCS): Scale of 1-5, animals scored a BCS of 3 or 4.
5: The mouse is obese, and bones cannot be felt at all
4: The mouse is well-fleshed, and bones are barely felt
3: The mouse is in optimal condition. The bones are palpable but not prominent
2: The mouse is thin, and bones are prominent
1: Muscle wasting is advanced; fat deposits are gone, and bones are very prominent.

At the time of necropsy, animals appeared well nourished, well groomed, active/curious and healthy with normal movement and gait. There were no observable dermal lesions and no nasal/ocular discharges. The gastrointestinal tract contained ample ingesta and the thoracic and most abdominal viscera showed no macroscopic abnormalities.
Please refer to individual animals for more macroscopic details.

Body weights (g):
Adar1L196C/L196C Ifih1-/- Pkr-/- animals.
Female mice showed higher average total body weight compared to their control. No concurrent male control for comparison.

Adar1-/- Ifih1-/- PKR-/- animals.
Both male and female mice showed lower average total body weights compared to their controls.

Blood Report:
Adar1L196C/L196C Ifih1-/- Pkr-/- animals.
Blood results show mostly comparable readings between the Adar1 mutant animals and their control.
Some notable findings:
151 Female – Low platelets (PLT) and elevated mean platelet volume (MPV).
151 Female – Low lymphocytes (%LYM and AbsLymph).
154 Female (control) – Elevated white blood cells (WBCB and WBCP).
154 Female (control) – Elevated lymphocytes (AbsLymph).
155 Female – Elevated red blood cell distribution width (RDW).
167 Male – Elevated red blood cell distribution width (RDW).
167 Male – Low neutrophils (%NEUT and AbsNeut).

Adar1-/- Ifih1-/- PKR-/- animals.
Blood results show mostly comparable readings between the Adar1 mutant animals and their controls.
Some notable findings:
492 Female – Elevated red blood cell distribution width (RDW).
492 Female – Low neutrophils (%NEUT and AbsNeut).

500 Male – Elevated red blood cell distribution width (RDW).
513 Male – Elevated red blood cell distribution width (RDW).
513 Male – Low neutrophils (%NEUT and AbsNeut).

Note: haematology values can vary with mouse strain/stock, age, sex, blood sampling method, fasting and environmental conditions, pathogen status, and laboratory. The reference intervals used for this report are based on published values of adult mice at 16 weeks of age.
For more details please see the accompanying APN22/057SVI (C. Walkley) Blood Report.

#### Microscopic Observations

SUMMARY of micromorphological changes.
Across multiple tissues, all animals showed various instances of known incidental, background, or spontaneous changes seen in mice.

Relative to the controls, mutant animals in this cohort showed changes in:
(1) Peripheral blood smear – anitocytosis (microcytes) and poikilocytosis.
(2) Small intestine – increased apoptotic bodies.
(3) Spleen – increased extramedullary haemoatopoiesis, and possible decreased ratio of white pulp to red pulp.

No abnormalities detected in the thymus.

Notes:
1. Reactive lymph nodes are defined as mild follicular hyperplasia, germinal centre formation and occasional sinus histiocytosis - a common finding in mice.
2. Hyperplastic lymph node follicles are identified by an increase in number and size of follicles and conversion to secondary follicles. Hyperplasia of the paracortex is characterized by an increase in the cell density and, depending on the degree of hyperplasia, an increase in the paracortical area.
3. Accumulation of leukocytes and other cells are common nonneoplastic lesions in many tissues. The term "inflammation" is used when the cell (leukocyte) accumulations are part of an active inflammatory process (typified by concurrent features such as vascular changes, necrosis, fibrosis, and/or tissue disruption). In contrast, cell "infiltration" is used when the cell (e.g., lymphocyte) accumulations are present in tissue without other disruption or pathology.
4. Focal inflammatory cell aggregates consisting of mononuclear, polymorphonuclear, and/or histiocytic cells are frequently observed in ageing mice (Maranport RR. 1999. Pathology of the Mouse.). These can be present as lymphoid aggregates found in various tissues including the renal pelvis, bladder, lungs, liver (Pettan-Brewer C and Treuting PM. 2011. Practical pathology of aging mice.) and salivary glands (Haines DC, Chattopadhyay S and Ward JM. 2001. Pathology of Aging B6;129 Mice).
5. Mild extramedullary haematopoiesis (EMH) in the red pulp of the spleen is a common finding in the mouse. EMH consists of erythroid precursors, myeloid precursors, megakaryocytes or all three. While some degree of extramedullary haematopoiesis is present in normal rodents, especially in mice, increased extramedullary haematopoiesis can result from haematotoxin insult, systemic anaemia, and infections elsewhere in the body. (Suttie AW. 2006. Histopathology of the spleen).
6. Cytoplasmic inclusions of homogeneous eosinophilic hyaline-like material may be seen in older mice in intrahepatic biliary epithelial cells, as well as epithelial cells in the gallbladder. In marked cases, there is hyperplasia of the glandular epithelium, and crystalline forms of the eosinophilic inclusion material may be present both intracellularly and extracellularly (Maronpot RR. 1999. Liver and gallbladder. In: Pathology of the Mouse: Reference and Atlas).
7. Focal fatty change of the liver can be a spontaneous lesion and may be more common in some strains than others (Maronpot RR. 2014. Liver-Fatty Change. In: National Toxicology Program Nonneoplastic Lesion Atlas).
8. Fatty change of the liver may occur in mice as a response to a toxicant. It is also seen in old obese controls and is more common in male than in female mice. The degree of fatty metamorphosis may vary and usually starts with a centrilobular distribution (Frith CH, Ward JM. 1988. Digestive System. In: Color Atlas of Neolastic and Non-neoplastic Lesions in Aging Mice).
9. Focal pancreatic infiltrates of lymphocytes, plasma cells, and macrophages are uncommon in aged mice. The infiltrates are generally minimal and may be associated with atrophy (Maronpot RR. 1999.

Exocrine and Endocrine Pancreas. In: Pathology of the Mouse: Reference and Atlas).
10. Valvular myxomatous changes or degeneration can be an age-related spontaneous or chemical-induced change. The lesion is characterized by focal or segmental thickening of the subendocardium in the valve leaflets and expansion of the spongiosa of the valve leaflet with extracellular fibromyxoid material composed predominantly of glycosaminoglycans. Occasionally, fibrin deposits or thrombi and collections of neutrophils or mononuclear cells are seen (Johnson CL, Nyska A. 2017. Heart, Valve, – Degeneration. In: National Toxicology Program Nonneoplastic Lesion Atlas).
11. A number of intestinal parasites may be seen within the lumen of the large intestine, the cecum, and less commonly the small intestine. The significance and presence of protozoan organisms is questionable since infected animals are normally asymptomatic. The presence of Protozoa such as flagellates, ciliatesm etc. have not been associated with any microscopic lesions of clinical significance in mice (Maronpot RR. 1999. Intestines and Mesentry: Small and Large Intestine. In: Pathology of the Mouse: Reference and Atlas).
12. Focal hepatic necrosis is a non-specific entity quite often encountered as an incidental finding in the liver of mice. It can be the result of viruses (mouse hepatitis), bacteria (Clostridium piliforme), toxicants, and ischemia while the etiology is often unknown. It may involve single cells, single or multiple lobules, and it may vary in distribution. Coagulation necrosis with distinct eosinophilic cytoplasm and pyknotic or absent nuclei is the typical morphologic feature (Frith CH, Ward JM. 1988. Color Atlas of Neolastic and Non-neoplastic Lesions in Aging Mice).
13. Ovarian cysts are a common finding in rats and mice. Ovarian cysts may be unilateral or bilateral, single or multiple, and may become quite large… size and number of cysts increase with age. (Willson G, Cimon KY. 2015. Ovary - Cyst. In: National Toxicology Program Nonneoplastic Lesion Atlas.)
14. Hydrocephalus may be communicating or noncommunicating; that is, the former has no apparent obstructive process, whereas the latter has an obstructive cause somewhere in the ventricular connections. Most commonly, communicating hydrocephalus is considered to result from an idiopathic increase in cerebrospinal fluid production or deceased resorption. (Little P, Rao DB. 2014. Brain – Hydrocephalus. In: National Toxicology Program Nonneoplastic Lesion Atlas).
15. Germ cell degeneration of the testes is a nonspecific term that generally includes a number of degenerative features, such as tubular vacuolation, partial depletion of germ cells, degenerating (multinucleated or apoptotic) germ cells, and disordered arrangement of the germ cell layers. Chemically induced germ cell degeneration can be multifocal in distribution, but it is most often a bilateral lesion that affects most of the seminiferous tubules to varying degrees. It can also be an incidental background finding in rats and mice of any age, but the incidence increases with age. (Wilson G and Cimon K Y. 2015. Testis, Germ cell – Degeneration. In: National Toxicology Program Nonneoplastic Lesion Atlas).
16. Cardiomyocyte vacuolation is considered to be a degenerative process, consisting of multifocal or widespread accumulation of multiple, well demarcated, round, variably sized (primarily small), clear vacuoles. Vacuoles are found within the cardiomyocyte sarcoplasm and occasionally coalesce into larger vacuoles. Myofiber cytoplasmic vacuolation can be the only morphologic manifestation seen in cardiotoxicity or may be associated with other changes reflecting cardiotoxicity (e.g., myofiber necrosis, mononuclear cell infiltration, fibrosis), or it may be a minor component in the case of spontaneous cardiomyopathy. (Johnson CL and Nyska A. 2017. Heart, Myocardium – Vacuolation, Cytoplasmic. In: National Toxicology Program Nonneoplastic Lesion Atlas).

#### 154 (control)

Adar1+/+ Ifih1-/- Eif2ak2-/-; DOB20/6/2022; female

Macro Observations

Tail suspension test for neurological defects - negative.
Dentition, tongue and oral cavity was unremarkable.
BCS: 3
Spleen: 14x4x2mm
Kidneys: 10x6x4mm, symmetrical
Thymus: 7x7x2mm
Lungs inflated.
Heart: 10x7x6mm
Brain: 14x10x5mm, symmetrical
Pituitary gland identified, macroscopically normal.
Tail length: 73mm (straight)
Head harvested for evaluation of auditory and vestibular structures.
Bone marrow smear taken from left hind leg.

No macroscopic lesions identified.

Micro Observations

Animal 154 was used as a female histological control for the Adar1L196C/L196C Ifih1-/- Pkr-/- animals in this cohort.

Marrow smear: Examination of the smear showed representative cells from the myeloid and erythroid series. Occasional cells from the lymphoid series. Occasional and unremarkable megakaryoblasts.
(91381)

Peripheral blood smear: Examination of the smear showed red blood cells (majority of cells shown), occasional white blood cells including lymphocytes, segmented neutrophils, monocytes and platelets (clumps). No discernible morphological changes or detectable parasites.
(91382)

Organs examined: Mammary glands (91388, 91391), Ovaries/oviducts (91383), Uterus/cervix/vagina (91383), Urinary Bladder (91383), Liver/Gall Bladder (91384), Stomach (91385), Duodenum/jejunum/ileum/GALT (91385, 91386, 91387), Cecum/Colon/GALT (91387), Mesenteric Lymph Node (91388), Spleen (91383), Pancreas (91383), Kidneys/Adrenal Gland (91389), Salivary Glands/regional lymph nodes (91388), Thyroids (91390), Trachea/Lungs (91390), Thymus (91390), Heart (91390), Skin (91391), Tail (91392), Eyes/Harderian Glands (91393), Brain (91394), Spinal cord (91395, 91396), Hind leg (91397, 91401), Head (91398, 91399), Sternum (91400).

Micromorphological changes-
Liver: minimal perivascular mononuclear cell infiltrates (91384).
Liver: several small parenchymal leukocyte aggregates with some hepatocyte cell loss/necrosis (91384).
Pancreas: minimal perivascular mononuclear cell infiltrates (91385).
Small intestine: increased goblet cells and intracellular granules in the Paneth cells particularly in the ileum and distal jejunum (91386, 91387). -Pathology to comment-
Small intestine: ileum also showed increased leukocyte infiltrates in the lamina propria (91387). -Pathology to comment-
Large intestine: abundant luminal protozoa (91837).
Large intestine: colon shows increased leukocyte cell infiltrates in the lamina propria, mild (91387). -Pathology to comment-
Salivary glands: minimal perivascular mononuclear cell infiltrates within the submandibular gland (91388).
Kidneys: minimal peripelvic mononuclear cell infiltrates (91389).
Heart: myxomatous valvular changes (thickened leaflet) (91390).
Lungs: minimal perivascular mononuclear cell infiltrates (91390).
Skin: mild focal polymorphonuclear cell inflammation in the subjacent abdominal wall muscle (91391).
Spinal cord (skin): subcutaneous oedematous changes (91395, 97396).
Hind leg: small focal leukocyte aggregate in the skeletal muscle (91397, 91401).
Hind leg: cortical bone changes in the tibia (91397), query plane of section artefact. -Pathology to comment-
Hind leg: subcutaneous oedematous changes in the distal hind leg and foot (91397, 91401).
Hind leg: mild focal polymorphonuclear cell infiltrates in the foot pad (91397, 91401).
Hind leg: query focal cortical bone changes (femur) near the knee joint (91401). -Pathology to comment-

#### 491 (control)

Adar1+/+ Ifih1-/- Eif2ak2-/-; DOB: 03/06/2022; female

Macro Observations

Tail suspension test for neurological defects - negative.
Dentition, tongue and oral cavity was unremarkable.
BCS: 3
Spleen: 16x4x2mm
Kidneys: 12x8x5mm, symmetrical
Thymus: 7x7x2mm

Lungs not inflated.
Heart: 12x8x6mm
Brain: 16x10x5mm, symmetrical
Pituitary gland identified, macroscopically normal.
Tail length: 82mm (straight)
Head harvested for evaluation of auditory and vestibular structures.
Bone marrow smear taken from left hind leg.

Small intestine appeared twisted and nodular.
No other macroscopic lesions identified.

Micro Observations

Animal 491 was used as a female histological control for Adar1-/- Ifih1-/- PKR-/- animals in this cohort.

Marrow smear: Examination of the smear showed representative cells from the myeloid and erythroid series. Occasional cells from the lymphoid series. Occasional and unremarkable megakaryoblasts.
(91447)

Peripheral blood smear: Examination of the smear showed red blood cells (majority of cells shown), occasional white blood cells including lymphocytes, segmented neutrophils, monocytes and platelets (clumps). No discernible morphological changes or detectable parasites. Sub-optimal staining.
(91448)

Organs examined: Mammary glands (91455, 91458), Ovaries/oviducts (91449, 91450), Uterus/cervix/vagina (91449, 91450), Urinary Bladder (91449, 91450), Liver/Gall Bladder (91451), Stomach (91452), Duodenum/jejunum/ileum/GALT (91452, 91453, 91454), Cecum/Colon/GALT (91455), Mesenteric Lymph Node (91449, 91450), Spleen (91449, 91450), Pancreas (91449, 91450), Kidneys/Adrenal Gland (91456), Salivary Glands/regional lymph nodes (91455), Thyroids (91457), Trachea/Lungs (91457), Thymus (91457), Heart (91457), Skin (91458), Tail (91459), Eyes/Harderian Glands (91460), Brain (91461), Spinal cord (91462, 91463), Hind leg (91464), Head (91465, 91466), Sternum (91467).

Micromorphological changes-
Ovary: small sized ovarian cyst (91450).
Liver: multiple small parenchymal leukocyte aggregates with some hepatocyte cell loss/necrosis (91451).
Liver: minimal perivascular mixed leukocyte infiltrates (91451).
Gall bladder: segment with epithelial hyperplasia and polymorphonuclear cell infiltration in the underlying lamina propria (91451).
Stomach: of the glandular portion, minimal focal mixed leukocyte infiltrates in the lamina propria (91452).
Small intestine: mainly within the ileum and distal jejunum - increased goblet cells, increased intracellular granules in the Paneth cells, and increased leukocyte infiltrates in the lamina propria (91453, 91454). -Pathology to comment-
Small intestine: two large submucosal leukocyte aggregates, query Peyer's Patches (91454). -Pathology to comment-
Large intestine (cecum): scattered luminal protozoa (91454).
Large intestine (colon): mild increase of leukocyte cell infiltrates in the lamina propria (91454).
Kidneys: multiple protein casts in the medulla, few are mildly dilated (91456).
Kidneys: minimal perivascular/peripelvic mononuclear cell infiltrates (91456).
Lungs: various degrees of parenchymal congestion and collapse (atelectasis), judged to be artefactual (91457).
Lungs: minimal perivascular mononuclear cell infiltrates (91457).
Heart: myxomatous valvular changes (thickened leaflets) (91457).
Hind leg: cortical bone changes (femur) near the knee joint (91464). -Pathology to comment-

#### 499 (control) Adar1+/+ Ifih1-/- Eif2ak2-/-, DOB 10/6/2022; male

Macro Observations

Tail suspension test for neurological defects - negative.
Dentition, tongue and oral cavity was unremarkable.
BCS: 4
Testes: 7x5x3mm, symmetrical
Spleen: 15x4x2mm
Kidneys: 12x6x5mm, symmetrical
Thymus: 7x7x2mm
Lungs inflated.
Heart: 10x8x6mm
Brain: 15x10x5mm, symmetrical
Pituitary gland identified, macroscopically normal.
Tail length: 81mm (straight)
Head harvested for evaluation of auditory and vestibular structures.
Bone marrow smear taken from left hind leg.

No macroscopic lesions identified.

Micro Observations

Animal 499 was used as a male histological control for the Adar1-/- Ifih1-/- PKR-/- animals in this cohort.

Marrow smear: Examination of the smear showed representative cells from the myeloid and erythroid series. Occasional cells from the lymphoid series. Occasional and unremarkable megakaryoblasts.
(91490)

Peripheral blood smear: Examination of the smear showed red blood cells (majority of cells shown), occasional white blood cells including lymphocytes, segmented neutrophils, monocytes and platelets (clumps). No discernible morphological changes or detectable parasites.
(91491)

Organs examined: Testes/Epididymes (91492), Seminal vesicles (91493), Prostate glands (91493), Penis/Preputial gland (91494), Urinary Bladder (91493), Liver/Gall Bladder (91495), Stomach (91496), Duodenum/jejunum/ileum/GALT (91496, 91497, 91498), Cecum/Colon/GALT (91498), Mesenteric Lymph Node (91499), Spleen (91493), Pancreas (91493), Kidneys/Adrenal Glands (91500), Salivary Glands/regional lymph nodes (91499), Thyroids (91501), Trachea/Lungs (91501), Thymus (91501), Heart (91501), Skin (91502), Tail (91503), Eyes/Harderian Glands (91504), Brain (91505), Spinal cord (91506, 91507), Hind leg (91508, 91509), Head (91558, 91559), Sternum (91510).

Micromorphological changes-
Liver: multiple small parenchymal leukocyte aggregates and some hepatocyte cell loss/necrosis (91495).
Liver: small cluster of vacuolated hepatocytes resembling hydropic degeneration or fatty change (steatosis) (91495).
Gall bladder: epithelial hyperplasia, cytoplasmic hyaline droplet accumulation, and polymorphonuclear cell infiltrates (91495).
Stomach: of the glandular portion, marked and mainly submucosal polymorphonuclear cell infiltrates (91496).
Small intestine: mainly within the ileum and distal jejunum - increased goblet cells and increased intracellular granules in the Paneth cells (91497, 91498). -Pathology to comment-
Cecum: abundant luminal protozoa (91498).
Salivary glands (submandibular): secretory depletion, ducts (91499).
Kidneys: minimal perivascular/peripelvic mononuclear cell infiltrates (91500).
Lungs: minimal perivascular mononuclear cell infiltrates (91501).
Heart: few foci of cytoplasmic vacuolation of cardiomyocytes, indicative of degeneration (91501).
Heart: myxomatous valvular changes (thickened leaflets) (91501).
Heart: focal cardiomyocyte degeneration/necrosis and associated infiltrates (91501). -Pathology to comment-
Skin: several small foci of epidermal hyperplasia and superficial dermal inflammation (91502).

Spinal cord: skin shows mild and mainly focal subcutaneous polymorphonuclear cell infiltrates with occasional skeletal myocyte degeneration and some necrosis (91506, 91507).
Hind leg: occasional skeletal myocytes show degeneration and necrosis (91508, 91509).
Head: mild focal inflammation (likely folliculitis) (91558).
Head: of the nasal epithelium, mild focal mucosal inflammation with some cytoplasmic hyaline droplet accumulation (91558, 91559), bilateral.

#### 151 Adar1p150-/- Ifih1-/- Eif2ak2-/-; DOB 18/6/2022; female

Macro Observations

Tail suspension test for neurological defects - negative.
Dentition, tongue and oral cavity was unremarkable.
BCS: 3
Spleen: 18x4x2mm
Kidneys: 11x8x6mm, symmetrical
Thymus: 7x7x2mm
Lungs inflated.
Heart: 12x8x6mm
Brain: 15x10x5mm, symmetrical
Pituitary gland identified, macroscopically normal.
Tail length: 80mm (straight)
Head harvested for evaluation of auditory and vestibular structures.
Bone marrow smear taken from left hind leg.

No macroscopic lesions identified.

Micro Observations

Marrow smear: Examination of the smear showed representative cells from the myeloid and erythroid series. Occasional cells from the lymphoid series. Occasional and unremarkable megakaryoblasts.
(91358)

Peripheral blood smear: Examination of the smear showed red blood cells (majority of cells shown), occasional white blood cells including lymphocytes, segmented neutrophils, monocytes and platelets (clumps). No detectable parasites.
Occasional irregular red blood cells showing variation in size (anitocytosis- microcytes) and/or shape (poikilocytosis).
(91359)
-Pathology to comment-

Mammary glands

Typical mammary fat pad with developing lactiferous ducts, blood vessels, and nerve bundles.
(91365, 91369)
No lesions of significance

Ovaries/Oviducts

Unremarkable ovaries containing follicles at various stages of development (primary through to antral) and several corpora lutea.
Unremarkable oviduct micromorphology with typical columnar epithelium and mucosal folds.
(91360)
No lesions of significance

Uterus/Cervix/Vagina/Clitoral gland

Unremarkable architecture of the endometrium/endometrial glands, myometrium and adventitia.
The micromorphology of the uterus and vagina places the animal at metestrus.
(91360)
No lesions of significance

Urinary Bladder

Unremarkable bladder with typical urothelium and detrusor muscle.
(91360)
No lesions of significance

Liver/Gall bladder

Typical liver parenchyma including hepatocytes, Kupffer cells, portal triads and central veins.
Minimal perivascular mixed leukocyte infiltrates, likely incidental – too mild to be significant.
Unremarkable Gall bladder.
(91361)

Stomach

Unremarkable fore and glandular portions of the stomach with limiting ridge.
Section includes duodenal bulb with Brunner's glands.
Minimal scattered polymorphonuclear cell infiltrates in the lamina propria, common incidental finding in mice.
(91362)

Small Intestine (Duodenum, Jejunum & Ileum)/GALT

Discernible mucosal villi and submucosal layers. Unremarkable muscularis and discernible ganglion cells of the plexuses.

Query crypt changes including: epithelial hyperplasia with nuclear crowding, prominent number of mitotic figures, and numerous apoptotic bodies.

Peyer's patches display typical reactive nodal histology. Also identified are smaller aggregates of lymphoid cells (cryptopatches).

(91362, 91363, 91364)

*Comments:*

*Pathology to comment*

Cecum/Colon/GALT

Typical mucosal folds and submucosal layers. Unremarkable muscularis and discernible ganglion cells of the plexuses. Occasional, typical lymphoid cluster (cryptopatches).
Abundant luminal protozoa, not usually clinically significant – feature seen in the control.
(91364)

Mesenteric lymph node

Typical nodal histology with representative cortex including the occasional follicle, an expansive paracortical area, and sinus histiocytosis.
(91360)
No lesions of significance

Spleen

Discernible red and white pulp.
Query decreased white pulp to red pulp ratio; increased extramedullary haematopoiesis.
Occasional haemosiderin laden macrophages in the red pulp.
(91360)

*Comments:*

*Pathology to comment*

Pancreas

Representative exocrine tissue (serous acini) and endocrine tissue (islets of Langerhans).
(91360)
No lesions of significance

Kidney

Section shows a cortex, medulla, and papilla. There is a uniform distribution of glomeruli and accompanying nephron components and the micromorphology of the tubules is unremarkable.
Includes small portion of typical renal lymph node.
(91366)
No lesions of significance

Adrenal glands

Typical cortex/medulla micromorphology.
(91366)
No lesions of significance

Salivary glands and Regional lymph nodes

Unremarkable submandibular, sublingual and parotid glands.
Regional lymph nodes display typical reactive nodal histology.
(91365)
No lesions of significance

Thyroids

Normal lateral lobe of the thyroid gland with typical colloid secreting follicles lined by cuboidal epithelium.
Includes small sheet-like mass of polygonal cells, characteristic of the parathyroid gland.
(91368)
No lesions of significance

Trachea/Lungs

Typical lung parenchyma/alveoli, bronchioles, and blood vessels.
Minimal perivascular mononuclear cell infiltrates, common incidental finding in mice – feature seen in the control.
Small portion of trachea with unremarkable mucosal epithelial lining and hyaline cartilage.
Oesophagus with typical features including stratified squamous epithelium.
(91367, 91368)
No lesions of significance

Thymus

Typical medulla/cortex distribution and micromorphology.
(91367, 91368)
No lesions of significance

*Comments:*

*Pathology to comment*

Heart/chambers/vessels/valves

Representative cardiac muscle, chambers, valves and great vessels of the heart.
The cardiac muscle fibres demonstrated typical features including central nuclei, branching fibres and striations.
(91367, 91368)
No lesions of significance

Skin

Typical dermal appendages and distribution. Unremarkable thin layer of striated muscle (panniculus carnosus).
(91369)
No lesions of significance

Tail

Typical tail components including keratinized squamous epithelium, dense regular connective tissue, tendons, caudal vertebra, bone marrow, intervertebral disc, skeletal muscle, nerves and blood vessels.
(91370)
No lesions of significance

Eyes/Harderian glands

Unremarkable retina, cornea, iris, ciliary body, lens, sclera and choroid.
Typical branched tubuloalveolar formation of the Harderian gland.
Includes portion of unremarkable optic nerve and extraocular muscles.
(91371)
No lesions of significance

Brain

Sections were prepared of the brain:

Level I: including cortex (or cortical canopy), corpus callosum, caudate putamen and lateral ventricles (approx. Bregma N/A)
Level II: including the hippocampus, thalamus, hypothalamus and lateral and third ventricles (approx. Bregma N/A)
Level III: includes the cerebellum, pons and fourth ventricle (approx. Bregma -5.80mm)

Sections of brain appear oblique but symmetrical with unremarkable meninges and typical lamination.
Ventricular dilation observed, query due to plane of section.

The cerebellum appears symmetrical with typical architecture and Purkinje cells.
There was no obvious neuronal loss, and the myelination appears normal.

(91372, 91373)

*Comments:*

*Neuropathology to comment*

Spinal cord

Representative thoracic and lumbar region of spinal cord, vertebral bone, intervertebral disc, striated muscle, peripheral nerves, brown adipose tissue, and bone marrow.
(91374, 91375)
No lesions of significance

*Comments:*

*Neuropathology to comment*

(Hind leg) Long bone/Bone marrow/Synovial joint/Skeletal muscle

Representative long bone, bone marrow, striated muscle, synovial joint, portion of tarsal bones, phalanges, cornified foot pad with discernible eccrine glands.
The skeletal muscle shows consistent fibre size with peripheral nuclei.

Cortical bone changes in the tibia, query plane of section artefact – feature seen in the controls.
Subcutaneous oedematous changes in the distal hind leg and foot, an incidental finding in mice (possibly due to handling) – feature seen in the control.

Focal cortical bone changes (femur) near the knee joint – feature seen in the controls.

(91376, 91377)

Head

Multiple levels through the head demonstrate dermal appendages, nasal cavity with unremarkable nasal epithelium, oral cavity, dentition, and tongue with keratinized papillae. Sections also show unremarkable pituitary gland including pars intermedia, pars distalis and pars nervosa as well and the trigeminal nerve/ganglia (91378). Some mechanical artefact.
The outer and middle regions of the ear are discernible. The tympanic membrane is intact, and the ossicles are unremarkable and include the stapedial annular ligaments.
Typical components of the inner ear including bony labyrinth, organ of corti, stria vascularis and scala cavities are discernible. Based on multiple levels, the organ of corti is unremarkable with no discernible loss of inner/outer hair cells and typical tectorial membrane.
The cochlear nerve and spiral ganglion is also demonstrated and based on several levels, there is no reduction in the density of the spiral ganglion cells. Examples of otolith organs can be seen with typical features such as the hair cells and mineral otoliths. The ampulla including the crista ridge with hair cells is discernible.

(91378, 91379)
No lesions of significance

Sternum

Representative sternebrae, costal cartilage, intersternebral joint, intercostal skeletal muscle and brown fat.
Sections show haematopoietic tissue islands and adipose cells surrounded by vascular sinuses interspersed within a meshwork of trabecular bone. The bone marrow morphology demonstrates typical myeloid features including conspicuous megakaryoblasts and lymphoid features.
(91380)
No lesions of significance

#### 155 Adar1p150-/- Ifih1-/- Eif2ak2-/-; DOB 20/6/2022; female

Macro Observations

Tail suspension test for neurological defects - negative.
Dentition, tongue and oral cavity was unremarkable.
BCS: 3
Spleen: 15x5x3mm
Kidneys: 12x6x5mm, symmetrical
Thymus: 7x7x2mm
Lungs inflated.
Heart: 10x7x5mm
Brain: 15x10x5mm, symmetrical
Pituitary gland identified, macroscopically normal.
Tail length: 80mm (straight)
Head harvested for evaluation of auditory and vestibular structures.
Bone marrow smear taken from left hind leg.

No macroscopic lesions identified.

Micro Observations

Marrow smear: Examination of the smear showed representative cells from the myeloid and erythroid series. Occasional cells from the lymphoid series. Occasional and unremarkable megakaryoblasts.
(91402)

Peripheral blood smear: Examination of the smear showed red blood cells (majority of cells shown), occasional white blood cells including lymphocytes, segmented neutrophils, monocytes and platelets (clumps). No detectable parasites.

Occasional irregular red blood cells showing variation in size (anitocytosis- microcytes) and/or shape (poikilocytosis).
(91403)
-Pathology to comment-

Mammary glands

Typical mammary fat pad with developing lactiferous ducts, blood vessels, and nerve bundles.
(91413)
No lesions of significance

Ovaries/Oviducts

Unremarkable ovaries containing follicles at various stages of development (primary through to antral) and several corpora lutea.
Unremarkable oviduct micromorphology with typical columnar epithelium and mucosal folds.
(91404, 91405)
No lesions of significance

Uterus/Cervix/Vagina/Clitoral gland

Typical architecture of the endometrium/endometrial glands, myometrium and adventitia.
Abundant polymorphonuclear cell infiltrates, appears greater than the level typically seen in the stages of estrus – query inflammatory.
Vaginal stage was not readily apparent.
(91404, 91405)

*Comments:*

*Pathology to comment*

Urinary Bladder

Unremarkable bladder with typical urothelium and detrusor muscle.
(91404, 91405)
No lesions of significance

Liver/Gall bladder

Typical liver parenchyma including hepatocytes, Kupffer cells, portal triads and central veins.
Unremarkable Gall bladder.
(91406)
No lesions of significance

Stomach

Unremarkable fore and glandular portions of the stomach with limiting ridge.
Mild and scattered mixed leukocyte cell infiltrates in the lamina propria and submucosa, common incidental finding in mice.
Includes pyloric sphincter and duodenal bulb with Brunner's glands.
(91407)

Small Intestine (Duodenum, Jejunum & Ileum)/GALT

Discernible mucosal villi and submucosal layers. Unremarkable muscularis and discernible ganglion cells of the plexuses.
Scattered luminal protozoa in the ileum (91409), not usually clinically significant – feature seen in the control.

Query crypt changes including: areas of epithelial hyperplasia with nuclear crowding, prominent number of mitotic figures, and scattered apoptotic bodies.

Increased goblet cells and intracellular granules in the Paneth cells of the distal jejunum/early ileum (91408, 91409) – features seen in the control.
Increased leukocyte infiltrates in the lamina propria of the ileum and distal jejunum (91408, 91409), query inflammatory – feature seen in the control.

Peyer’s patches display typical reactive nodal histology. Also identified are smaller aggregates of lymphoid cells (cryptopatches).

(91407, 91408, 91409)

*Comments:*

*Pathology to comment*

Cecum/Colon/GALT

Typical mucosal folds and submucosal layers. Unremarkable muscularis and discernible ganglion cells of the plexuses. Occasional, typical lymphoid cluster (cryptopatches).
Abundant luminal protozoa, not usually clinically significant – feature seen in the control.
(91409)

Mesenteric lymph node

Typical nodal histology including cortex, medullary cords and sinuses.
(91410)
No lesions of significance

Spleen

Discernible red and white pulp.
Query decreased white pulp to red pulp ratio; increased extramedullary haematopoiesis.
Occasional haemosiderin laden macrophages in the red pulp.
(91404, 91405)

*Comments:*

*Pathology to comment*

Pancreas

Representative exocrine tissue (serous acini) and endocrine tissue (islets of Langerhans).
Irregular focus in the exocrine pancreas: resembles splenic tissue?
(91404, 91405)

*Comments:*

*Pathology to comment*

Kidney

Section shows a cortex, medulla, and papilla. There is a uniform distribution of glomeruli and accompanying nephron components and the micromorphology of the tubules is unremarkable.
(91411)
No lesions of significance

Adrenal glands

Typical cortex/medulla micromorphology.
(91411)
No lesions of significance

Salivary glands and Regional lymph nodes

Unremarkable submandibular, sublingual and parotid glands.
Regional lymph nodes display typical nodal histology.
(91410)
No lesions of significance

Thyroids

Normal lateral lobes of the thyroid gland with typical colloid secreting follicles lined by cuboidal epithelium.
(91412)
No lesions of significance

Trachea/Lungs

Typical lung parenchyma/alveoli, bronchioles, and blood vessels.
Minimal perivascular mononuclear cell infiltrates, common incidental finding in mice – feature seen in the controls.
Trachea with unremarkable mucosal epithelial lining and hyaline cartilage.
Oesophagus with typical features including stratified squamous epithelium.
(91412)
No lesions of significance

Thymus

Typical medulla/cortex distribution and micromorphology.
(91412)
No lesions of significance

*Comments:*

*Pathology to comment*

Heart/chambers/vessels/valves

Representative cardiac muscle, chambers, valves and great vessels of the heart.
Myxomatous valvular changes (thickened leaflets), an incidental finding in mice – feature seen in the controls.
The cardiac muscle fibres demonstrated typical features including central nuclei, branching fibres and striations.
(91412)
No lesions of significance

Skin

Typical dermal appendages and distribution. Unremarkable thin layer of striated muscle (panniculus carnosus).
(91413)
No lesions of significance

Tail

Typical tail components including keratinized squamous epithelium, dense regular connective tissue, tendons, caudal vertebra, bone marrow, intervertebral disc, skeletal muscle, nerves and blood vessels.
(91414)
No lesions of significance

Eyes/Harderian glands

Unremarkable retina, cornea, iris, ciliary body, lens, sclera and choroid.
Typical branched tubuloalveolar formation of the Harderian gland.
Includes portion of unremarkable optic nerve and extraocular muscles.
(91415)
No lesions of significance

Brain

Sections were prepared from the standard levels of the brain:

Level I: including cortex (or cortical canopy), corpus callosum, caudate putamen and lateral ventricles (approx. Bregma 0.70mm)
Level II: including the hippocampus, thalamus, hypothalamus and lateral and third ventricles (approx. Bregma -2.18mm)
Level III: includes the cerebellum, pons and fourth ventricle (approx. Bregma -5.34mm)

Sections of brain appear symmetrical with no ventricular dilation observed, unremarkable meninges and typical lamination.
The cerebellum appears symmetrical with typical architecture and Purkinje cells.
There was no obvious neuronal loss, and the myelination appears normal.
(91416)
No lesions of significance

*Comments:*

*Neuropathology to comment*

Spinal cord

Representative thoracic and lumbar region of spinal cord, vertebral bone, intervertebral disc, striated muscle, peripheral nerves, brown adipose tissue, and bone marrow.
(91417, 91418)
No lesions of significance

*Comments:*

*Neuropathology to comment*

(Hind leg) Long bone/Bone marrow/Synovial joint/Skeletal muscle

Unremarkable long bone, bone marrow, striated muscle, synovial joint, portion of tarsal bones, phalanges, cornified foot pad with discernible eccrine glands.
The skeletal muscle shows consistent fibre size with peripheral nuclei.
(91419)
No lesions of significance

Head

Multiple levels through the head demonstrate dermal appendages, nasal cavity with unremarkable nasal epithelium, oral cavity, dentition, and tongue with keratinized papillae.
Sections also show unremarkable pituitary gland including pars intermedia, pars distalis and pars nervosa as well and the trigeminal nerve/ganglia (91420).
The outer and middle regions of the ear are discernible. The tympanic membrane is intact, and the ossicles are unremarkable and include the stapedial annular ligaments.
Typical components of the inner ear including bony labyrinth, organ of corti, stria vascularis and scala cavities are discernible. Based on multiple levels, the organ of corti is unremarkable with no discernible loss of inner/outer hair cells and typical tectorial membrane.
The cochlear nerve and spiral ganglion is also demonstrated and based on several levels, there is no reduction in the density of the spiral ganglion cells. Examples of otolith organs can be seen with typical features such as the hair cells and mineral otoliths. The ampulla including the crista ridge with hair cells is discernible (91421).

(91420, 91421)
No lesions of significance

Sternum

Representative sternebrae, costal cartilage, intersternebral joint, intercostal skeletal muscle and brown fat.
Sections show haematopoietic tissue islands and adipose cells surrounded by vascular sinuses interspersed within a meshwork of trabecular bone. The bone marrow morphology demonstrates

typical myeloid features including conspicuous megakaryoblasts and lymphoid features.
(91422)
No lesions of significance

#### 167 Adar1p150-/- Ifih1-/- Eif2ak2-/-; DOB 16/7/2022; male

Macro Observations

Tail suspension test for neurological defects - negative.
Dentition, tongue and oral cavity was unremarkable.
BCS: 4
Testes: 7x5x3mm, symmetrical
Spleen: 15x4x2mm
Kidneys: 12x5x5mm, symmetrical
Thymus: 7x7x2mm
Lungs inflated.
Heart: 11x7x5mm
Brain: 15x10x5mm, symmetrical
Pituitary gland identified, macroscopically normal.
Tail length: 80mm (straight)
Head harvested for evaluation of auditory and vestibular structures.
Bone marrow smear taken from left hind leg.

No macroscopic lesions identified.

Micro Observations

Marrow smear: Examination of the smear showed representative cells from the myeloid and erythroid series. Occasional cells from the lymphoid series. Occasional and unremarkable megakaryoblasts.
(91423)

Peripheral blood smear: Examination of the smear showed red blood cells (majority of cells shown), occasional white blood cells including lymphocytes, scarce neutrophils, monocytes and platelets (clumps). No detectable parasites.
Occasional irregular red blood cells showing variation in size (anitocytosis- microcytes) and/or shape (poikilocytosis).
(91424)
-Pathology to comment-

Testes/Epididymes

Typical convoluted seminiferous tubules at various stages of cycle surrounded by the tunica albuginea. Within the tubules, unremarkable spermatogenic cells including, Sertoli cells, spermatogonia, developing spermatocytes and spermatids. Typical interstitial Leydig cells.
The architecture of the epididymis is typical, with numerous intraluminal elongated spermatozoa.
(91425)
No lesions of significance

Seminal vesicles

Unremarkable tall columnar epithelium and folded mucosa.
Presence of typical intraluminal eosinophilic secretions.
(91426, 91427)
No lesions of significance

Prostate glands

Unremarkable dorsal lateral/ventral/coagulating glands with typical intraluminal secretions.
Includes portion of unremarkable vas deferens.
(91426, 91427)
No lesions of significance

Penis/Preputial gland

Typical penile structures including prepuce, epithelium of the glans, corpus cavernosum and urethra.
Typical preputial glands including basal and secretory cells.
(91428)
No lesions of significance

Urinary Bladder

Unremarkable bladder with typical urothelium and detrusor muscle.
(91426, 91427)
No lesions of significance

Liver/Gall bladder

Typical liver parenchyma including hepatocytes, Kupffer cells, portal triads and central veins.
Single small parenchymal leukocyte aggregate with some hepatocyte cell loss/necrosis, common incidental/background finding – feature seen in the control.
Unremarkable Gall bladder.
(91429)
No lesions of significance

Stomach

Unremarkable fore and glandular portions of the stomach with limiting ridge.
Single dilated gland, common incidental finding.
Includes pyloric sphincter and duodenal bulb with Brunner's glands.
(91430)
No lesions of significance

Small Intestine (Duodenum, Jejunum & Ileum)/GALT

Discernible mucosal villi and submucosal layers. Unremarkable muscularis and discernible ganglion cells of the plexuses.
Singular cryptolysis in the duodenum (91430), an incidental finding in mice.

Mainly in the jejunum, query crypt changes including: epithelial hyperplasia with nuclear crowding, prominent number of mitotic figures, and numerous apoptotic bodies.

Peyer’s patches display typical reactive nodal histology. Also identified are smaller aggregates of lymphoid cells (cryptopatches).

(91430, 91431, 91432)

*Comments:*

*Pathology to comment*

Cecum/Colon/GALT

Typical mucosal folds and submucosal layers. Unremarkable muscularis and discernible ganglion cells of the plexuses. Occasional, typical lymphoid cluster (cryptopatches).
Abundant luminal protozoa, not usually clinically significant – feature seen in the control.
(91432)

Mesenteric lymph node

Typical reactive micromorphology including follicular hyperplasia, germinal centre formation, an expansive paracortical area, and sinus histiocytosis.
(91433)
No lesions of significance

Spleen

Discernible red and white pulp.
Query decreased white pulp to red pulp ratio; increased extramedullary haematopoiesis.
Occasional haemosiderin laden macrophages in the red pulp.
(91426, 91427)

*Comments:*

*Pathology to comment*

Pancreas

Representative exocrine tissue (serous acini) and endocrine tissue (islets of Langerhans).
(91426, 91427)
No lesions of significance

Kidney

Section shows a cortex, medulla, and papilla. There is a uniform distribution of glomeruli and accompanying nephron components and the micromorphology of the tubules is unremarkable.
Includes renal lymph nodes with typical nodal histology.
(91434)
No lesions of significance

Adrenal glands

An adrenal gland showing typical cortex/medulla micromorphology.
(91434)
No lesions of significance

Salivary glands and Regional lymph nodes

Unremarkable submandibular, sublingual and parotid glands.
Regional lymph nodes display typical nodal histology.
(91433)
No lesions of significance

Thyroids

Normal lateral lobe of the thyroid gland with typical colloid secreting follicles lined by cuboidal epithelium.
(91435)
No lesions of significance

Trachea/Lungs

Typical lung parenchyma/alveoli, bronchioles, blood vessels and parabronchial lymph nodes.
Minimal perivascular mononuclear cell infiltrates, incidental finding in mice – feature seen in the controls
Trachea with unremarkable mucosal epithelial lining and hyaline cartilage.
Oesophagus with typical features including stratified squamous epithelium.
(91435)
No lesions of significance

Thymus

Typical medulla/cortex distribution and micromorphology.
(91435)
No lesions of significance

*Comments:*

*Pathology to comment*

Heart/chambers/vessels/valves

Representative cardiac muscle, chambers, valves and great vessels of the heart.
Myxomatous valvular changes (thickened leaflets), an incidental finding in mice – feature seen in the controls.
The cardiac muscle fibres demonstrated typical features including central nuclei, branching fibres and striations.
(91435)
No lesions of significance

Skin

Typical dermal appendages and distribution. Unremarkable thin layer of striated muscle (panniculus carnosus).
(91436)
No lesions of significance

Tail

Typical tail components including keratinized squamous epithelium, dense regular connective tissue, tendons, caudal vertebra, bone marrow, intervertebral disc, skeletal muscle, nerves and blood vessels.
(91437)
No lesions of significance

Eyes/Harderian glands

Unremarkable retina, cornea, iris, ciliary body, lens, sclera and choroid.
Typical branched tubuloalveolar formation of the Harderian gland.
Includes portion of unremarkable optic nerve and extraocular muscles.
(91438)
No lesions of significance

Brain

Sections were prepared from the standard levels of the brain:

Level I: including cortex (or cortical canopy), corpus callosum, caudate putamen and lateral ventricles (approx. Bregma 0.50mm)
Level II: including the hippocampus, thalamus, hypothalamus and lateral and third ventricles (approx. Bregma N/A)
Level III: includes the cerebellum, pons and fourth ventricle (approx. Bregma -5.68mm)

Sections of brain appear symmetrical with no ventricular dilation observed, unremarkable meninges and typical lamination.
The cerebellum appears symmetrical with typical architecture and Purkinje cells.
There was no obvious neuronal loss, and the myelination appears normal.
(91439)
No lesions of significance

*Comments:*

*Neuropathology to comment*

Spinal cord

Representative thoracic and lumbar region of spinal cord, vertebral bone, intervertebral disc, striated muscle, peripheral nerves, brown adipose tissue, and bone marrow.
(91440, 91441)
No lesions of significance

*Comments:*

*Neuropathology to comment*

(Hind leg) Long bone/Bone marrow/Synovial joint/Skeletal muscle

Unremarkable long bone, bone marrow, striated muscle, synovial joint, portion of tarsal bones, phalanges, cornified foot pad with discernible eccrine glands.
The skeletal muscle shows consistent fibre size with peripheral nuclei.
(91442, 91443)
No lesions of significance

Head

Multiple levels through the head demonstrate dermal appendages, nasal cavity with unremarkable nasal epithelium, oral cavity, dentition, and tongue with keratinized papillae.
Sections also show unremarkable pituitary gland including pars intermedia, pars distalis and pars nervosa as well and the trigeminal nerve/ganglia (91444).
The outer and middle regions of the ear are discernible. The tympanic membrane is intact, and the ossicles are unremarkable and include the stapedial annular ligaments.
Typical components of the inner ear including bony labyrinth, organ of corti, stria vascularis and scala cavities are discernible. Based on multiple levels, the organ of corti is unremarkable with no discernible loss of inner/outer hair cells and typical tectorial membrane.
The cochlear nerve and spiral ganglion is also demonstrated and based on several levels, there is no reduction in the density of the spiral ganglion cells. Examples of otolith organs can be seen with typical features such as the hair cells and mineral otoliths. The ampulla including the crista ridge with hair cells is discernible (91445).

(91444, 91445)
No lesions of significance

Sternum

Representative sternebrae, costal cartilage, intersternebral joint, intercostal skeletal muscle and brown fat.
Several skeletal myocytes show degeneration/necrosis and associated infiltrates, likely incidental.

Sections also show haematopoietic tissue islands and adipose cells surrounded by vascular sinuses interspersed within a meshwork of trabecular bone. The bone marrow morphology demonstrates typical myeloid features including conspicuous megakaryoblasts and lymphoid features.

(91446)

*Comments:*

*Pathology to comment*

#### 492 Adar1-/- Ifih1-/- Eif2ak2-/-; DOB 3/6/2022; female

Macro Observations

Tail suspension test for neurological defects - negative.
Dentition, tongue and oral cavity was unremarkable.
BCS: 3
Spleen: 14x5x2mm
Kidneys: 11x6x4mm, symmetrical
Thymus: 9x9x2mm, large.
Lungs inflated.
Heart: 10x8x7mm
Brain: 15x10x5mm, symmetrical
Pituitary gland identified, macroscopically normal.
Tail length: 80mm (straight)
Head harvested for evaluation of auditory and vestibular structures.
Bone marrow smear taken from left hind leg.

No macroscopic lesions identified.

Micro Observations

Marrow smear: Examination of the smear showed representative cells from the myeloid and erythroid series. Occasional cells from the lymphoid series. Occasional and unremarkable megakaryoblasts.
(91468)

Peripheral blood smear: Examination of the smear showed red blood cells (majority of cells shown), occasional white blood cells including lymphocytes, few segmented neutrophils, monocytes and platelets (clumps). No detectable parasites.
Occasional irregular red blood cells showing variation in size (anitocytosis- microcytes) and/or shape (poikilocytosis).
Of the few neutrophils seen, some appear immature (banded).
(91469)
-Pathology to comment-

Mammary glands

Typical mammary fat pad with developing lactiferous ducts, blood vessels, and nerve bundles.
(91476)
No lesions of significance

Ovaries/Oviducts

Unremarkable ovaries containing follicles at various stages of development (primary through to antral) and several corpora lutea.
Unremarkable oviduct micromorphology with typical columnar epithelium and mucosal folds.
(91470, 91471)
No lesions of significance

Uterus/Cervix/Vagina/Clitoral gland

Unremarkable architecture of the endometrium/endometrial glands, myometrium and adventitia.
The micromorphology of the uterus and vagina places the animal at metestrus/diestrus.
(91470, 91471)
No lesions of significance

Urinary Bladder

Unremarkable bladder with typical urothelium and detrusor muscle.
(91470, 91471)
No lesions of significance

Liver/Gall bladder

Mostly typical liver parenchyma including hepatocytes, Kupffer cells, portal triads and central veins.
Scattered foci of hepatocyte vacuolation resembling hydropic degeneration or fatty change (steatosis), likely incidental/age-related change – feature seen in the controls.
Multiple small parenchymal and perivascular leukocyte infiltrates with some hepatocyte cell loss/necrosis, common incidental/background finding – features seen in the controls.
Gall bladder shows segment of epithelial hyperplasia and polymorphonuclear cell infiltration in the underlying lamina propria, a common incidental finding – feature seen in the controls.
(91472)

Stomach

Unremarkable fore and glandular portions of the stomach with limiting ridge.
Includes duodenal bulb with Brunner's glands.
(91473)
No lesions of significance

Small Intestine (Duodenum, Jejunum & Ileum)/GALT

Discernible mucosal villi and submucosal layers. Unremarkable muscularis and discernible ganglion cells of the plexuses.

Query crypt changes including: some areas of epithelial hyperplasia with nuclear crowding, prominent number of mitotic figures, and numerous apoptotic bodies.

Peyer's patches display typical reactive nodal histology. Also identified are smaller aggregates of lymphoid cells (cryptopatches).

(91473, 91474, 91475)

*Comments:*

*Pathology to comment*

Cecum/Colon/GALT

Typical mucosal folds and submucosal layers. Unremarkable muscularis and discernible ganglion cells of the plexuses. Occasional, typical lymphoid cluster (cryptopatches).
Abundant luminal protozoa in the cecum and proximal colon, not usually clinically significant – feature seen in the control.
(91475)

Mesenteric lymph node

Typical reactive micromorphology including follicular hyperplasia, germinal centre formation, an expansive paracortical area, and sinus histiocytosis.
(91476)
No lesions of significance

Spleen

Discernible red and white pulp.
Query decreased white pulp to red pulp ratio; increased extramedullary haematopoiesis.
Occasional haemosiderin laden macrophages in the red pulp.
(91470, 91471)

*Comments:*

*Pathology to comment*

Pancreas

Representative exocrine tissue (serous acini) and endocrine tissue (islets of Langerhans).
(91470, 91471)
No lesions of significance

Kidney

Section shows a cortex, medulla, and papilla. There is a uniform distribution of glomeruli and accompanying nephron components and the micromorphology of the tubules is unremarkable.
Multiple small protein casts within the medulla, an incidental finding in mice – feature seen in the controls.
Section does not includes renal lymph nodes.
(91477)
No lesions of significance

Adrenal glands

Typical cortex/medulla micromorphology.
(91477)
No lesions of significance

Salivary glands and Regional lymph nodes

Unremarkable submandibular, sublingual and parotid glands.
Regional lymph nodes display nodal histology.
(91476)
No lesions of significance

Thyroids

Normal lateral lobes of the thyroid gland with typical colloid secreting follicles lined by cuboidal epithelium.
(91478)
No lesions of significance

Trachea/Lungs

Typical lung parenchyma/alveoli, bronchioles, blood vessels and parabronchial lymph node.
Minimal perivascular mononuclear cell infiltrates, common incidental finding – feature seen in the controls.
Trachea with unremarkable mucosal epithelial lining and hyaline cartilage.
Oesophagus with typical features including stratified squamous epithelium.
(91478)
No lesions of significance

Thymus

Typical medulla/cortex distribution and micromorphology.
(91478)
No lesions of significance

*Comments:*

*Pathology to comment*

Heart/chambers/vessels/valves

Representative cardiac muscle, chambers, valves and great vessels of the heart.
The cardiac muscle fibres demonstrated typical features including central nuclei, branching fibres and striations.
Focal cardiomyocyte degeneration/necrosis – feature seen in the controls.
Query focal changes to a vessel wall, possible plane of section artefact.
Occasional vacuolation of cardiomyocytes, indicative of degeneration – feature seen in the controls.
(91478)

*Comments:*

*Pathology to comment*

Skin

Typical dermal appendages and distribution. Unremarkable thin layer of striated muscle (panniculus carnosus).
(91479)
No lesions of significance

Tail

Typical tail components including keratinized squamous epithelium, dense regular connective tissue, tendons, caudal vertebra, bone marrow, intervertebral disc, skeletal muscle, nerves and blood vessels.
(91480)
No lesions of significance

Eyes/Harderian glands

Unremarkable retina, cornea, iris, ciliary body, lens, sclera and choroid.
Typical branched tubuloalveolar formation of the Harderian gland.
Includes portion of unremarkable optic nerve and extraocular muscles.
(91481)
No lesions of significance

Brain

Sections were prepared from the standard levels of the brain:

Level I: including cortex (or cortical canopy), corpus callosum, caudate putamen and lateral ventricles (approx. Bregma 0.98mm)
Level II: including the hippocampus, thalamus, hypothalamus and lateral and third ventricles (approx. Bregma -1.34mm)
Level III: includes the cerebellum, pons and fourth ventricle (approx. Bregma -5.20mm)

Sections of brain appear symmetrical with no ventricular dilation observed, unremarkable meninges and typical lamination.
The cerebellum appears symmetrical with typical architecture and Purkinje cells.
There was no obvious neuronal loss, and the myelination appears normal.
(91482)
No lesions of significance

*Comments:*

*Neuropathology to comment*

Spinal cord

Representative thoracic and lumbar region of spinal cord, vertebral bone, intervertebral disc, striated muscle, peripheral nerves, brown adipose tissue, and bone marrow.
Subcutaneous oedematous changes, possible incidental finding.
(91483, 91484)

*Comments:*

*Neuropathology to comment*

(Hind leg) Long bone/Bone marrow/Synovial joint/Skeletal muscle

Unremarkable long bone, bone marrow, striated muscle, synovial joint, portion of tarsal bones, phalanges, cornified foot pad with discernible eccrine glands.
The skeletal muscle shows consistent fibre size with peripheral nuclei.
(91485, 91486)
No lesions of significance

Head

Multiple levels through the head demonstrate dermal appendages, nasal cavity with unremarkable nasal epithelium, oral cavity, dentition, and tongue with keratinized papillae.
Sections also show unremarkable pituitary gland including pars intermedia, pars distalis and pars nervosa as well and the trigeminal nerve/ganglia (91487).
The outer and middle regions of the ear are discernible. The tympanic membrane is intact, and the ossicles are unremarkable and include the stapedial annular ligaments.
Typical components of the inner ear including bony labyrinth, organ of corti, stria vascularis and scala cavities are discernible. Based on multiple levels, the organ of corti is unremarkable with no discernible loss of inner/outer hair cells and typical tectorial membrane.
The cochlear nerve and spiral ganglion is also demonstrated and based on several levels, there is no reduction in the density of the spiral ganglion cells. Examples of otolith organs can be seen with typical features such as the hair cells and mineral otoliths. The ampulla including the crista ridge with hair cells is discernible (91488).

Middle ear inflammation (otitis media), unilateral - mucosal oedema with small foci of

mineralisation, and some luminal macrophages/cellular debris – an incidental finding in mice.
Focal inflammation of one hair follicle (folliculitis) (91488), common incidental finding in mice.

(91487, 91488)

Sternum

Representative sternebrae, costal cartilage, intersternebral joint, intercostal skeletal muscle and brown fat.
Sections show haematopoietic tissue islands and adipose cells surrounded by vascular sinuses interspersed within a meshwork of trabecular bone. The bone marrow morphology demonstrates typical myeloid features including conspicuous megakaryoblasts and lymphoid features.
(91489)
No lesions of significance

#### 500 Adar1-/- Ifih1-/- Eif2ak2-/-; DOB 10/6/2022; male

Macro Observations

Tail suspension test for neurological defects - negative.
Dentition, tongue and oral cavity was unremarkable.
BCS: 3
Testes: 8x5x5mm, symmetrical
Spleen: 12x5x2mm
Kidneys: 12x8x6mm, symmetrical
Thymus: 7x7x2mm
Lungs inflated.
Heart: 11x10x6mm
Brain: 16x10x5mm, symmetrical
Pituitary gland identified, macroscopically normal.
Tail length: 80mm (straight)
Head harvested for evaluation of auditory and vestibular structures.
Bone marrow smear taken from left hind leg.

No macroscopic lesions identified.

Micro Observations

Marrow smear: Examination of the smear showed representative cells from the myeloid and erythroid series. Occasional cells from the lymphoid series. Occasional and unremarkable megakaryoblasts.
(91512)

Peripheral blood smear: Examination of the smear showed red blood cells (majority of cells shown), occasional white blood cells including lymphocytes, segmented neutrophils, monocytes and platelets (clumps). No detectable parasites.
Occasional irregular red blood cells showing variation in size (anitocytosis- microcytes) and/or shape (poikilocytosis).
(91511)
-Pathology to comment-

Testes/Epididymes

Typical convoluted seminiferous tubules at various stages of cycle surrounded by the tunica albuginea. Within the tubules, unremarkable spermatogenic cells including, Sertoli cells, spermatogonia, developing spermatocytes and spermatids. Typical interstitial Leydig cells.
The architecture of the epididymis is typical, with numerous intraluminal elongated spermatozoa.
(91513)
No lesions of significance

Seminal vesicles

Unremarkable tall columnar epithelium and folded mucosa.
Presence of typical intraluminal eosinophilic secretions.
(91514, 91515)
No lesions of significance

Prostate glands

Unremarkable dorsal lateral/ventral/coagulating glands with typical intraluminal secretions.
Includes portion of unremarkable vas deferens.
(91514, 91515)
No lesions of significance

Penis/Preputial gland

Typical penile structures including prepuce, epithelium of the glans, corpus cavernosum and urethra.
Typical preputial glands including basal and secretory cells.
Surrounding skin shows mild focal accumulation of dermal melanocytes, common incidental/background finding in mice.
(91516)
No lesions of significance

Urinary Bladder

Unremarkable bladder with typical urothelium and detrusor muscle.
(91514, 91515)
No lesions of significance

Liver/Gall bladder

Mostly typical liver parenchyma including hepatocytes, Kupffer cells, portal triads and central veins.
Mainly focal hepatocyte necrosis with some polymorphonuclear cell infiltration, an incidental finding in mice.
Gall bladder shows mild epithelial hyperplasia and polymorphonuclear cell infiltration in the underlying lamina propria, common incidental finding – feature seen in the controls.
(91517)

*Comments:*

*Pathology to comment*

Stomach

Unremarkable fore and glandular portions of the stomach with limiting ridge.
Includes pyloric sphincter and duodenal bulb with Brunner's glands.
(91518)
No lesions of significance

Small Intestine (Duodenum, Jejunum & Ileum)/GALT

Discernible mucosal villi and submucosal layers. Unremarkable muscularis and discernible ganglion cells of the plexuses.

Query crypt changes including: some areas of epithelial hyperplasia with nuclear crowding, prominent number of mitotic figures, and areas of scattered apoptotic bodies.

Peyer’s patches display typical reactive nodal histology. Also identified are smaller aggregates of lymphoid cells (cryptopatches).

(91518, 91519, 91520)

*Comments:*

*Pathology to comment*

Cecum/Colon/GALT

Typical mucosal folds and submucosal layers. Unremarkable muscularis and discernible ganglion cells of the plexuses. Occasional, typical lymphoid cluster (cryptopatches).
Abundant luminal protozoa in the cecum, not usually clinically significant – feature seen in the control.
(91520

Mesenteric lymph node

Typical reactive micromorphology including follicular hyperplasia, germinal centre formation, an expansive paracortical area, and sinus histiocytosis.
(91514, 91515)
No lesions of significance

Spleen

Readily discernible red and white pulp.
White pulp appears largely comparable to the control.
Increased extramedullary haematopoiesis and scattered haemosiderin laden macrophages in the red pulp.
(91514, 91515)

*Comments:*

*Pathology to comment*

Pancreas

Representative exocrine tissue (serous acini) and endocrine tissue (islets of Langerhans).
Minimal focal perivascular mononuclear cell infiltrate (91514), likely incidental/age-related – too mild to be significant.
(91514, 91515)

Kidney

Section shows a cortex, medulla, and papilla. There is a uniform distribution of glomeruli and accompanying nephron components and the micromorphology of the tubules is unremarkable.
Few protein casts within the medulla and papilla, incidental finding – feature seen in the controls.
Includes renal lymph nodes with typical nodal histology.
(91521)
No lesions of significance

Adrenal glands

Sections do not include adrenal glands.

Salivary glands and Regional lymph nodes

Unremarkable submandibular, sublingual and parotid glands.
Secretory depletion (ducts) in the submandibular gland – feature seen in the control.
Regional lymph nodes display typical nodal histology.
(91514, 91515)

Thyroids

Portion of thyroid gland with typical colloid secreting follicles lined by cuboidal epithelium.
(91522)
No lesions of significance

Trachea/Lungs

Typical lung parenchyma/alveoli, bronchioles, and blood vessels.
Minimal perivascular mononuclear cell infiltrate, an incidental/age-related change – feature seen in the controls.
Oesophagus with typical features including stratified squamous epithelium.
Sections do not include trachea.
(91522)
No lesions of significance

Thymus

Typical medulla/cortex distribution and micromorphology.
(91522)
No lesions of significance

*Comments:*

*Pathology to comment*

Heart/chambers/vessels/valves

Representative cardiac muscle, chambers, valves and great vessels of the heart.
Scattered foci of cytoplasmic vacuolation of cardiomyocytes, indicative of degeneration – feature seen in the controls.
Remaining cardiac muscle fibres demonstrated typical features including central nuclei, branching fibres and striations.
Myxomatous valvular changes (thickened leaflet), an incidental finding in mice – feature seen in the controls.
(91522)

Skin

Typical dermal appendages and distribution. Unremarkable thin layer of striated muscle (panniculus carnosus).
Mild focal subcutaneous inflammation, common incidental finding in mice – similar feature seen in the controls.
(91523)

Tail

Typical tail components including keratinized squamous epithelium, dense regular connective tissue, tendons, caudal vertebra, bone marrow, intervertebral disc, skeletal muscle, nerves and blood vessels.
Focal myocyte degeneration/necrosis and associated infiltrates, an incidental finding in mice.
(91524)

Eyes/Harderian glands

Unremarkable retina, cornea, iris, ciliary body, lens, sclera and choroid.
Typical branched tubuloalveolar formation of the Harderian gland.
Includes portion of unremarkable optic nerve and extraocular muscles.
(91525)
No lesions of significance

Brain

Sections were prepared from the standard levels of the brain:

Level I: including cortex (or cortical canopy), corpus callosum, caudate putamen and lateral ventricles (approx. Bregma 0.98mm)
Level II: including the hippocampus, thalamus, hypothalamus and lateral and third ventricles (approx. Bregma -1.70mm)
Level III: includes the cerebellum, pons and fourth ventricle (approx. Bregma N/A)

Sections of brain appear symmetrical with no ventricular dilation observed, unremarkable meninges and typical lamination.
The cerebellum appears symmetrical with typical architecture and Purkinje cells.
There was no obvious neuronal loss, and the myelination appears normal.
(91526)
No lesions of significance

*Comments:*

*Neuropathology to comment*

Spinal cord

Representative thoracic and lumbar region of spinal cord, vertebral bone, intervertebral disc, striated muscle, peripheral nerves, brown adipose tissue, and bone marrow.
Small cluster of myocytes show degeneration/necrosis and associated polymorphonuclear cell infiltrates (91527), likely incidental.
(91527, 91528)

*Comments:*

*Neuropathology to comment*

(Hind leg) Long bone/Bone marrow/Synovial joint/Skeletal muscle

Representative long bone, bone marrow, striated muscle, synovial joint, portion of tarsal bones, phalanges, cornified foot pad with discernible eccrine glands.
The skeletal muscle shows consistent fibre size with peripheral nuclei.

Cortical bone changes (tibia), query due to plane of section (91529).
Focal cortical bone changes (femur) near the knee joint – feature seen in the controls.
Mild focal subcutaneous inflammation in the foot pad (91530), common incidental finding in mice.

(91529, 91530)

*Comments:*

*Pathology to comment*

Head

Multiple levels through the head demonstrate dermal appendages, nasal cavity with unremarkable nasal epithelium, oral cavity, dentition, and tongue with keratinized papillae.
Sections also show unremarkable pituitary gland including pars intermedia, pars distalis and pars nervosa as well and the trigeminal nerve/ganglia (91531).
The outer and middle regions of the ear are discernible. The tympanic membrane is intact, and the ossicles are unremarkable and include the stapedial annular ligaments.
Typical components of the inner ear including bony labyrinth, organ of corti, stria vascularis and scala cavities are discernible. Based on multiple levels, the organ of corti is unremarkable with no discernible loss of inner/outer hair cells and typical tectorial membrane.
The cochlear nerve and spiral ganglion is also demonstrated and based on several levels, there is no reduction in the density of the spiral ganglion cells. Examples of otolith organs can be seen with typical features such as the hair cells and mineral otoliths. The ampulla including the crista ridge with hair cells is discernible (91532).

Of the nasal epithelium, mild cytoplasmic hyaline droplet accumulation, bilateral, an incidental finding – feature seen in the controls.
Mild to moderate mixed leukocyte inflammation surrounding the hair follicles near the oral cavity, bilateral (91532), an incidental finding in mice.

(91531, 91532)

Sternum

Representative sternebrae, costal cartilage, intersternebral joint, intercostal skeletal muscle and brown fat.
Sections show haematopoietic tissue islands and adipose cells surrounded by vascular sinuses interspersed within a meshwork of trabecular bone. The bone marrow morphology demonstrates typical myeloid features including conspicuous megakaryoblasts and lymphoid features.
(91533)
No lesions of significance

#### 513 Adar1-/- Ifih1-/- Eif2ak2-/-; DOB 13/6/2022; male

Macro Observations

Tail suspension test for neurological defects - negative.
Dentition, tongue and oral cavity was unremarkable.
BCS: 3
Testes: 7x5x4mm, symmetrical
Spleen: 14x5x2mm
Kidneys: 12x5x5mm, symmetrical
Thymus: 7x7x2mm
Lungs partially inflated.
Heart: 10x7x6mm
Brain: 15x10x5mm, symmetrical
Pituitary gland identified, macroscopically normal.
Tail length: 85mm (straight)
Head harvested for evaluation of auditory and vestibular structures.
Bone marrow smear taken from left hind leg.

No macroscopic lesions identified.

Micro Observations

Marrow smear: Examination of the smear showed representative cells from the myeloid and erythroid series. Occasional cells from the lymphoid series. A single unremarkable megakaryoblast identified.
(91534)

Peripheral blood smear: Examination of the smear showed red blood cells (majority of cells shown), occasional white blood cells including lymphocytes, segmented neutrophils, monocytes and platelets (clumps). No detectable parasites.
Occasional irregular red blood cells showing variation in size (anitocytosis- microcytes) and/or shape (poikilocytosis).
(91535)
-Pathology to comment-

Testes/Epididymes

Typical convoluted seminiferous tubules at various stages of cycle surrounded by the tunica albuginea. Within the tubules, unremarkable spermatogenic cells including, Sertoli cells, spermatogonia, developing spermatocytes and spermatids. Typical interstitial Leydig cells.
Single tubule shows testicular degeneration/atrophy, an incidental/age-related finding in mice – too mild to be significant.
Portion of unremarkable vas deferens with some typical intraluminal sperm.
Mostly typical epididymis with numerous intraluminal elongated spermatozoa. Mild focal and mainly polymorphonuclear cell inflammation in the loose connective tissue, possible incidental finding.
(91536)

Seminal vesicles

Unremarkable tall columnar epithelium and folded mucosa.
Presence of typical intraluminal eosinophilic secretions.
(91537, 91538)
No lesions of significance

Prostate glands

Unremarkable dorsal lateral/ventral/coagulating glands with typical intraluminal secretions.
Includes portion of unremarkable vas deferens.
(91537, 91538)
No lesions of significance

Penis/Preputial gland

Typical penile structures including prepuce, epithelium of the glans, corpus cavernosum and urethra.
Typical preputial glands including basal and secretory cells.
(91539)
No lesions of significance

Urinary Bladder

Unremarkable bladder with typical urothelium and detrusor muscle.
(91537, 91538)
No lesions of significance

Liver/Gall bladder

Typical liver parenchyma including hepatocytes, Kupffer cells, portal triads and central veins.
Gall bladder shows mild epithelial hyperplasia and polymorphonuclear cell infiltration in the underlying lamina propria, common incidental finding – feature seen in the controls.
(91540)
No lesions of significance

Stomach

Unremarkable fore and glandular portions of the stomach with limiting ridge.
Of the glandular portion, minimal submucosal mixed leukocyte infiltrates, common incidental change – feature seen in the controls
Includes duodenal bulb with Brunner's glands.
(91541)
No lesions of significance

Small Intestine (Duodenum, Jejunum & Ileum)/GALT

Discernible mucosal villi and submucosal layers. Unremarkable muscularis and discernible ganglion cells of the plexuses.
Single dilated crypt containing luminal cellular debris (91542), likely incidental.

Query crypt changes including: some areas of epithelial hyperplasia with nuclear crowding, prominent number of mitotic figures, and numerous apoptotic bodies.

Peyer’s patches display typical reactive nodal histology. Also identified are smaller aggregates of lymphoid cells (cryptopatches).

(91541, 91542, 91543)

*Comments:*

*Pathology to comment*

Cecum/Colon/GALT

Typical mucosal folds and submucosal layers. Unremarkable muscularis and discernible ganglion cells of the plexuses. Occasional, typical lymphoid cluster (cryptopatches).
Abundant luminal protozoa in the cecum and proximal colon, not usually clinically significant – feature seen in the control.
(91543)

Mesenteric lymph node

Representative cortex including the occasional follicle, an expansive paracortical area with lymphoid apoptosis, and sinus histiocytosis.
(91544)
No lesions of significance

Spleen

Discernible red and white pulp.
Noticeable decrease in white pulp to red pulp ratio. Increased extramedullary haematopoiesis.
Occasional haemosiderin laden macrophages in the red pulp.
(91537, 91538)

*Comments:*

*Pathology to comment*

Pancreas

Representative exocrine tissue (serous acini) and endocrine tissue (islets of Langerhans).
(91537, 91538)
No lesions of significance

Kidney

Section shows a cortex, medulla, and papilla. There is a uniform distribution of glomeruli and accompanying nephron components and the micromorphology of the tubules is unremarkable.
Includes renal lymph node with typical nodal histology.
(91545)
No lesions of significance

Adrenal glands

Typical cortex/medulla micromorphology.
(91545)
No lesions of significance

Salivary glands and Regional lymph nodes

Unremarkable submandibular, sublingual and parotid glands.
Regional lymph nodes display typical nodal histology.
(91544)
No lesions of significance

Thyroids

Portion of thyroid gland with typical colloid secreting follicles lined by cuboidal epithelium.
Includes small sheet-like mass of polygonal cells, characteristic of the parathyroid gland.
(91546)
No lesions of significance

Trachea/Lungs

Typical lung parenchyma/alveoli, bronchioles, and blood vessels.
Various degrees of parenchymal congestion and collapse (atelectasis), judged to be artefactual.
Portion of trachea with unremarkable mucosal epithelial lining and hyaline cartilage.
Sections do not include oesophagus.

(91546)
No lesions of significance

Thymus

Typical medulla/cortex distribution and micromorphology.
(91546)
No lesions of significance

*Comments:*

*Pathology to comment*

Heart/chambers/vessels/valves

Representative cardiac muscle, chambers, valves and great vessels of the heart.
Myxomatous valvular changes (thickened leaflet), an incidental finding in mice – feature seen in the controls.
The cardiac muscle fibres demonstrated typical features including central nuclei, branching fibres and striations.

Query severe focal inflammation and necrosis of the vascular/atrial(?) wall.

(91546)

*Comments:*

*Pathology to comment*

Skin

Mostly typical sections through the skin with readily discernible dermal appendages and distribution.
Mild to moderate multifocal subcutaneous inflammation with mainly mononuclear cell infiltrates, some hyperplasia of the epidermis, and myocyte degeneration of the panniculus muscle. An incidental finding in mice.
(91547)

Tail

Typical tail components including keratinized squamous epithelium, dense regular connective tissue, tendons, caudal vertebra, bone marrow, intervertebral disc, skeletal muscle, nerves and blood vessels.
(91548)
No lesions of significance

Eyes/Harderian glands

Unremarkable retina, cornea, iris, ciliary body, lens, sclera and choroid.
Typical branched tubuloalveolar formation of the Harderian gland.
Includes portion of unremarkable optic nerve and extraocular muscles.
(91549)
No lesions of significance

Brain

Sections were prepared from the standard levels of the brain:

Level I: including cortex (or cortical canopy), corpus callosum, caudate putamen and lateral ventricles (approx. Bregma -0.10mm)
Level II: including the hippocampus, thalamus, hypothalamus and lateral and third ventricles (approx. Bregma -1.70mm)
Level III: includes the cerebellum, pons and fourth ventricle (approx. Bregma -5.52mm)

Sections of brain appear symmetrical with no ventricular dilation observed, unremarkable meninges and typical lamination.

The cerebellum appears symmetrical with typical architecture and Purkinje cells.
There was no obvious neuronal loss, and the myelination appears normal.
(91550)
No lesions of significance

*Comments:*

*Neuropathology to comment*

Spinal cord

Representative thoracic and lumbar region of spinal cord, vertebral bone, intervertebral disc, striated muscle, peripheral nerves, brown adipose tissue, and bone marrow.
(91551, 91552)
No lesions of significance

*Comments:*

*Neuropathology to comment*

(Hind leg) Long bone/Bone marrow/Synovial joint/Skeletal muscle

Representative long bone, bone marrow, striated muscle, synovial joint, portion of tarsal bones, phalanges, cornified foot pad with discernible eccrine glands.
The skeletal muscle shows consistent fibre size with peripheral nuclei.

Subcutaneous oedematous changes in the distal hind leg, an incidental finding in mice (possibly due to handling).
Focal cortical bone changes (femur) near the knee joint – feature seen in the controls.

Query inflammatory changes near the knee joint.

(91553, 91554)

*Comments:*

*Pathology to comment*

Head

Multiple levels through the head demonstrate dermal appendages, nasal cavity with unremarkable nasal epithelium, oral cavity, dentition, and tongue with keratinized papillae.
Sections also show unremarkable pituitary gland including small portion of pars intermedia, pars distalis and pars nervosa as well and the trigeminal nerve/ganglia (91555).
The outer and middle regions of the ear are discernible. The tympanic membrane is intact, and the ossicles are unremarkable and include the stapedial annular ligaments.
Typical components of the inner ear including bony labyrinth, organ of corti, stria vascularis and scala cavities are discernible. Based on multiple levels, the organ of corti is unremarkable with no discernible loss of inner/outer hair cells and typical tectorial membrane.
The cochlear nerve and spiral ganglion is also demonstrated and based on several levels, there is no reduction in the density of the spiral ganglion cells. Examples of otolith organs can be seen with typical features such as the hair cells and mineral otoliths. The ampulla including the crista ridge with hair cells is discernible (91556).

(91555, 91556)
No lesions of significance

Sternum

Representative sternebrae, costal cartilage, intersternebral joint, intercostal skeletal muscle and brown fat.
Sections show haematopoietic tissue islands and adipose cells surrounded by vascular sinuses interspersed within a meshwork of trabecular bone. The bone marrow morphology demonstrates typical myeloid features including conspicuous megakaryoblasts and lymphoid features.
(91557)
No lesions of significance

#### Comment / Plan

Case APN22/057SVI (C. Walkley) will be referred to (text removed CW190123) for neuropathology comment.

(text removed CW190123)
25th October, 2022

______________________

Representative slides for case APN22/057SVI (C. Walkley) will be referred to (text removed CW190123), Specialist Veterinary Pathologist for comment.

(text removed CW190123)
28th October, 2022

### Supplementary Pathology Report

#### 154 (control)

Small intestine: proliferative and inflammatory enteritis.

Large intestine: mild colitis.

Hind leg: increased osteolysis and remodelling, consider stifle joint instability as a result of anatomical anomaly perhaps.

Spleen: no abnormalities detected.

Thymus: no abnormalities detected.

#### 491 (control)

Small intestine: proliferative and inflammatory enteritis. The queried cellular structures are Peyer's patches.

Hind leg: increased osteolytic activity at the neck of the femur, remodelling, joint instability a possible cause? The tibia looks bowed cranially?

Spleen: no abnormalities detected.

Thymus: no abnormalities detected.

#### 499 (control)

Small intestine: mild goblet cell metaplasia.

Heart: mild focal myocarditis.

Spleen: no abnormalities detected.

Thymus: no abnormalities detected.

## 151

Peripheral blood smear: difficult to interpret these blood films because they are too thick without a decent feathered edge and monolayer, and the background very blue (high protein?). The poikilocytes seen in this smear could be artefactual.

Small intestine: proliferative and inflammatory enteritis.

Spleen: extramedullary haematopoiesis, diffuse mild to moderate. Moderate diffuse reactive lymphoid hyperplasia.

Thymus: typical medulla/cortex distribution and micromorphology.

## 155

Peripheral blood smear: I am not comfortable calling these changes (poikilocytes and microcytes) on this smear.

Uterus/cervix/vagina: mild to moderate neutrophilic endometritis/vaginitis. No visible bacteria on H&E. Stage of cycle oestrus.

Small intestine: proliferative and inflammatory enteritis.

Spleen: extramedullary haematopoiesis, diffuse mild to moderate. Moderate diffuse reactive lymphoid hyperplasia.

Pancreas: "irregular focus" is “accessory splenic tissue” in the pancreas or adjacent to it, incidental.

Thymus: typical medulla/cortex distribution and micromorphology.

## 167

Peripheral blood smear: smear is too thick, but there is minimal anisocytosis with scanty poikilocytes and rare microcytes.

Small intestine: proliferative and inflammatory enteritis.

Spleen: extramedullary haematopoiesis, diffuse mild to moderate. Moderate diffuse reactive lymphoid hyperplasia.

Thymus: typical medulla/cortex distribution and micromorphology.

Sternum: multifocal myositis, mild.

## 492

Peripheral blood smear: smear is too thick, but there is minimal anisocytosis with scanty poikilocytes and rare microcytes. Rare band neutrophils.

Small intestine: proliferative and inflammatory enteritis.

Spleen: extramedullary haematopoiesis, diffuse moderate to severe. Moderate diffuse reactive lymphoid hyperplasia.

Thymus: typical medulla/cortex distribution and micromorphology.

Heart: queried vascular changes; possible plane of section artefact.

## 500

Peripheral blood smear: smear is too thick, but there is minimal anisocytosis with scanty poikilocytes and rare microcytes. Rare band neutrophils.

Small Intestine: proliferative and inflammatory enteritis.

Spleen: extramedullary haematopoiesis, diffuse mild to moderate. Moderate diffuse reactive lymphoid hyperplasia.

Thymus: typical medulla/cortex distribution and micromorphology.

Hind leg: increased osteolysis and remodelling, consider stifle joint instability because of anatomical anomaly perhaps.

## 513

Peripheral blood smear: smear is too thick, but there is minimal anisocytosis with scanty poikilocytes and rare microcytes.

Small Intestine: proliferative and inflammatory enteritis.

Spleen: increased extramedullary haematopoiesis, and atrophy of the lymphoid tissue.

Thymus: typical medulla/cortex distribution and micromorphology.

Heart: severe locally extensive haemorrhage, necrosis, and mononuclear inflammation of the mediastinal tissues at the heart base, extending into the right atrial wall. Mononuclear inflammation is also present on the pericardial surface. Did this animal have oral gavage or cardiac bleed? At any stage? This could be a traumatic lesion. Alternatively, a spontaneous haemorrhage with inflammation?

Hind leg: reactive change in the periosteum of the proximal tibia, increased osteolysis/remodelling of the femur near the joint, apparent hypercellularity of tendons to the tibia-kneecap. Again query joint instability, or anatomical anomaly?

#### Summary

Although there is some degree of proliferative and inflammatory enteritis in all animals, the control animals (154, 491, 499) did not show an increase in intestinal mucosal apoptosis. Apoptosis occurs as part of the normal intestinal homeostasis and renewal; however excessive apoptosis can result in pathology. Increased apoptosis is associated with many conditions including nutrient deficiency, endoplasmic reticulum stress, growth factor withdrawal, heat shock, developmental anomalies, and damage to or ligation of death-receptors on the cell surface e.g. Fas ligand.

The bone changes could be secondary to joint instability; not identified on the sections examined. The pathology appears to be remodelling of bone. The splenic reactive hyperplasia may suggest the inflammation in the intestines, or be secondary to other antigenic stimulation, e.g. infectious agents or parasites (not observed in the sections examined, apart from intestinal trichomonads). There are abundant Trichomonad protozoa in the caecum and colon, these have been associated with inflammation in the intestines. The variety of heart changes are interesting, but I cannot explain them. Blood films should have a large monolayer for erythrocyte and leucocyte morphology to be evaluated.

(text removed CW190123)
Tuesday 8th November, 2022, 8:45am APN22/057
Specialist Veterinary Pathologist
08/11/2022

#### 154 (control)

91394 – Brain – NAD.

91396 – Spinal cord – NAD.

#### 491 (control)

91461 – Brain – NAD.

91463 – Spinal cord – NAD.

#### 499 (control)

91505 – Brain – NAD.

91507 – Spinal cord – NAD.

## 151

91372, 91373 – Brain – Query hydrocephalus. Lateral ventricles appear dilated (but they often do depending on plane of section) and periventricular spongy change (but this is common in immersion-fixed brains). Difficult to determine.

91375 – Spinal cord – NAD.

## 155

91416 – Brain – NAD.

91418 – Spinal cord – NAD.

## 167

91439 – Brain – NAD.

91441 – Spinal cord – NAD.

## 492

91482 – Brain – NAD.

91484 – Spinal cord – NAD.

## 500

91526 – Brain – NAD.

91528 – Spinal cord – NAD.

## 513

91550 – Brain – NAD.

91552 – Spinal cord – NAD.

#### Summary

Sections of the brain and spinal cord show no significant findings.
Possible hydrocephalus in #151 but could be artefact.

NAD = No abnormalities detected.

(text removed CW190123)
Senior Veterinary Pathologist

28th October, 2022

Phenomics Australia advises all research groups that images or results obtained are to be acknowledged in resultant publications. Example acknowledgement: "This study utilised Phenomics Australia Histopathology and Slide Scanning Service, University of Melbourne".

### APN22/057SVI
### Macro Images

Carl Walkley

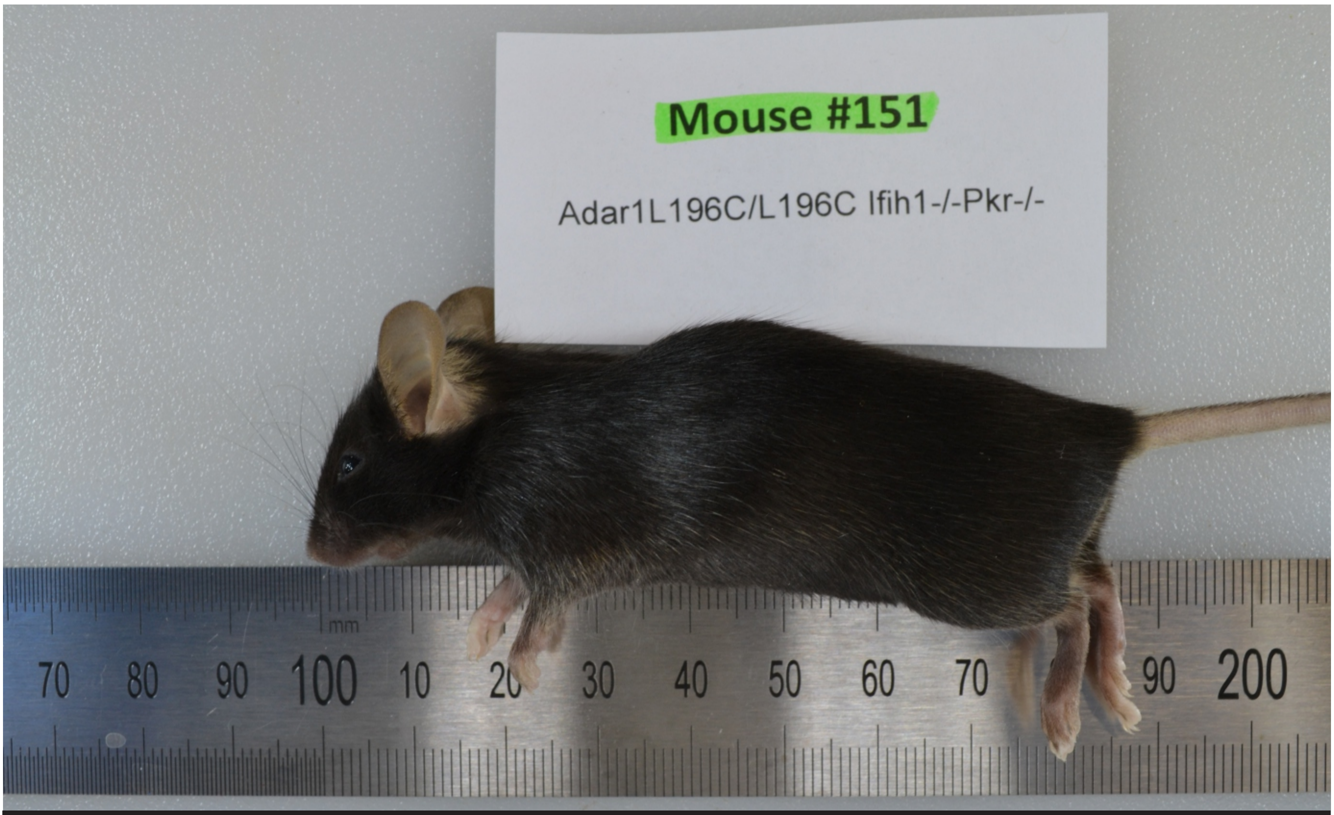


**#151**
**Adar1L196C/L196C Ifih1-/- Pkr-/- Female 169/22**

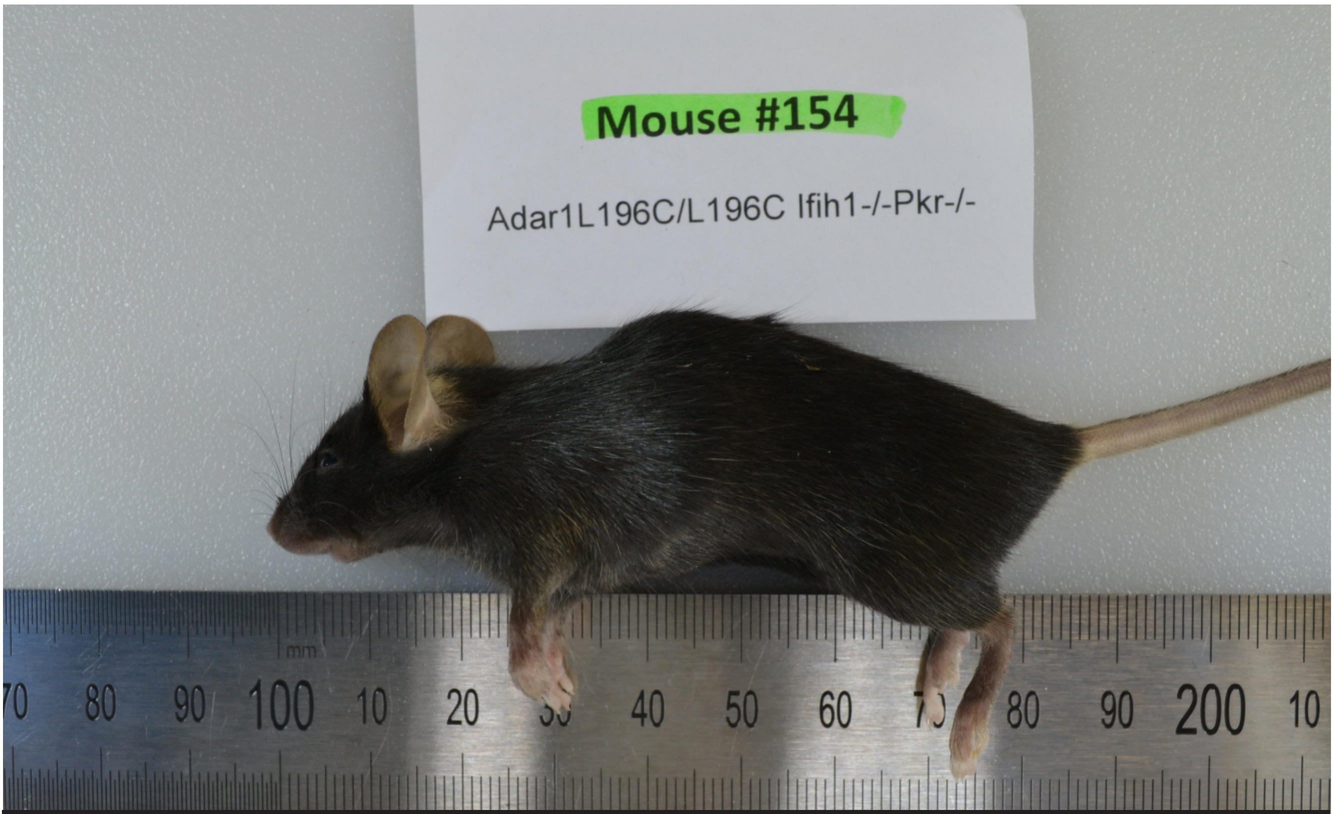


**#154**
**Adar1L196C/L196C Ifih1-/- Pkr-/- (control) Female 171/22**

### APN22/057SVI
### Macro Images

Carl Walkley

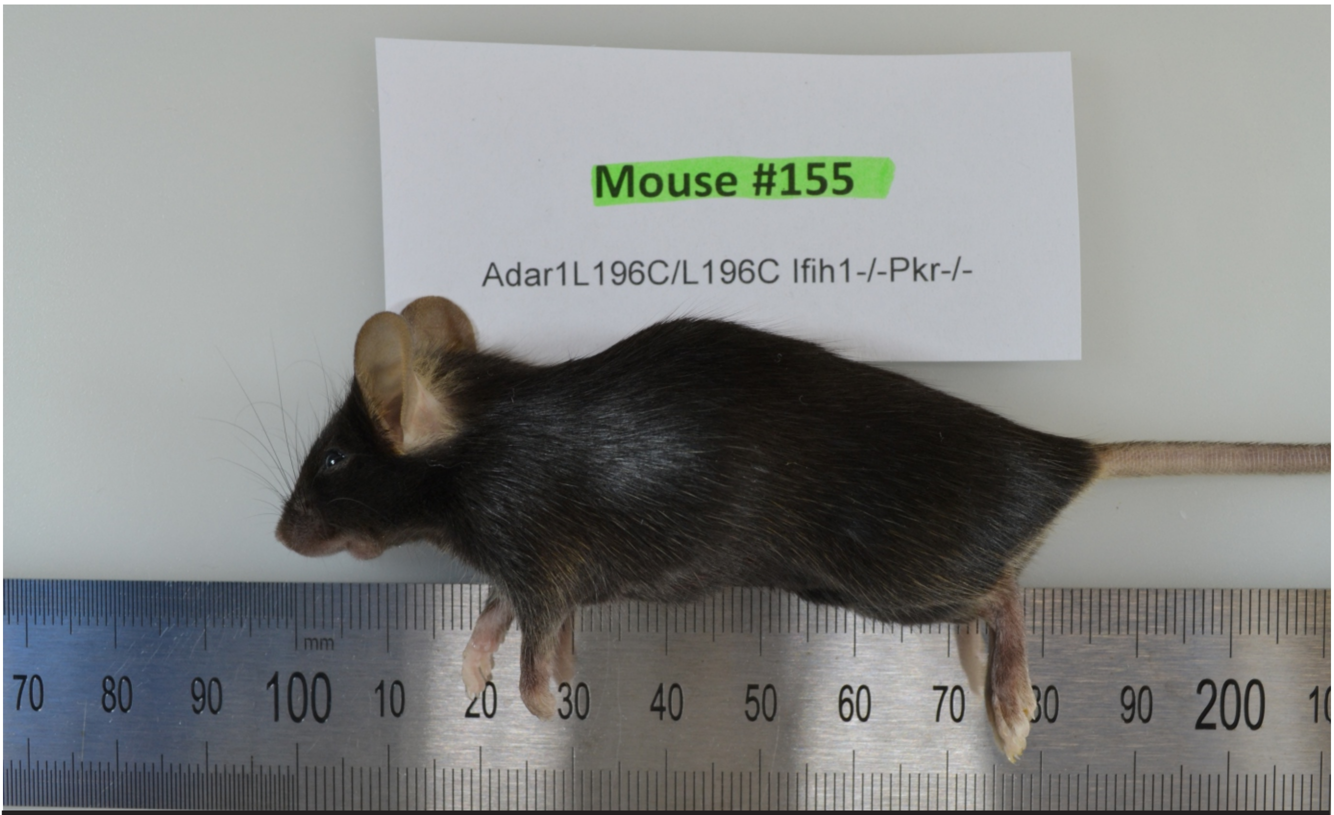


#155
Adar1L196C/L196C Ifih1-/- Pkr-/- Female 171/22

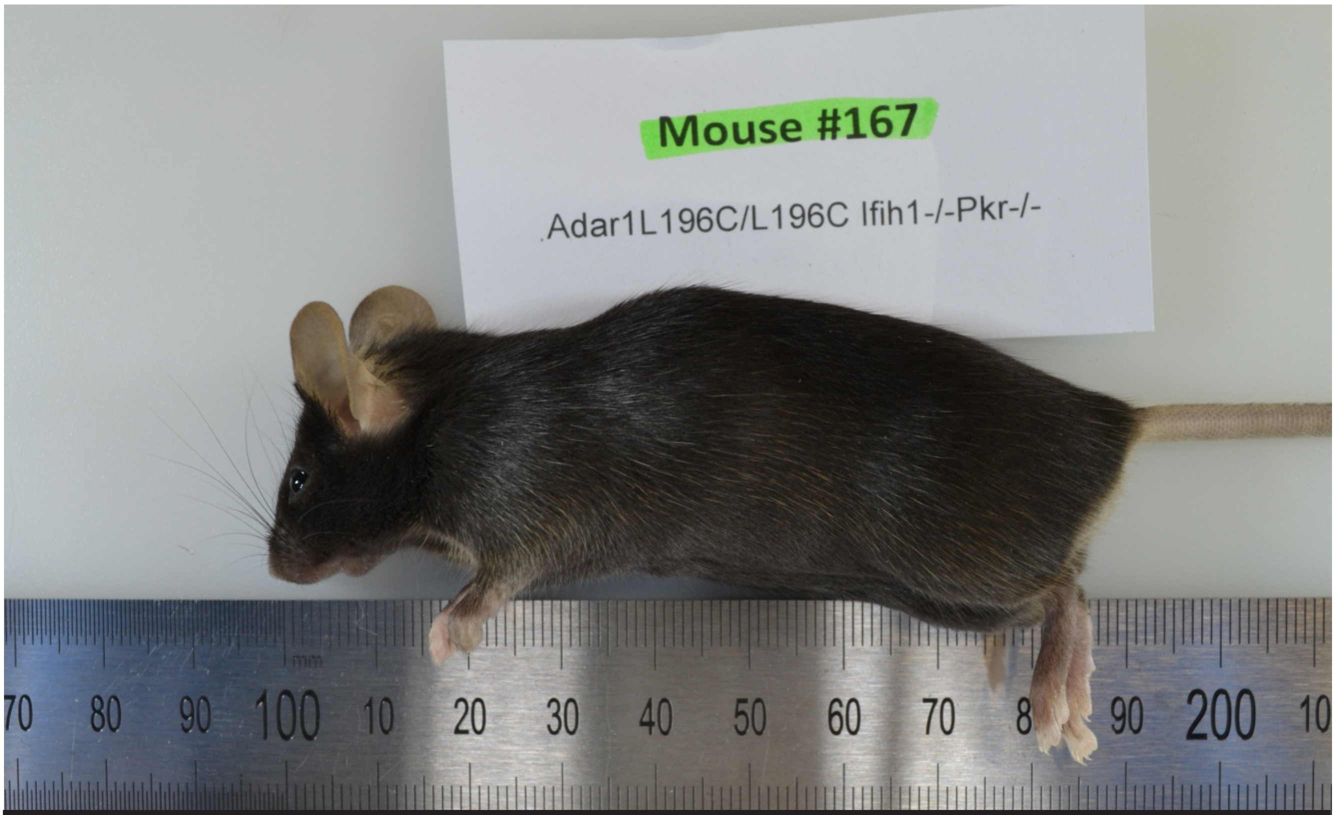


#167
Adar1L196C/L196C Ifih1-/- Pkr-/- Male 197/22

### APN22/057SVI
### Macro Images

Carl Walkley

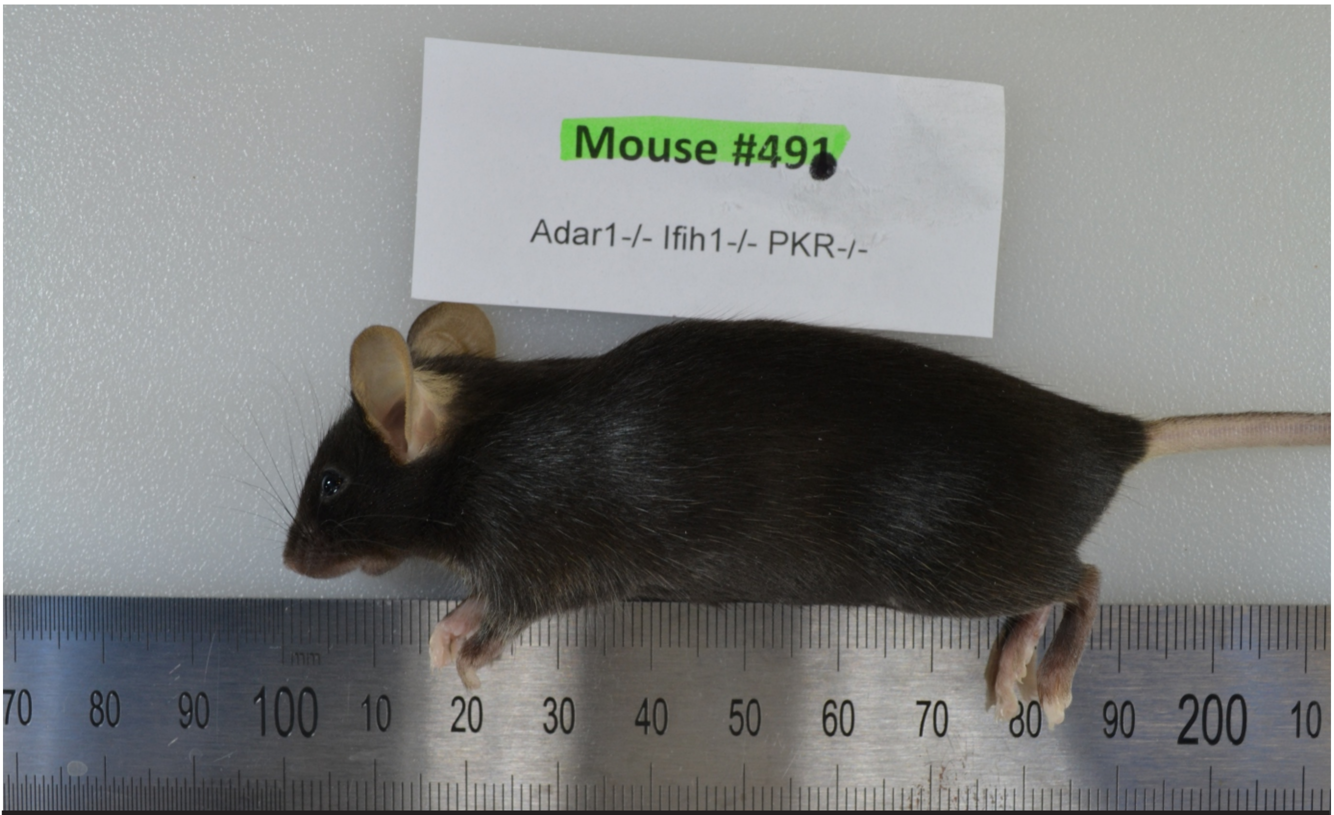


#491
Adar1-/- Ifih1-/- PKR-/- (control) Female 154/22

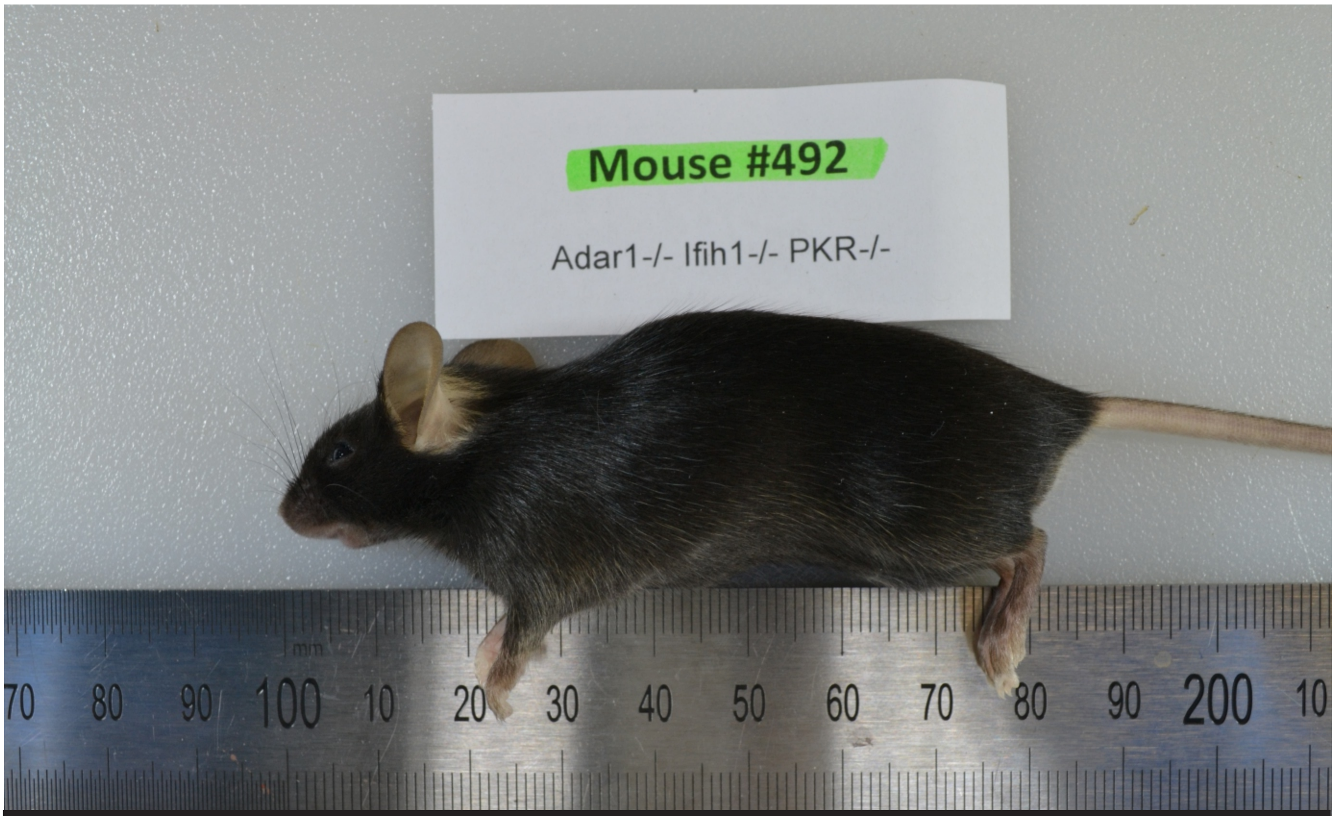


#492
Adar1-/- Ifih1-/- PKR-/- Female 154/22

### APN22/057SVI
### Macro Images

Carl Walkley

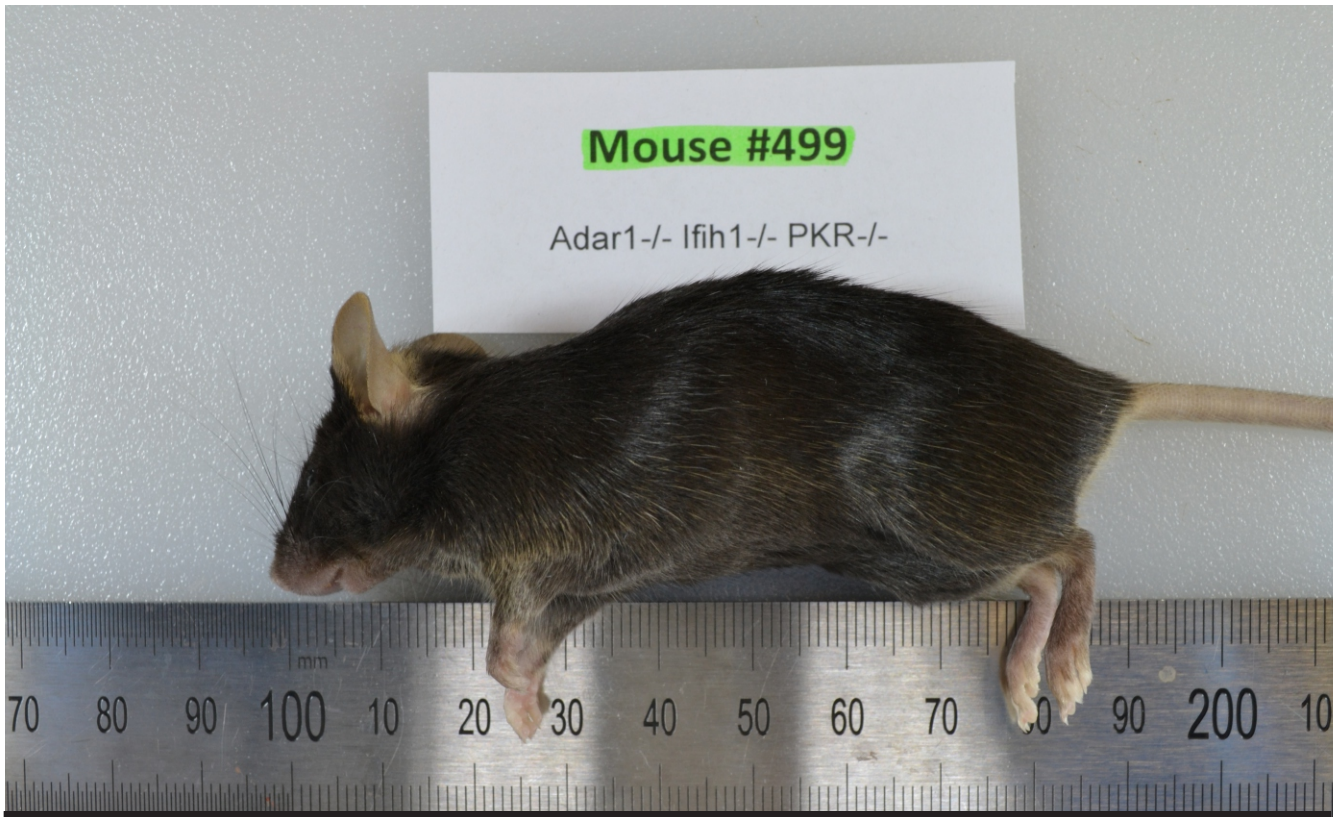


**#499**
**Adar1-/- Ifih1-/- PKR-/- (control) Male 161/22**

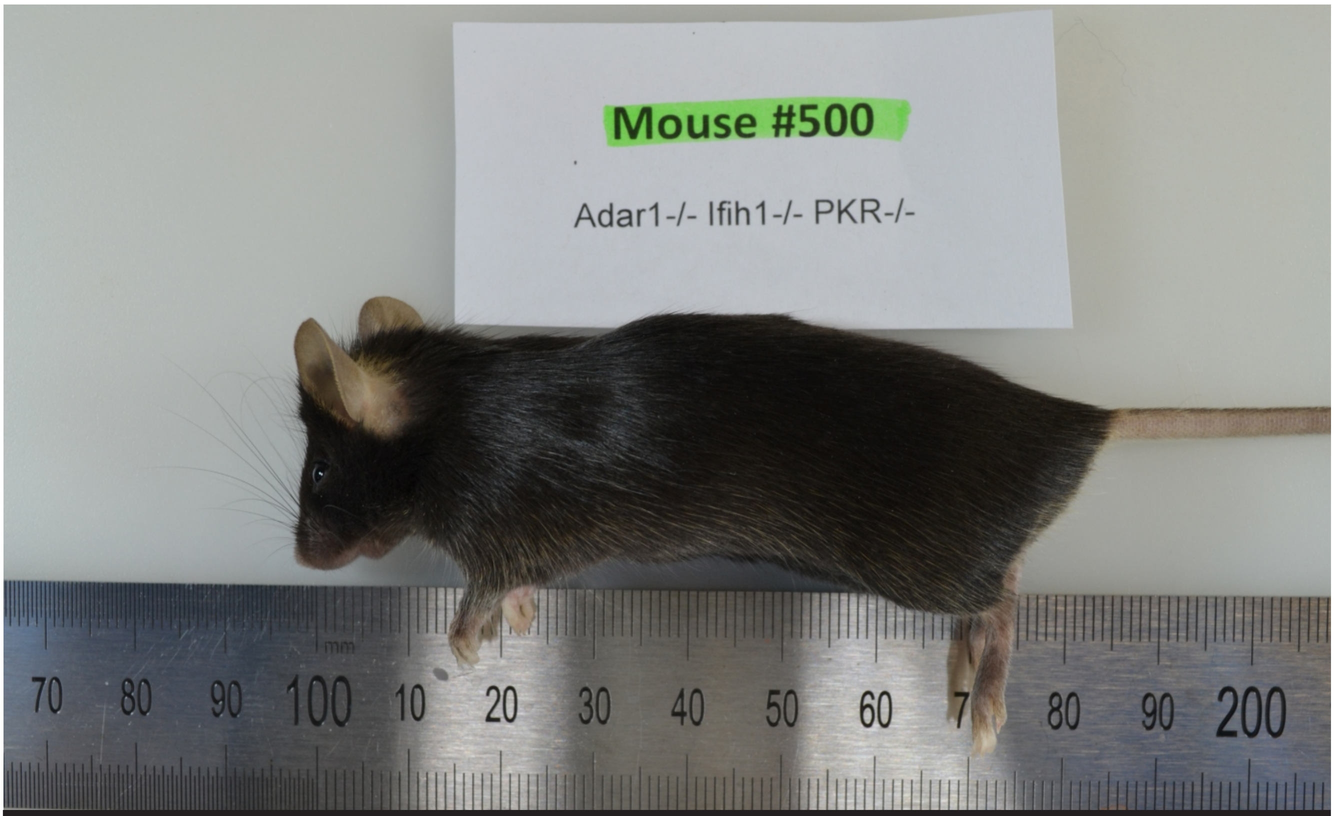


**#500**
**Adar1-/- Ifih1-/- PKR-/- Male 161/22**

### APN22/057SVI
### Macro Images
Carl Walkley

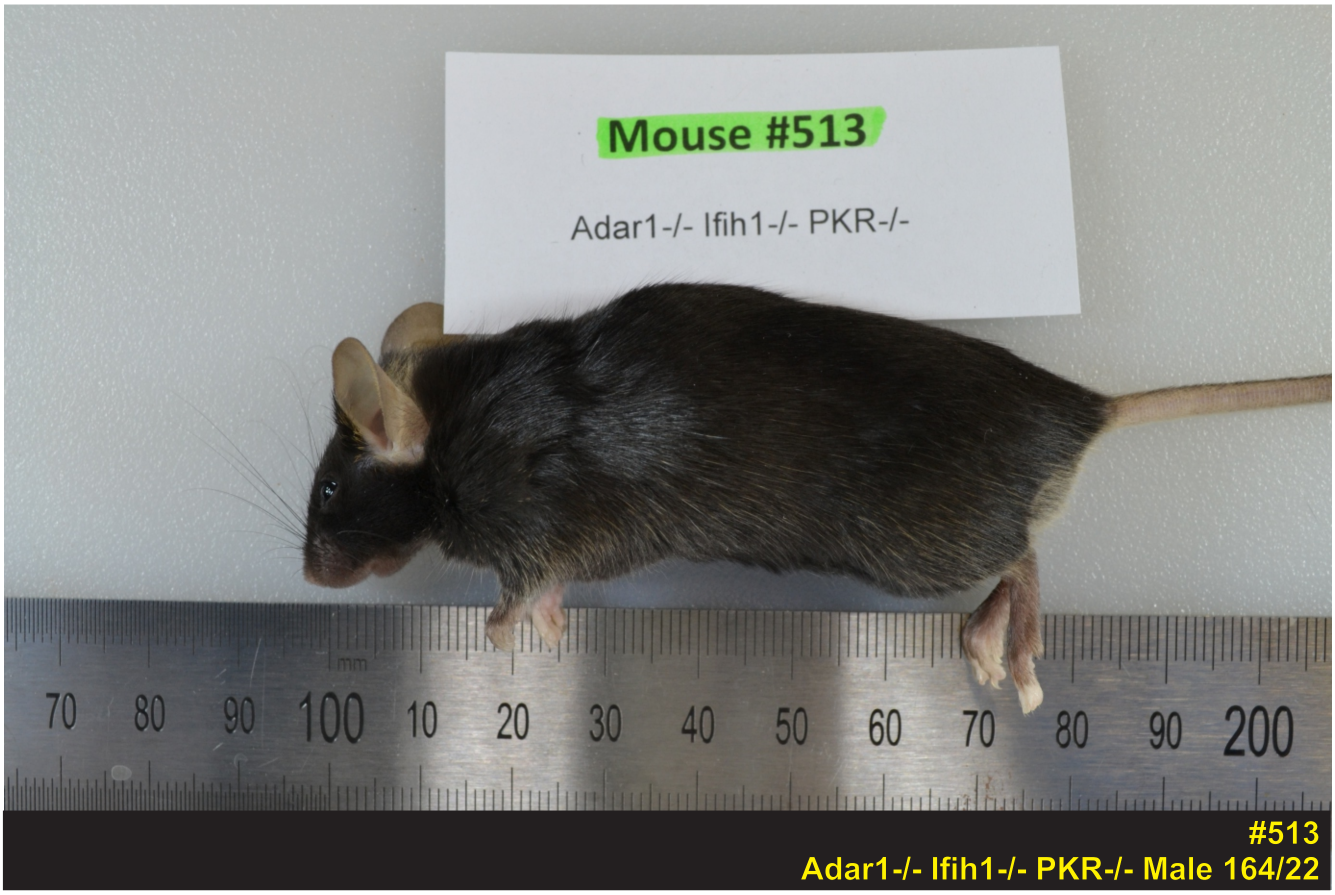

### APN22/057SVI
### Micro Images

Carl Walkley

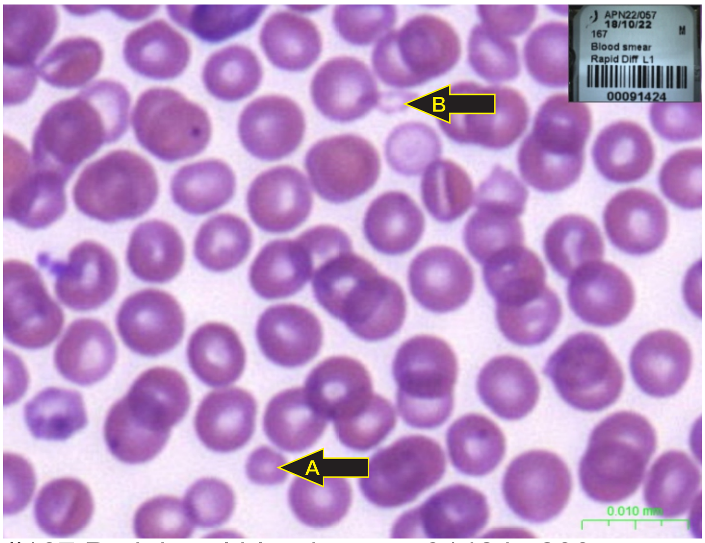


#167 Peripheral blood smear 91424 x200
Red blood cells - Microcyte (A), poikilocyte (B)

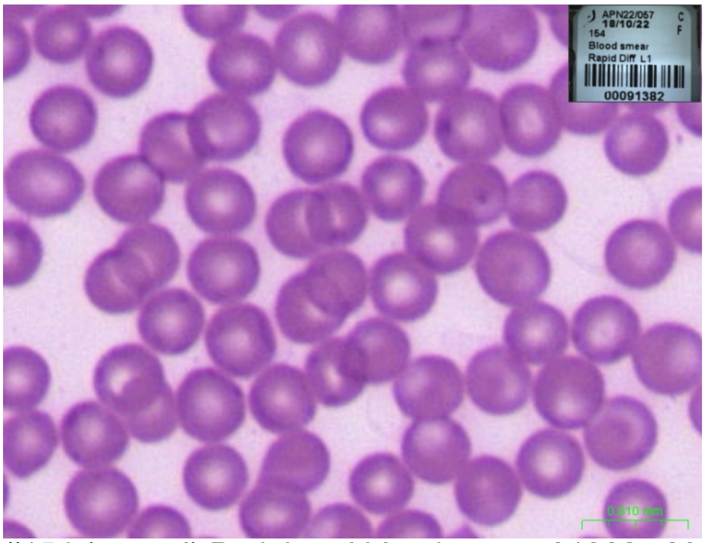

#154 (control) Peripheral blood smear 91382 x200
Red blood cells - for comparison

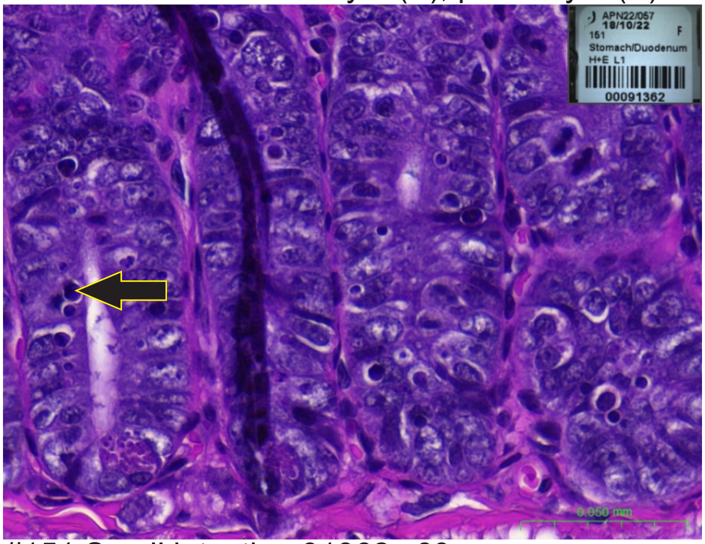

#151 Small intestine 91362 x63
Duodenum, crypts - apoptosis

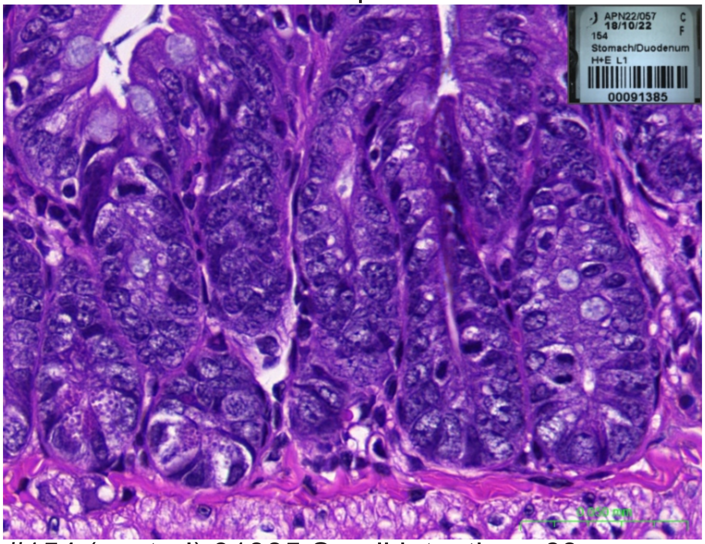

#154 (control) 91385 Small intestine x63
Duodenum, crypts - for comparison

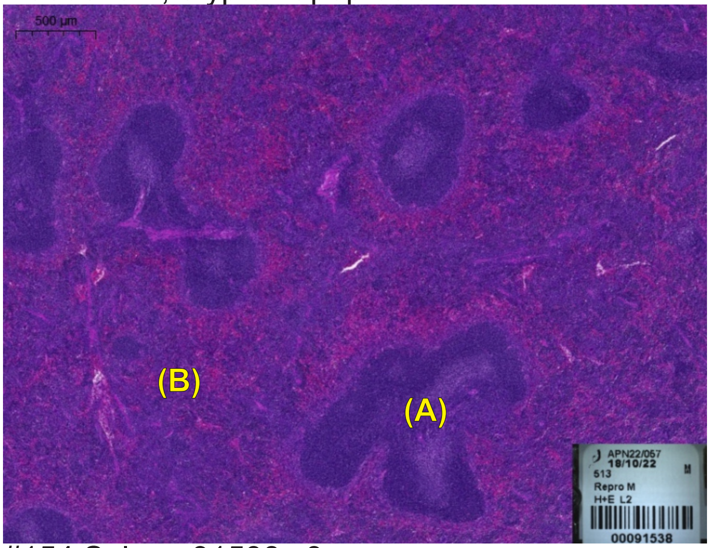


#154 Spleen 91538 x3
Decreased white pulp (A), increased extramedullary haematopoiesis (B)

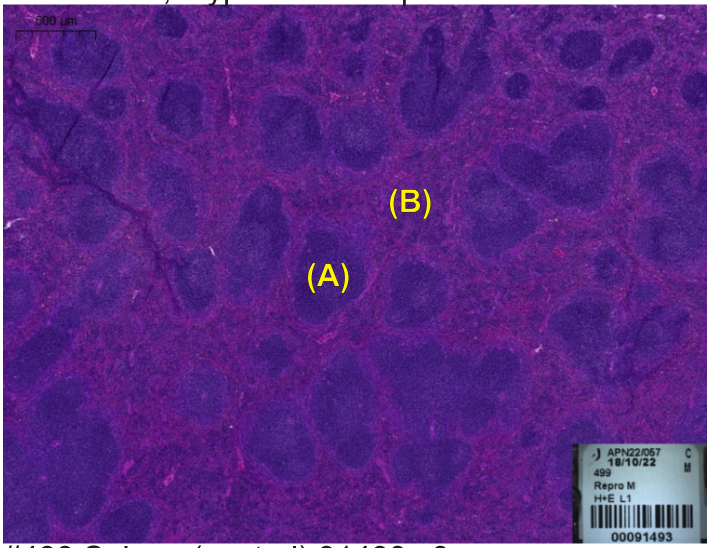


#499 Spleen (control) 91493 x3
White pulp (A), Red pulp (B) - for comparison
